# Engineered DNA materials shape microbial physiology and ecological selection

**DOI:** 10.64898/2026.11.06.764811

**Authors:** Laura Meisch, Marta Velaz Martín, Svenja A. Moench, Simone Weigel, John Vollmers, Paula S. Jäger, Anne-Kristin Kaster, Kersten S. Rabe, Christof M. Niemeyer

**Author notes:** Corresponding Author*, **Christof M. Niemeyer** - Institute for Biological Interfaces 1 (IBG-1), Biomolecular Micro- and Nanostructures, Karlsruhe Institute of Technology (KIT), D-76344 Eggenstein-Leopoldshafen, Germany.

## Abstract

Extracellular DNA (eDNA) is both a structural component of microbial biofilms and a potential reservoir of genetic information, yet its role as an ecological material remains largely unexplored. Here we establish engineered extracellular materials as determinants of microbial physiology and community assembly across increasing biological complexity. A three-dimensional DNA hydrogel altered bacterial stress-associated responses, elicited species-dependent phenotypes and transcriptional changes in a defined consortium, and shaped community succession in wastewater-derived microbiomes. Distinct responses to surface-bound oligonucleotides reveal that DNA presentation matters beyond its mere presence. Strikingly, metagenomic sequencing identified rare, hydrogel-specific candidate junctions linking material-derived DNA to microbial genomic sequences, suggesting a previously unexplored interface between extracellular DNA materials and microbial genetic information. These findings reveal a dual role for engineered DNA: its material organization shapes microbial physiology and ecological selection, while its encoded sequences may engage with microbial genomic processes.

## Introduction

Microbial communities inhabit virtually every natural and engineered environment and contribute fundamentally to global biogeochemical cycles, environmental processes and host-associated ecosystems. Their phylogenetic and functional diversity offers considerable biotechnological potential, yet many environmental microorganisms remain difficult to cultivate under conventional laboratory conditions, contributing to the vast reservoir of microbial dark matter (MDM).^[1–3]^ In situ cultivation (ISC) approaches seek to access this diversity by enabling microbial colonization within devices that maintain exchange with the surrounding environment.^[4–6]^ Beyond improving microbial recovery, these approaches raise a fundamental question: to what extent do the materials used for cultivation themselves influence microbial selection and the resulting community composition?

Material surface chemistry and physicochemical properties influence microbial attachment, biofilm formation and community composition.^[7–10]^ Recent investigations of microplastic-associated biofilms further illustrate how synthetic materials can establish distinct microbial habitats with consequences for microbial interactions and biofilm function.^[11]^ However, engineered interfaces need not be limited to conventional surface chemistries. Incorporating biologically active macromolecules may create extracellular environments that influence microbial physiology and community development beyond their effects on initial attachment. Such functional materials may therefore act as ecological filters rather than passive colonization substrates.

DNA is particularly attractive for engineering such environments because sequence-specific recognition and self-assembly enable programmable structures across multiple length scales. DNA hydrogels extend these principles to hydrated polymer networks with tunable structural and dynamic properties, offering opportunities in biomedical and biotechnological applications.^[12–18]^

DNA-based materials have also been interfaced with microorganisms, including approaches for selective bacterial capture,^[19]^ microbial biohybrid systems incorporating DNA-containing matrices,^[20]^ and bacterially produced components of engineered living materials.^[21]^ These advances demonstrate the technological versatility of DNA-based materials, but provide limited insight into how concentrated, three-dimensional extracellular DNA networks influence microbial physiology and multispecies community assembly.

This gap is especially relevant because extracellular DNA (eDNA) is a structural and functional component of natural microbial biofilms, contributing to adhesion and organization of the extracellular polymeric substance (EPS) matrix.^[22–25]^ The identification of supramolecular DNA structures, including G-quadruplex motifs associated with viscoelastic eDNA networks, further emphasizes that extracellular DNA can form organized material architectures rather than merely occur as dispersed nucleic acid.^[26]^ Yet the physiological and ecological consequences of exposing microorganisms to defined, DNA-rich extracellular materials remain insufficiently understood. Moreover, unlike most structural biomaterials, DNA also carries sequence information that could potentially interact with microbial genetic processes. Engineered DNA hydrogels offer an experimentally accessible way to investigate these material and informational dimensions without reproducing the compositional complexity of natural biofilm matrices.

Here, we adapted macroporous elastomeric silicone foam (MESIF), previously developed for microbial colonization and in situ cultivation,^[6]^ as a three-dimensional platform for systematically varying extracellular material environments (Fig. 1a). Seven conditions encompassed native PDMS, amino-, hydroxyl- and PEG-modified interfaces, immobilized catalase, surface-bound oligodeoxynucleotides (ODNs) and an extended DNA hydrogel generated by rolling-circle amplification (RCA) (Fig. 1b).^[27]^ The ODN and DNA hydrogel conditions contrasted DNA presentation without a hydrogel network with a highly concentrated, spatially organized DNA matrix, within a broader set of chemical and biomolecular environments.

**Fig 1.**
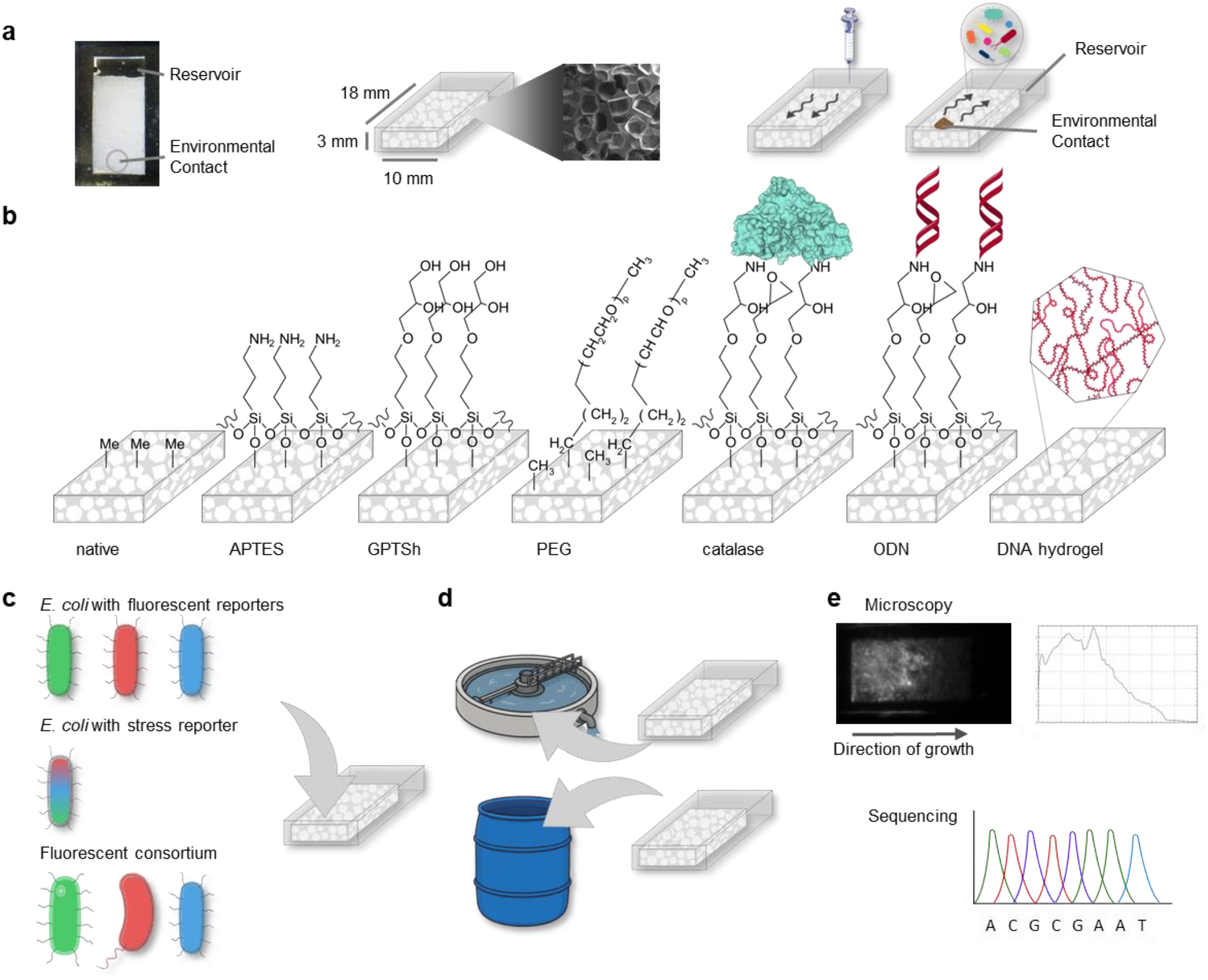
Experimental framework for investigating material-dependent microbial responses. **(a)** Schematic representation of the macroporous elastomeric silicone foam (MESIF) cultivation platform, comprising an interconnected porous scaffold integrated into a polydimethylsiloxane (PDMS) housing with a medium reservoir and an environmental contact region. Culture medium can be supplied through the reservoir, while the environmental contact enables exposure of the porous scaffold to surrounding microbial communities. **(b)** Seven material environments investigated in this study: native PDMS presenting methyl-terminated interfaces; APTES-functionalized surfaces bearing primary amino groups; hydrolysed GPTS (GPTSh) interfaces presenting hydroxyl groups; PEG-modified interfaces; immobilized catalase; surface-bound oligodeoxynucleotide (ODN) duplexes; and an extended DNA hydrogel generated by rolling-circle amplification (RCA) within the MESIF pore network. The representations are schematic and not drawn to scale. The DNA hydrogel is illustrated at the pore-network level to distinguish its extended polymeric architecture from the other, predominantly interface-associated functionalizations. **(c)** Experimental progression from fluorescently labelled *Escherichia coli* strains used to monitor growth-associated fluorescence and spatial colonization, through multiplexed RGB-S stress-reporter measurements in *E. coli*, to a fluorescently labelled three-species consortium comprising *Bacillus subtilis, Shewanella oneidensis* and *E. coli* for investigating species-specific responses within a shared material environment. **(d)** Exposure of functionalized MESIF to complex environmental microbial communities in a chemical wastewater treatment plant and a wastewater mesocosm to investigate material-associated community dynamics over different cultivation periods. **(e)** Fluorescence microscopy and RNA- and DNA-based sequencing approaches used to characterize microbial phenotypes, species-specific transcriptional states, community composition and DNA-associated molecular signatures.

We examined material-associated responses across increasing levels of biological complexity: growth-associated fluorescence and stress-reporter activity in *Escherichia coli*, species-specific fluorescence phenotypes and transcriptional states in a defined three-species consortium, and community succession in complex wastewater-derived microbiomes (Fig. 1c–e). We hypothesized that engineered extracellular materials can differentially shape microbial physiological states and community assembly, with DNA-rich environments providing a particularly informative test of how material architecture relates to biological responses. We further investigated whether hydrogel-derived DNA sequences could be detected in association with microbial genomic sequences, thereby probing a potential informational dimension of the material–microbe interface.

## Results and Discussion

### Engineering chemically and biomolecularly defined MESIF interfaces

To investigate how engineered extracellular materials shape microbial responses, we established seven distinct environments within a common macroporous elastomeric silicone foam (MESIF) platform (Fig. 1a,b). These comprised native polydimethylsiloxane (PDMS), amino-functionalized APTES, hydroxyl-functionalized GPTSh and PEG-modified interfaces, together with immobilized catalase, surface-bound oligodeoxynucleotides (ODNs) and an extended DNA hydrogel generated in situ by rolling-circle amplification (RCA). The physicochemical modifications followed established procedures,^[6,28]^ while RCA-based DNA material formation was adapted from our previous approach (Supplementary Fig. S1a–h).^[27]^

Modification-specific assays confirmed the establishment and functionality of the engineered environments (Supplementary Fig. S2a–h). APTES modified MESIF showed an approximately 25-fold increase in fluorescamine fluorescence relative to native PDMS, consistent with accessible primary amino groups (Supplementary Fig. S2a). SpyCatcher–SpyTag-mediated catalase immobilization yielded a loading of 29.2 ± 5.0 nmol per MESIF and retained hydrogen peroxide decomposition over prolonged incubation (Supplementary Fig. S2b–d). Surface attachment and hybridization of complementary ODN strands were supported by Cy3/Cy5 fluorescence measurements, including donor quenching and a FRET-associated hybridization signal (Supplementary Fig. S2e).

In contrast to these surface-bound DNA duplexes, the DNA hydrogel was generated within the interconnected MESIF pores using a 72-nucleotide circular RCA template containing three self-complementary domains (Supplementary Figs. S1h and S2f). After 72 h of polymerization and washing, fluorescent DNA material remained associated with MESIF, whereas polymerase-free controls showed minimal retained fluorescence (Supplementary Fig. S2g,h). Extracted DNA was quantified at 1.60 ± 0.23 µM. Together, these results establish a common porous platform presenting chemically and biomolecularly distinct environments, including two structurally different forms of extracellular DNA. Detailed fabrication and validation are provided in the Extended Description to Supplementary Figs. S1–S2. This platform enabled us to examine how material composition and DNA presentation relate to microbial phenotypes across increasing levels of biological complexity (Fig. 1c–e).

### Material interfaces differentially affect *E. coli* growth

To investigate how engineered material environments influence bacterial colonization, we cultivated *Escherichia coli* K-12 MG1655 strains expressing GFP, RFP or BFP within seven differently functionalized MESIF environments. Because fluorescent proteins differ in maturation, stability and cellular burden, the three strains were analyzed independently.^[29–31]^ In liquid culture, fluorescence correlated strongly with optical density for all three reporters (Spearman’s ρ = 0.94–0.96; Supplementary Fig. S3a–c), supporting its use as a growth-associated, rather than direct biomass, readout.

Longitudinal whole-MESIF imaging over 15 days revealed reporter- and material-dependent fluorescence trajectories (Supplementary Fig. S4a–i). GFP and RFP fluorescence generally persisted longer, whereas BFP signals declined more rapidly after their initial maximum (Supplementary Fig. S4d–f). Time-averaged measurements showed comparatively similar fluorescence across native, APTES-, GPTSh- and PEG-functionalized MESIF, while biomolecular environments produced more pronounced differences (Fig. 2a–c). Catalase-functionalized MESIF consistently exhibited low fluorescence, whereas ODN-functionalized MESIF maintained comparatively prominent signals. The DNA hydrogel showed a particularly divergent response: GFP-associated fluorescence remained relatively high, while RFP- and BFP-associated signals were comparatively weak (Fig. 2a–c; Supplementary Fig. S4g–i). Three-dimensional confocal imaging further revealed fluorescence-associated bacterial structures within individual MESIF pores, with material- and reporter-dependent spatial patterns (Fig. 2d).

**Fig 2.**
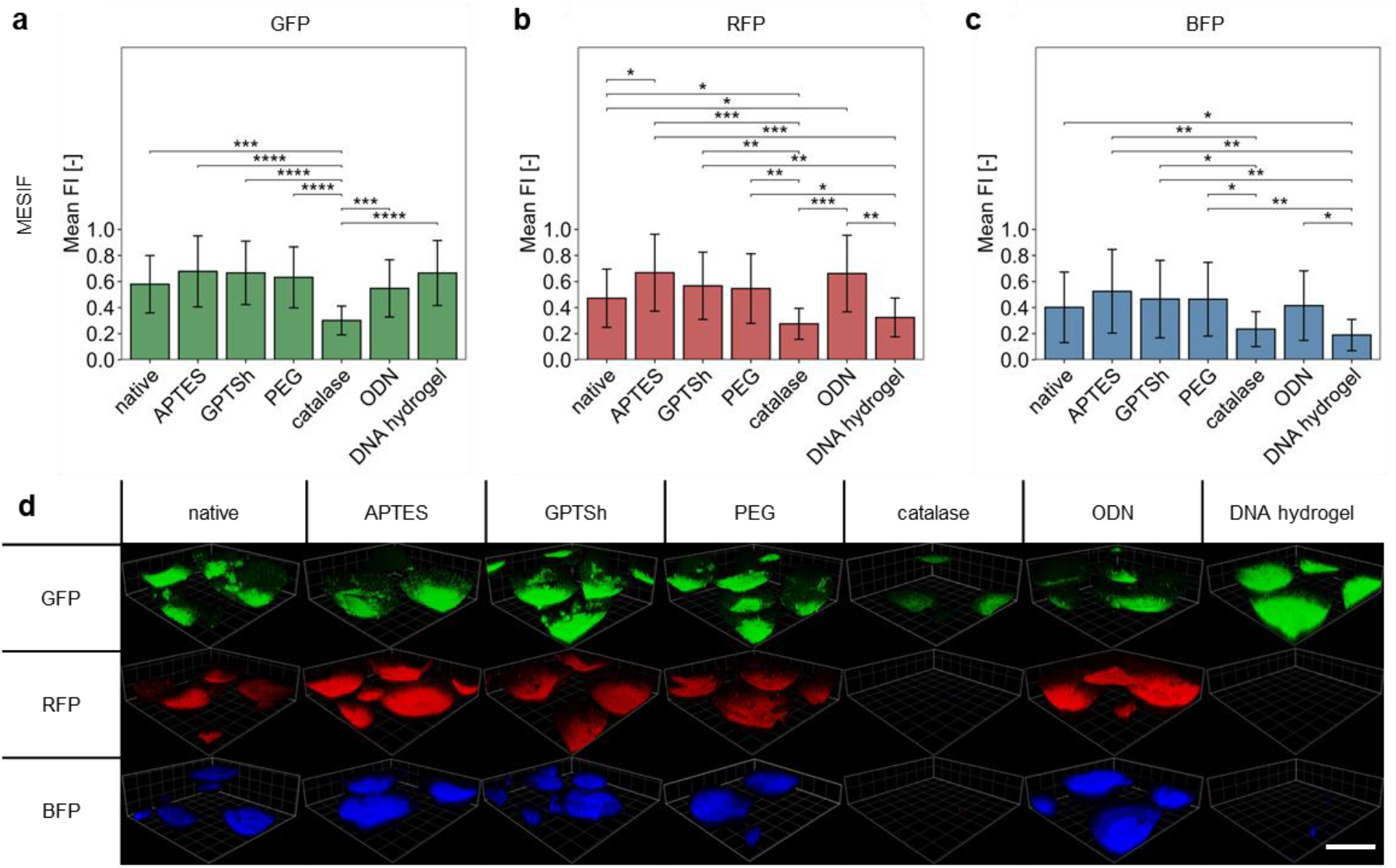
Material environments differentially affect growth-associated fluorescence of *E. coli*. *Escherichia coli* K-12 MG1655 strains expressing GFP, RFP or BFP were cultivated for 15 days in native, APTES-, GPTSh-, PEG-, catalase-, ODN- or DNA hydrogel-functionalized MESIF. **(a–c)** Time-averaged normalized whole-MESIF fluorescence intensities (FI) of GFP-expressing (a), RFP-expressing (b) and BFP-expressing (c) *E. coli* across the seven material environments. Bars indicate means and error bars represent standard deviations from n = 3 independent MESIF chips per material condition. Fluorescence intensities were normalized to the maximum value measured for the respective reporter strain. Statistical significance was assessed using linear mixed-effects models with material condition, cultivation time and their interaction as fixed effects and MESIF chip as a random effect. Pairwise comparisons between material conditions were performed using estimated marginal means with Benjamini–Hochberg correction for multiple testing. Significance levels are indicated as p < 0.05 (*), p < 0.01 (**), p < 0.001 (***), and p < 0.0001 (****). **(d)** Representative three-dimensional confocal laser scanning microscopy (CLSM) reconstructions of individual MESIF pores colonized by GFP-, RFP- or BFP-expressing *E. coli* under the indicated material conditions. Scale bar, 200 µm.

Together, these findings establish that extracellular material composition influences *E. coli*-associated fluorescence, but that the observed phenotypes depend substantially on the reporter used. Detailed analyses of reporter characteristics, temporal dynamics and spatial colonization are provided in the Extended Description to Supplementary Figs. S3–S4. The contrasting fluorescence profiles under DNA hydrogel conditions raised the question of whether this DNA-rich extracellular environment also induces distinct physiological responses beyond differences in growth-associated fluorescence. We therefore investigated material-dependent stress-associated transcriptional activity using a multiplexed reporter system.

### DNA hydrogel environments induce a distinct *E. coli* physiological state

To determine whether the material-dependent fluorescence phenotypes were accompanied by physiological changes, we employed the previously established RGB-S reporter system in *E. coli*.^[32]^ The reporter monitors SOS-associated genotoxic stress through PsulA–GFP, RpoS-mediated general stress through PosmY–RFP and RpoH-associated proteotoxic stress through PgrpE–BFP (Supplementary Fig. S5a,b).^[33–35]^

Longitudinal whole-MESIF imaging revealed material- and pathway-dependent fluorescence responses over 15 days (Fig. 3a–c; Supplementary Fig. S6a–f). DNA hydrogel-functionalized MESIF exhibited particularly pronounced SOS-associated fluorescence, elevated RpoS-associated signals and an early RpoH-associated response, whereas catalase-functionalized MESIF generally showed comparatively low absolute fluorescence (Fig. 3a–c; Supplementary Fig. S6d–f). Confocal imaging further revealed heterogeneous, temporally changing reporter fluorescence within individual pores (Fig. 3g; Supplementary Fig. S7a–c). Corresponding CLSM- and ROI-based whole-MESIF measurements showed broadly consistent material-associated patterns, although their agreement varied between reporter channels and time points (Supplementary Fig. S7d–g).

**Fig 3.**
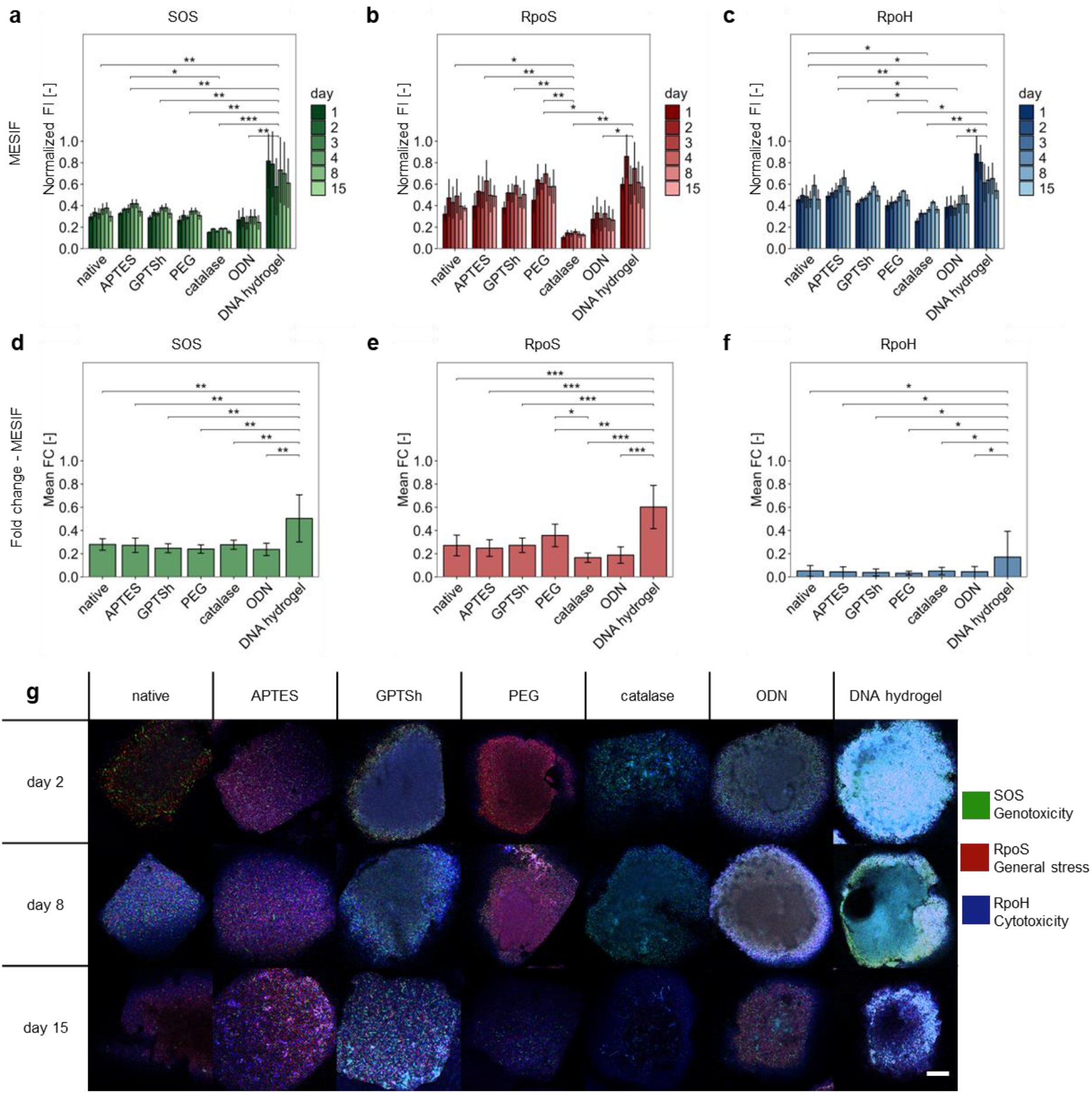
DNA hydrogel-functionalized MESIF elicits distinct stress-associated reporter responses in *E. coli*. *Escherichia coli* K-12 MG1655 carrying the RGB-S reporter plasmid was cultivated for 15 days in native, APTES-, GPTSh- , PEG-, catalase-, ODN- or DNA hydrogel-functionalized MESIF. The RGB-S reporter monitors SOS-associated genotoxic stress (GFP), RpoS-mediated general stress (RFP) and RpoH-associated proteotoxic stress (BFP). **(a–c)** Longitudinal whole-MESIF fluorescence intensities (FI) for the SOS (a), RpoS (b) and RpoH (c) reporter channels across the seven material environments. **(d–f)** Relative SOS (d), RpoS (e) and RpoH (f) reporter responses, expressed as fold changes (FC). For each cultivation day, material condition and reporter channel, the mean fluorescence intensity of the corresponding single-reporter strain was first calculated from the independent growth-associated fluorescence experiment. Individual RGB-S fluorescence intensities were then divided by this matched single-reporter mean, and the resulting FC values were subsequently averaged across the cultivation period for each material condition. Bars represent mean ± SD from n = 3 independent MESIF chips per material condition. Statistical significance was assessed using linear mixed-effects models with material condition, cultivation time and their interaction as fixed effects and MESIF chip as a random effect. Pairwise comparisons between material conditions were performed using estimated marginal means with Benjamini–Hochberg correction for multiple testing. Significance levels are indicated as p < 0.05 (*), p < 0.01 (**), p < 0.001 (***), and p < 0.0001 (****). **(g)** Representative confocal laser scanning microscopy images of individual MESIF pores showing merged SOS-associated (green), RpoS-associated (red) and RpoH-associated (blue) reporter fluorescence on days 2, 8 and 15 under the indicated material conditions. Scale bar, 100 µm.

Because absolute reporter fluorescence also depends on cell abundance and fluorescence-associated cellular state, we normalized each RGB-S channel to the corresponding single-reporter strain under matched material and cultivation-time conditions (Fig. 3d–f). DNA hydrogel-associated responses remained elevated across all three reporter pathways after normalization, indicating that the pronounced fluorescence phenotype could not be explained solely by differences in growth-associated reporter signals. Other materials showed more pathway-selective relative responses, including PEG-associated RpoS and catalase-associated RpoH signals (Fig. 3e,f).

Together, these findings identify the DNA hydrogel as a material environment associated with a distinctive, multi-pathway physiological reporter state in *E. coli*, rather than merely altered growth-associated fluorescence. Detailed interpretation of the reporter system, temporal profiles and spatial measurements is provided in the Extended Description to Supplementary Figs. S5–S7. We next investigated whether such material-associated responses differ among species cultivated within a shared microbial consortium.

### Material environments reshape species-specific performance in a microbial consortium

To determine whether material-dependent responses observed in individual *E. coli* populations extend to a multispecies setting, we established a defined consortium comprising fluorescently labelled *E. coli, Bacillus subtilis* and *Shewanella oneidensis*. Reporter fluorescence was characterized in liquid culture before being used as a species-specific, growth-associated readout in MESIF (Supplementary Fig. S8a–e). The organisms were cultivated individually or as an initially equal-ratio consortium across seven material environments, with longitudinal whole-MESIF fluorescence measurements normalized to the species-specific maximum observed in monoculture (Fig. 4a–f and Supplementary Fig. S9). In monoculture, all three species exhibited distinct material-associated fluorescence profiles, with comparatively low signals frequently observed in catalase-functionalized MESIF (Fig. 4a–c). Notably, *E. coli* showed reduced fluorescence in DNA hydrogel-functionalized MESIF, consistent with the phenotype observed in the preceding independent *E. coli* experiments (Fig. 4c and Supplementary Fig. S9c,f).

**Fig 4.**
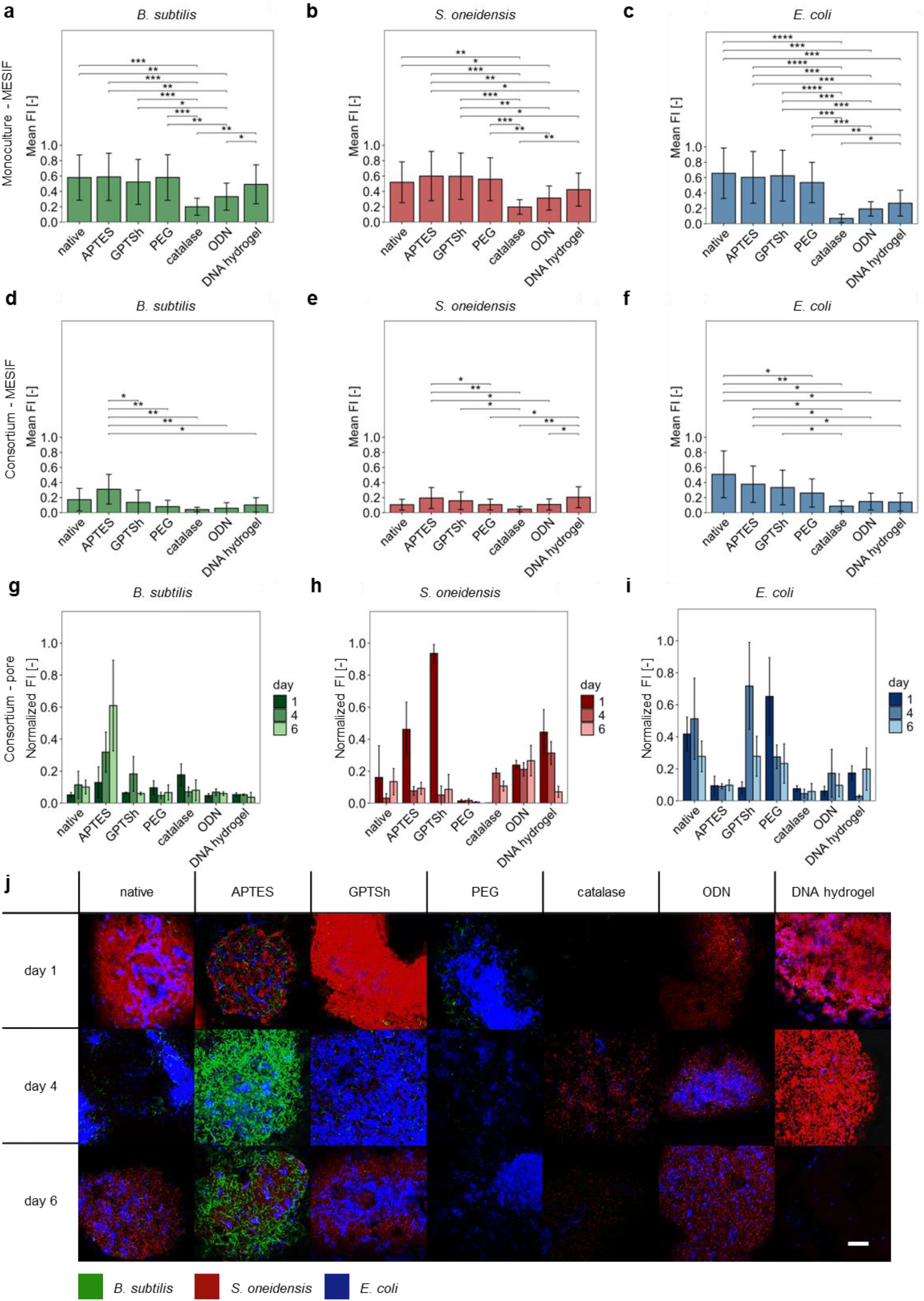
Material environments reshape species-specific fluorescence phenotypes and spatial organization in a three-species microbial consortium. GFP-expressing *Bacillus subtilis* 1012, RFP-expressing *Shewanella oneidensis* MR-1 and BFP-expressing *Escherichia coli* K-12 MG1655 were cultivated individually or as an initially equal-ratio three-species consortium in MESIF presenting native, APTES-, GPTSh-, PEG-, catalase-, ODN- or DNA hydrogel-functionalized material environments. **(a–c)** Species-specific fluorescence responses of *B. subtilis* (a), *S. oneidensis* (b) and *E. coli* (c) during monoculture cultivation. **(d–f)** Corresponding fluorescence responses of *B. subtilis* (d), *S. oneidensis* (e) and *E. coli* (f) during consortium cultivation. For each species, fluorescence measurements were normalized by dividing each value by the maximum fluorescence intensity observed for that species under monoculture conditions. The resulting values were averaged across all analysed time points to obtain the time-averaged normalized fluorescence intensities shown in a–f. Normalization permits comparisons of material-associated fluorescence responses within each species but does not provide a quantitative measure of relative abundance between species. Data are presented as mean ± standard deviation from three independent MESIF chips per material environment and culture condition. Longitudinal fluorescence trajectories were analysed separately for monoculture and consortium cultivation using linear mixed-effects models with material environment, cultivation time and their interaction as fixed effects and MESIF chip as a random effect. Pairwise comparisons between material environments within each species and cultivation context were performed using estimated marginal means with Benjamini–Hochberg correction for multiple testing. Asterisks indicate adjusted *p* values of p < 0.05 (*), p < 0.01 (**), p < 0.001 (***), and p < 0.0001 (****). **(g–i)** Time-resolved confocal laser scanning microscopy (CLSM) measurements of fluorescence intensity within individual MESIF pores for *B. subtilis* (g), *S. oneidensis* (h) and *E. coli* (i) during consortium cultivation on days 1, 4 and 6. Bars represent mean fluorescence intensity ± standard deviation across multiple CLSM images acquired from a representative MESIF chip per condition. These measurements are descriptive; individual images were not treated as independent biological replicates, and no inferential statistical comparisons were performed. **(j)** Representative merged CLSM images showing the spatial distribution of *B. subtilis* (green), *S. oneidensis* (red) and *E. coli* (blue) within MESIF pores across the indicated material environments on days 1, 4 and 6. Scale bar, 20 µm.

The response patterns differed substantially during consortium cultivation (Fig. 4d–f). *B. subtilis* exhibited particularly prominent fluorescence under APTES conditions, whereas *S. oneidensis* maintained comparatively strong signals in several environments, including the DNA hydrogel. By contrast, *E. coli* fluorescence remained highest in native MESIF and comparatively low in catalase-, ODN- and DNA hydrogel-functionalized environments (Fig. 4d–f; Supplementary Fig. S9g–l). Thus, the DNA hydrogel was associated with divergent fluorescence phenotypes among consortium members rather than a uniform reduction in species-associated signals.

Spatially resolved CLSM revealed heterogeneous, temporally changing fluorescence distributions within individual MESIF pores (Fig. 4g–j and Supplementary Fig. S10a–c). Comparison of CLSM-derived fluorescence with ROI-based whole-MESIF measurements showed species- and time-dependent correspondence (Supplementary Fig. S11a–d). Detailed analyses of the temporal and spatial responses are provided in the Extended Description to Supplementary Figs. S8–S11.

Together, these findings demonstrate that material-associated microbial phenotypes extend beyond monocultures and differ among members of a defined consortium. The DNA hydrogel provides a particularly revealing example: despite the comparatively weak *E. coli* fluorescence, *S. oneidensis* retained prominent fluorescence within the same DNA-rich material environment. This species-dependent response suggests that engineered extracellular materials may influence microbial community organization without affecting all constituent species equivalently. We next investigated whether these divergent phenotypes were accompanied by corresponding species-specific transcriptional states.

### Material-dependent phenotypes are accompanied by species-specific transcriptional states

To investigate whether the distinct physiological and spatial phenotypes of the three-species consortium were accompanied by corresponding molecular responses, we performed metatranscriptomic profiling of MESIF-associated communities after seven days of cultivation. Species-resolved transcriptomes were obtained from *Bacillus subtilis, Shewanella oneidensis* and *Escherichia coli* grown together in native, APTES-, GPTSh-, PEG-, catalase-, ODN- and DNA hydrogel-functionalized material environments, with native monocultures providing a reference for community context. Principal component analysis revealed distinct patterns of transcriptional variation among the three species, including a pronounced separation of DNA hydrogel-associated *B. subtilis* transcriptomes (Extended Data Fig. 1a–c). Differential expression analysis identified material-associated responses across multiple functional categories, with species-dependent distributions of significant genes (Extended Data Fig. 1d). Comparisons between native monoculture and consortium cultivation revealed a different, generally more restricted pattern of detectable transcriptional differences (Extended Data Fig. 1e). These findings indicate that the same material environments are associated with divergent transcriptional responses across species.

The underlying expression profiles revealed distinct physiological programs (Extended Data Fig. 1f–h). In *B. subtilis*, material-associated variation encompassed matrix production, motility, competence, stress regulation and developmental pathways, including sporulation-associated regulators and sigma factors (Extended Data Fig. 1f). These signatures provide molecular context for its fluorescence dynamics (Fig. 4d; Supplementary Fig. S9j), without establishing sporulation. *S. oneidensis* exhibited variation in adhesion, motility, c-di-GMP signaling, stress responses and respiratory functions, including extracellular electron-transfer-associated genes (Extended Data Fig. 1g). In *E. coli*, general, proteotoxic, oxidative and SOS stress-related transcripts complemented the independently measured RpoS-, RpoH- and SOS-associated reporter responses (Fig. 3; Extended Data Fig. 1h). Notably, the strong transcriptional separation of *B. subtilis* under DNA hydrogel conditions, alongside the contrasting *S. oneidensis* and *E. coli* fluorescence phenotypes (Fig. 4e,f), further highlights the species-specific nature of responses to this DNA-rich environment.

**Extended Data Fig 1.**
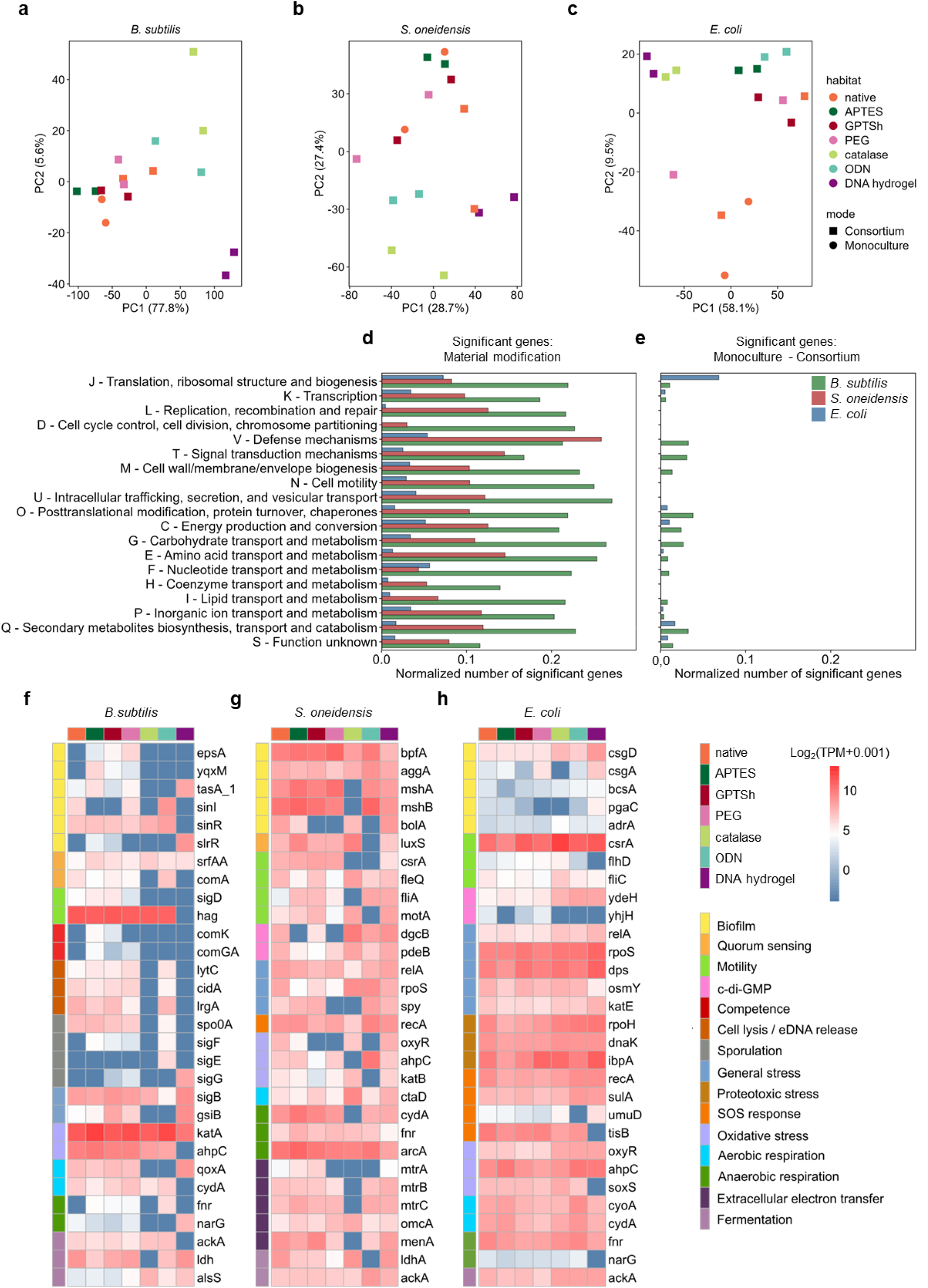
Distinct material environments elicit species-specific transcriptional responses in a three-species microbial consortium. A consortium comprising GFP-expressing *Bacillus subtilis*, RFP-expressing *Shewanella oneidensis* and BFP-expressing *Escherichia coli* was cultivated in MESIF presenting native, APTES-, GPTSh-, PEG-, catalase-, ODN- or DNA hydrogel-functionalized material environments. Metatranscriptomic profiling was performed after 7 days of cultivation. **(a–c)** Species-resolved principal component analysis (PCA) of variance-stabilized transcript abundances for *B. subtilis* (a), *S. oneidensis* (b) and *E. coli* (c). PCA was performed independently for each species using its 500 most variable genes across the analysed samples. Colors indicate material environments, and symbols distinguish consortium cultivation (squares) from native monoculture controls (circles). Each point represents an individual metatranscriptomic sample. Axis labels indicate the proportion of variance explained by each principal component. **(d)** Proportion of significantly differentially expressed genes within individual Clusters of Orthologous Groups (COG) functional categories in response to material environment, analysed separately for each species. Differential expression was assessed within consortium samples using DESeq2 likelihood-ratio tests comparing a full model containing material environment (∼ habitat) with a reduced intercept-only model (∼ 1). **(e)** Proportion of significantly differentially expressed genes between native monoculture and native consortium cultivation within the corresponding COG categories. Differential expression was assessed separately for each species using DESeq2 likelihood-ratio tests on native MESIF samples, comparing a full model containing cultivation condition (∼ detail) with a reduced intercept-only model (∼ 1). For each species and comparison, the number of significantly differentially expressed genes (Benjamini–Hochberg-adjusted *p* < 0.05; no fold-change cutoff) was divided by the total number of annotated genes assigned to the respective COG category. Genes assigned to multiple COG categories could contribute to more than one category. **(f–h)** Transcript abundance profiles of manually selected genes associated with biofilm formation, quorum sensing, motility, c-di-GMP signalling, competence, cell lysis and extracellular DNA release, sporulation, stress responses, respiration, extracellular electron transfer and metabolic processes in *B. subtilis* (f), *S. oneidensis* (g) and *E. coli* (h). Genes were selected according to their annotated functions and biological relevance, rather than on the basis of statistical significance. Columns represent the seven material environments under consortium cultivation. Transcript abundances were calculated independently for each metatranscriptomic sample using species-specific scaling factors. Transcripts per million (TPM) values were subsequently averaged across biological replicates within each condition and the resulting mean TPM values were log_2_-transformed after addition of a pseudocount of 0.01. Colored annotations indicate material environments and functional gene categories. Heatmap colors represent transcript abundance rather than statistical significance of differential expression.

Analysis of species-specific contributions to the recovered metatranscriptome revealed a pronounced material-dependent redistribution of transcriptional signals (Supplementary Fig. S12). Under DNA hydrogel conditions, *S. oneidensis* accounted for a substantially greater proportion of the total metatranscriptomic signal than under the other material environments, accompanied by a reduced relative contribution from *E. coli*. Because transcript abundances were normalized jointly across all three species within each sample, these differences reflect changes in their relative contributions to the recovered metatranscriptome rather than merely differences in mean gene expression. Although they do not directly quantify species abundance or per-cell transcriptional activity, these findings provide an additional molecular dimension to the contrasting species-specific fluorescence phenotypes observed in Fig. 4. Detailed analyses of the transcriptional programs, functional categories and their physiological implications are provided in the Extended Description accompanying Supplementary Fig. S12.

Together, these findings connect material-associated physiological and spatial phenotypes with distinct species-resolved transcriptional states and reveal a marked shift in the relative transcriptional contributions of consortium members under DNA hydrogel conditions. We next asked whether such material-dependent effects remain detectable during community succession in taxonomically complex environmental microbiomes.

### Material environments shape microbial community assembly during environmental succession

To determine whether material-dependent microbial responses extend to taxonomically complex environmental communities, we exposed functionalized MESIF to wastewater-derived microbiomes in two complementary settings: direct cultivation in a chemical wastewater treatment plant (WWTP) for up to 21 days and prolonged cultivation in a wastewater-derived mesocosm for up to 100 days (Supplementary Fig. S13a,b). The interconnected MESIF matrix permits exchange with the surrounding medium, allowing microbial communities to assemble under sustained environmental exposure. Initial colonization was accompanied by material-associated differences in alpha diversity, with DNA hydrogel-associated communities showing comparatively low Shannon diversity that subsequently increased during cultivation (Supplementary Fig. S13c,d).

Despite this temporal development, community composition remained strongly associated with the material environment. Bray–Curtis-based ordination revealed distinct material-associated community profiles in both experimental settings, including pronounced differentiation of DNA hydrogel-associated communities in the mesocosm (Fig. 5a,b). Across the complete time series, material environment explained 39% of compositional variation in the WWTP and 15% in the mesocosm (PERMANOVA, *p* = 0.001), whereas sampling time explained 16% and 43%, respectively (Fig. 5c,d). Material environment and sampling time were assessed in separate models, revealing contrasting associations with community composition across the two cultivation regimes. Thus, material-associated differentiation was particularly prominent during direct WWTP exposure, while temporal succession showed the stronger association during prolonged mesocosm cultivation.

**Fig 5.**
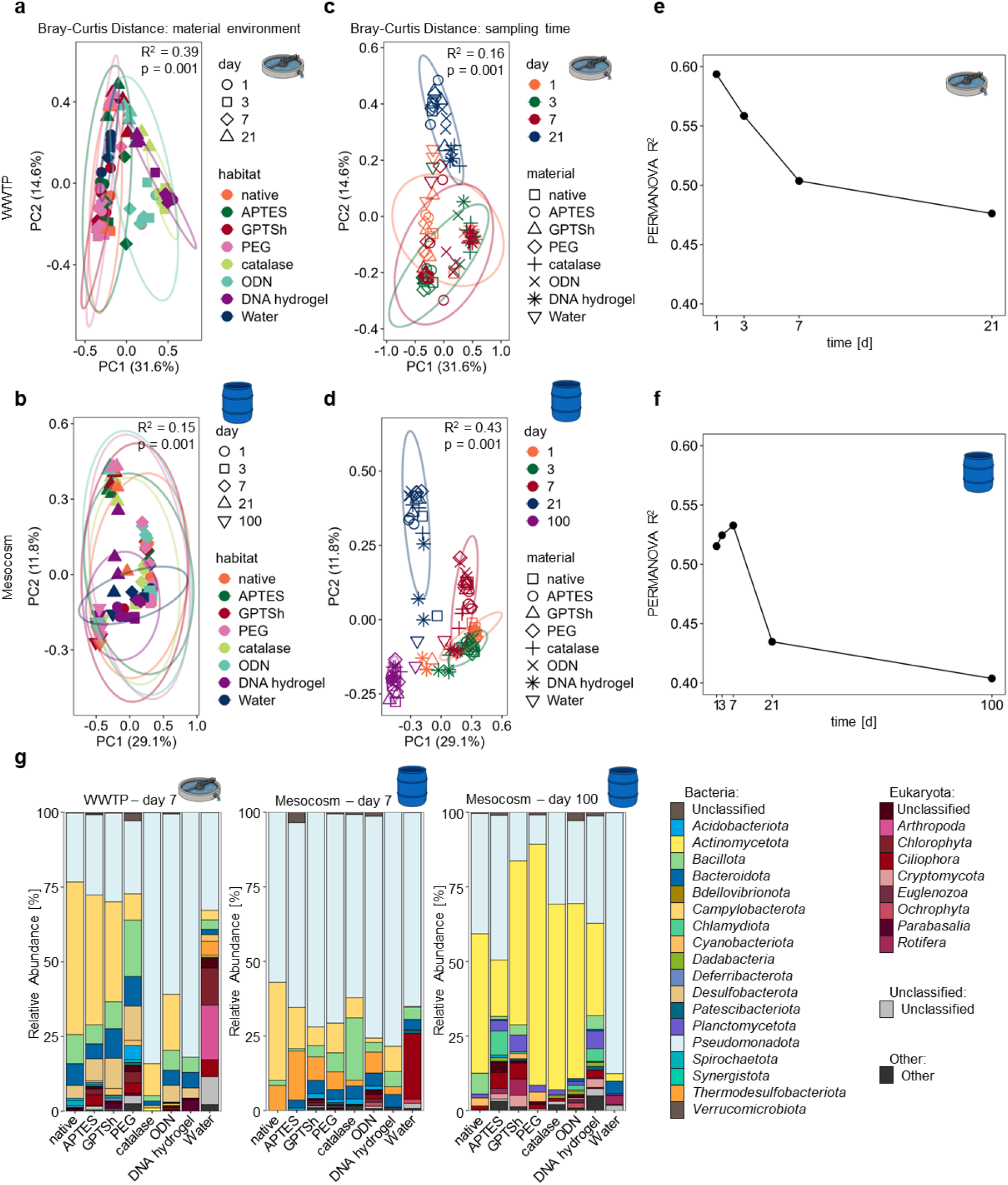
Material environments shape microbial community assembly during environmental succession. MESIF presenting native, APTES-, GPTSh-, PEG-, catalase-, ODN- or DNA hydrogel-functionalized material environments were exposed to complex environmental microbial communities in a chemical wastewater treatment plant (WWTP) for up to 21 days or in a wastewater-derived mesocosm for up to 100 days. **(a–d)** Principal coordinate analysis (PCoA) of Bray–Curtis dissimilarities calculated from 16S rRNA gene amplicon sequencing data for WWTP (a) and mesocosm (b) communities. Samples are coloured by material environment and symbols indicate sampling time. The same datasets are colored by sampling time, with symbols indicating material environment for the WWTP (c) and mesocosm (d), respectively. Individual MESIF chips represent independent biological replicates and were treated as independent observations. Different MESIF chips were sampled at each cultivation time, and PERMANOVA was performed using unrestricted permutations. Ellipses indicate 95% confidence regions estimated from multivariate *t*-distributions. Water samples are included as environmental controls. Differences in community composition were assessed by PERMANOVA using Bray–Curtis dissimilarities and 999 permutations. Material environment and sampling time were evaluated in separate models for each experimental setting. Corresponding PERMANOVA R^2^ and *p* values are indicated in each panel. **(e,f)** Temporal changes in the proportion of community compositional variation explained by material environment in the WWTP (e) and mesocosm (f). For each sampling time, a separate PERMANOVA was performed using Aitchison distances calculated from centred log-ratio (CLR)-transformed ASV abundances, with material environment as the sole explanatory variable and 9,999 permutations. Water controls were excluded from these analyses and only the seven MESIF material environments were considered. R^2^ values indicate the proportion of compositional variation explained by material environment at each time point. Benjamini–Hochberg-adjusted *p* values were below 0.005 at all time points. Homogeneity of multivariate dispersion was assessed using PERMDISP with 9,999 permutations. A significant difference in dispersion was detected only at WWTP day 1 (adjusted *p* = 0.0245). **(g)** Taxonomic composition of MESIF-associated communities determined by shotgun metagenomic sequencing after 7 days in the WWTP and after 7 and 100 days in the mesocosm. Stacked bars show the relative representation of taxonomically assigned small- and large-subunit ribosomal RNA (SSU and LSU rRNA) marker sequences identified using MDMcleaner and classified with SINA against the SILVA database. The category “Unclassified” comprises SSU and LSU rRNA marker sequences that could not be assigned taxonomically. Biological replication comprised *n* = 3 independent MESIF chips per condition for 16S rRNA gene amplicon sequencing; metagenomic profiles represent *n* = 1 dataset per condition.

We next asked whether the material-associated differences persisted as microbial communities underwent succession. Time-resolved analysis revealed that material environment remained significantly associated with community composition at every sampling point, although its explanatory contribution declined over time (Fig. 5e,f). The proportion of variation explained by material environment decreased from approximately 0.59 to 0.48 in the WWTP and from approximately 0.52 to 0.40 in the mesocosm. Notably, material environment still explained approximately 40% of compositional variation after 100 days of mesocosm cultivation (Fig. 5f). Prolonged succession therefore did not erase material-associated ecological differentiation. Yet these persistent material-associated differences did not reflect static community states. Material-specific communities followed distinct temporal trajectories, and their relative compositional relationships were progressively reorganized (Supplementary Fig. S14a–d). Cross-time Aitchison distances further revealed heterogeneous changes among material environments (Supplementary Fig. S14e,f). Engineered material environments thus left a persistent ecological imprint on community assembly, even as succession progressively reorganized community composition.

The two DNA-containing material environments were particularly informative. ODN- and DNA hydrogel-functionalized MESIF generated distinguishable community profiles (Fig. 5a,b) and temporal trajectories (Supplementary Fig. S14a,b), demonstrating that extracellular DNA-containing environments are not ecologically interchangeable. Unlike ODN functionalization, the DNA hydrogel presents a highly concentrated, spatially organized three-dimensional DNA matrix within the microbial habitat. Because these materials differ in DNA abundance, polymer length, local density and physical organization, the observed differences cannot be attributed to three-dimensional DNA architecture alone. Nevertheless, their divergent ecological responses suggest that the influence of extracellular DNA depends not merely on its presence, but also on its local abundance and presentation as a material.

Shotgun metagenomic sequencing provided complementary taxonomic and functional perspectives. Community taxonomic profiles differed among material environments and changed substantially during prolonged mesocosm cultivation (Fig. 5g). Functional gene profiles also varied across materials, including categories associated with surface-associated lifestyles, signaling, DNA processing, stress responses and metabolism (Supplementary Fig. S15). ODN- and DNA hydrogel-associated communities also displayed distinct functional gene profiles, extending their observed ecological differences beyond amplicon-based community composition (Supplementary Fig. S15). Genome-resolved analysis revealed heterogeneous relative abundance patterns among reconstructed microbial populations (Supplementary Fig. S16 and Supplementary Table S4). These metagenomic comparisons were descriptive and did not establish preferential recruitment or differential functional activity. A detailed analysis of alpha diversity, temporal community trajectories, functional gene profiles and population-level dynamics is provided in the Extended Description to Supplementary Figs. S13–S16.

These findings extend the material-dependent physiological responses observed in defined systems to complex environmental microbiomes. Material-associated differences remained detectable throughout prolonged ecological succession, while ODN and DNA hydrogel produced distinguishable community trajectories and functional profiles (Fig. 5e,f; Supplementary Figs. S14a,b and S15). The concentrated, three-dimensional presentation of DNA in the hydrogel therefore emerges as a distinctive material feature associated with microbial ecological differentiation, although the contributions of DNA abundance and physical organization cannot yet be separated. These observations prompted us to investigate whether the DNA hydrogel might also interact with environmental microorganisms at the level of DNA sequence.

### Material-derived DNA yields sequence-defined junction signatures with microbial genomic DNA

Having established that the DNA hydrogel influences microbial physiology and community assembly, we next asked whether DNA introduced as an extracellular material could also interact with environmental microorganisms at the sequence level. We first tracked hydrogel-derived DNA during environmental cultivation using a sequence-specific qPCR assay targeting the RCA product. The proportion of qPCR-detectable hydrogel-derived DNA declined rapidly in the wastewater treatment plant (WWTP) and more gradually in the wastewater-derived mesocosm, where hydrogel-derived sequences remained detectable after 100 days (Fig. 6a,b). Thus, the material-derived DNA signal persisted within the environmental system even as its relative abundance decreased substantially.

**Fig 6.**
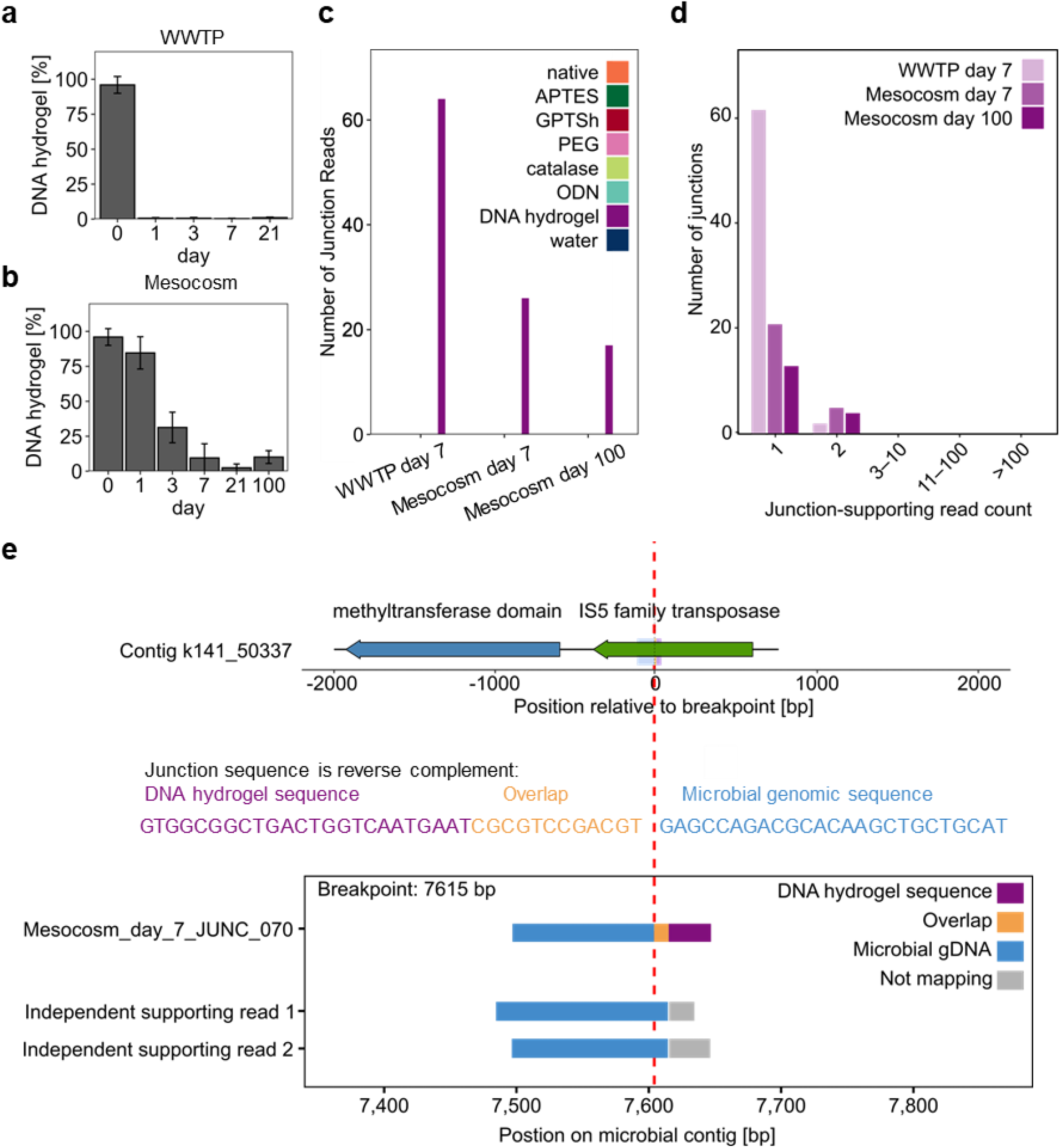
Material-derived DNA yields candidate junctions with microbial genomic sequences. **(a,b)** Relative abundance of DNA hydrogel-derived sequences during environmental cultivation in the wastewater treatment plant (WWTP; a) and wastewater-derived mesocosm (b). Hydrogel-derived DNA was quantified by sequence-specific qPCR and expressed as a percentage of the total extracted DNA. The y-axis label “DNA hydrogel [%]” therefore denotes the relative abundance of qPCR-detectable hydrogel-derived sequences rather than the proportion of physically intact DNA hydrogel. Bars indicate mean values and error bars indicate standard deviations from n = 3 biological replicates. **(c)** Number of distinct candidate DNA hydrogel–microbial DNA junctions identified in metagenomic datasets after 7 days in the WWTP and after 7 and 100 days in the mesocosm. A total of 64, 26 and 17 candidates, respectively, passed the junction-detection and filtering procedure. Candidate junctions were identified from sequencing reads containing regions aligning to the DNA hydrogel template and microbial genomic sequences; junction definition and filtering criteria are described in Methods and Supplementary Table S5. **(d)** Distribution of the number of independent read pairs supporting individual candidate junctions across the three environmental datasets. Independent read-pair counts were determined by breakpoint-specific rescreening of paired-end sequencing datasets after exclusion of PCR duplicates. Most candidates were supported by one independent read pair, whereas a smaller number were supported by two independent read pairs. **(e)** Genomic context and sequencing-read support for the candidate junction *Mesocosm_day_7_JUNC_070*. The upper panel shows a 4.0-kb region of the assembled microbial contig k141_50337, containing genes annotated as a methyltransferase domain-containing protein and an IS5 family transposase. The candidate junction sequence is displayed in reverse-complement orientation relative to the contig. The dashed red line marks the computationally inferred breakpoint at the transition between the microbial-specific and overlapping sequence regions. The lower panel illustrates the reference-alignment structure of the candidate junction sequence and two independent supporting reads. Blue denotes the microbial-specific segment aligning to the assembled contig but not to the DNA hydrogel reference, orange denotes the overlapping segment aligning to both references, and lilac denotes the segment aligning to the DNA hydrogel reference but not to the assembled microbial contig. Grey indicates read segments that do not map to the respective reference in the displayed alignment. Two independent reads supported this candidate junction. These alignments establish a candidate sequence junction but do not demonstrate stable genomic integration or horizontal DNA transfer. Additional information on junction detection, genomic context, alignments and associated sequences is provided in Supplementary Figs. S17–S21 and Supplementary Tables S5 and S6.

We then interrogated the metagenomic datasets for sequencing reads containing candidate junctions between the DNA hydrogel template and sequences mapping to microbial genomic DNA. The detection pipeline identified reads with directly adjacent or overlapping hydrogel- and microbial-mapping segments, followed by paired-end mapping, sequence filtering and breakpoint-specific rescreening across datasets (Methods; Supplementary Tables S5 and S6). Strikingly, candidate junctions were detected exclusively in DNA-hydrogel-exposed communities and were absent from the other material environments and water controls examined (Fig. 6c, Supplementary Fig. S17). We identified 64 distinct candidates after 7 days in the WWTP, 26 after 7 days in the mesocosm and 17 after 100 days in the mesocosm. Most candidates were supported by a single independent read pair, whereas a smaller subset was supported by two independent read pairs following exclusion of PCR duplicates (Fig. 6d, Supplementary Fig. S17). Junction-supporting sequences occurred in both hydrogel-to-microbial and microbial-to-hydrogel orientations and were recovered from both members of paired-end reads. Together, these observations reveal material-specific sequence associations across independent environmental cultivation settings.

The microbial-mapping portions of these reads could be assigned to assembled metagenomic contigs, placing the candidate junction signatures within diverse microbial genomic neighborhoods (Supplementary Figs. S18–S20). These regions contained genes associated with metabolism, transport, regulation and other cellular functions, alongside hypothetical proteins. Notably, mobile-element- or phage-associated genes occurred within 8 of the 64 WWTP day-7 and 7 of the 26 mesocosm day-7 junction-associated regions, but were not identified within the corresponding ±2-kb windows of the 17 mesocosm day-100 candidates (Supplementary Fig. S21). The heterogeneous functional contexts argue against a single genomic setting underlying the observed sequence associations, while the proximity of mobile-element-associated genes raises mechanistic questions that cannot be resolved from genomic neighborhood alone.

Breakpoint-specific rescreening did not identify any candidate junction recurring across the environmental datasets under the applied sequence-matching criteria (Supplementary Fig. S17). To examine the sequence-level evidence underlying an individual candidate, we focused on *Mesocosm_day_7_JUNC_070*, identified after 7 days of mesocosm cultivation (Fig. 6e, Supplementary Fig. S19). The microbial-mapping portion of this candidate aligned to an assembled contig containing genes annotated as a methyltransferase domain-containing protein and an IS5 family transposase (contig k141_50337). The candidate sequence contained a microbial-specific segment, a short region aligning to both the microbial contig and the DNA hydrogel reference, and a hydrogel-reference-specific segment that did not align to the assembled contig. Two independent reads supported the candidate junction, with microbial-mapping segments associated with the same genomic region. This read-level evidence provides a defined example of a material-associated sequence junction, although neither the genomic integration of the hydrogel-specific segment nor a mechanistic contribution of the neighboring genes can be inferred from these alignments alone.

Collectively, these results reveal a previously unexplored informational dimension of engineered DNA materials in microbial environments. DNA introduced as an extracellular hydrogel was associated with rare, material-specific sequence signatures linking hydrogel-derived DNA to microbial genomic sequences. Although the current short-read evidence does not establish stable genomic incorporation, inheritance or horizontal transmission, these findings suggest that extracellular DNA materials may provide a route for the transfer of sequence information into microbial genomes. This possibility extends the biological significance of extracellular DNA beyond its structural and physiological roles and raises the intriguing prospect that eDNA within natural extracellular polymeric substance (EPS) matrices may also participate in genetic interactions between the extracellular environment and microbial cells.

## Conclusion

Our results establish engineered extracellular materials as active determinants of microbial physiology and ecological selection. Across increasing levels of biological complexity, material composition shaped stress-associated physiological responses, species-specific behavior within a defined consortium and the succession of complex environmental microbiomes. These effects extended beyond differences in growth-associated fluorescence and remained evident during prolonged environmental cultivation. Material-associated community differentiation remained detectable after 100 days of mesocosm succession, even as community composition was extensively reorganized (Fig. 5e,f; Supplementary Fig. S14). Thus, the extracellular material environment is not merely a substrate for microbial colonization but an ecological variable capable of shaping microbial community assembly over extended timescales.

DNA was particularly consequential in this context. Extracellular DNA (eDNA) is a multifunctional component of microbial extracellular polymeric substances (EPS), contributing to adhesion, matrix organization and biofilm development as well as nutrient source.^[22,23,36]^ Its organization into supramolecular networks further demonstrates that DNA architecture is an important feature of natural biofilm matrices.^[26]^ Here, we extend this concept by investigating DNA as an engineered extracellular biomaterial. An RCA-generated DNA hydrogel, presenting DNA at high local concentration within a three-dimensional polymeric matrix, elicited a distinct stress-associated physiological response in *E. coli*, produced species-dependent phenotypes and transcriptional responses within a defined consortium, and was associated with distinct community assembly during prolonged cultivation of wastewater-derived microbiomes. Importantly, surface-bound oligodeoxynucleotides (ODNs) produced different biological responses, including distinguishable community trajectories and functional gene profiles (Supplementary Figs. S14a,b and S15). This distinction identifies the local abundance and material organization of DNA as critical dimensions of the material–microbe interface, rather than treating extracellular DNA simply as a molecular component whose presence or absence determines biological behavior. The individual contributions of DNA abundance, polymer length, network architecture and local accessibility remain to be disentangled.

A further distinction makes DNA unusual among engineered biomaterials: its structural properties and biological information are encoded in the same molecule. Beyond its effects on microbial physiology and community assembly, the DNA hydrogel introduced identifiable sequences into the extracellular environment. Metagenomic analysis revealed rare, sequence-defined candidate junctions between hydrogel-derived DNA and sequences mapping to microbial genomes exclusively in DNA-hydrogel-exposed communities. These findings extend the biological significance of the hydrogel beyond its role as a physical extracellular environment and reveal a previously unexplored interface between engineered DNA materials and microbial genomic sequence space.

Although the present short-read evidence does not establish stable genomic incorporation, inheritance or horizontal transmission, the material-specific sequence signatures raise the possibility that extracellular DNA materials may provide a route for sequence information to enter microbial genomes. Establishing whether these signatures reflect genuine genetic exchange will require direct experimental validation.

Engineered DNA materials thus unite two biologically consequential properties: their concentrated, spatially organized extracellular presentation can shape microbial physiology and ecological selection, while their encoded sequences introduce the possibility of interactions with microbial genomic information. This dual role offers a new perspective on eDNA-rich biofilm matrices. The concept of extracellular polymeric matrices as reservoirs of genetic information has been proposed previously,^[37]^ and experimental studies have demonstrated that extracellular DNA within biofilms or associated with environmental materials can remain available for natural transformation.^[38–41]^. Our findings extend this perspective by suggesting that concentrated, spatially organized DNA networks within natural EPS matrices may represent dynamic interfaces between extracellular material architecture and microbial genetic exchange. This possibility also opens opportunities to engineer extracellular environments towards contrasting technological objectives, from promoting productive microbial communities in biohybrid materials and bioprocesses to suppressing unwanted colonization in antifouling and biofilm-control applications. More broadly, our findings provide a framework for designing not merely the surfaces on which microorganisms grow, but the extracellular material environments that shape their behavior and ecological trajectories.

## Author contributions

Conceptualization, C.M.N.; Methodology, L.M.; Investigation, L.M., M.V.M, S.M., P.S.J.; Data curation, L.M.; Microscopy, L.M. and S.W., Formal analysis, L.M. and M.V.M.; Visualization, L.M.; Supervision, K.S.R., A.K.K. and J.V.; Resources, A.K.K. and J.V.; Writing – original draft, L.M.; Writing – review & editing, L.M., M.V.M., C.M.N., K.S.R., A.K.K. and J.V.

## Acknowledgements

This work was financially supported through the Helmholtz program “Materials Systems Engineering” under the topic “Adaptive and Bioinstructive Materials Systems” (43.33.11), KIT EXU project DigiteLiSE and BMBF project 161L0284A MicroMatrix. The authors thank Anna Edinger, Cornelia Ziegler, David Thiele and Hagen Rießland for experimental help.

