## Supplementary Information for "Engineered DNA materials shape microbial physiology and ecological selection"

### 1. Materials and Methods

#### 1.1. Materials

All chemicals were purchased from Fisher Scientific, Merck, Roth, Sigma Aldrich or VWR, if not stated otherwise.

#### 1.2. Preparation of MESIF Chips

The preparation of the MESIF has been described in detail previously by Zoheir *et al.*[1] Casting molds for both the chip housing and the fluidic chamber were designed using Autodesk Inventor 2020 software (Autodesk Inc., USA) and subsequently fabricated from poly(methyl methacrylate) (PMMA) using a CNC micromilling machine (Mini Mill GX, Minitech Machinery, USA).

Both the MESIF and its housing were fabricated from polydimethylsiloxane (PDMS). For this purpose, PDMS prepolymer and curing agent (Sylgard 184, Dow Corning, USA) were mixed at a ratio of 10:1 and degassed under vacuum for 30 min to remove entrapped air bubbles. For MESIF fabrication, the PDMS mixture was poured into the prepared molds and filled with salt crystals (500–707 µM). The molds were then centrifuged at 4600 rpm and 4 °C for 20 min (5804 R, Eppendorf, Germany) to obtain a densely packed porogen structure. Subsequently, the PDMS–porogen composite was cured at 70 °C for 1 h in an oven (Heraeus, Thermo Scientific Inc., USA). After curing, the solidified composite (approximately 65 × 10 × 3 $mM^3$) was carefully removed from the mold and placed in warm water. The salt crystals were leached out by repeated manual compression of the PDMS structure. To ensure complete removal of residual salt, the samples were further stirred in water at 40 °C for 48 h.

The chip housing and fluidic chamber were fabricated separately by pouring the degassed PDMS mixture into the corresponding PMMA molds, followed by curing at 70 °C for 30 min. For assembly, the MESIF was cut into uniform pieces (18 × 10 × 3 $mM^3$) using a scalpel. Depending on the experimental requirements, the MESIF was either used in its native state or subjected to material modification as described below (see Section S1.3) prior to integration into the chip.

For chip assembly, 4 mm diameter environmental contact holes were manually punched into the housing lid. For MESIF, whose surface was to be chemically modified, both the lid and the housing were treated with oxygen plasma (30 s, 300 W, 0.2 mbar; SmartPlasma 10, PlasmaTechnology, Germany) to promote bonding. The MESIF was then placed in the housing, covered with a lid, and bonded by applying pressure at 90 °C for 30 min to achieve permanent sealing. For biomolecule functionalization, an alternative assembly protocol was employed. In this case, the housing and lid were coated with degassed PDMS, after which the MESIF was placed in the housing base. The lid was applied, and the entire assembly was fixed under pressure and cured overnight to ensure stable integration.

The fluidic chamber was assembled by placing a MESIF into its designated cavity and sealing it with an adhesive foil (HJ-Bioanalytik, Germany). The chamber was equipped with two syringe connections, allowing controlled perfusion of solutions through the porous MESIF structure.

#### 1.3. Material Modification of MESIF

##### 1.3.1. Surface silanization with APTES and GPTS

To achieve APTES and GPTS material environments, silanization was performed following a modified protocol based on Shu *et al.*[2] The MESIF surfaces were first activated using an aqueous solution composed of dd$H_2O$, concentrated hydrochloric acid (37%), and concentrated hydrogen peroxide (30%) in a 5:1:1 ratio. For activation, the solution was placed in a mortar, and the MESIF were mechanically squeezed using a pestle for 2 min, followed by stirring for 3 min. This activation cycle was repeated five times to ensure thorough surface treatment. Subsequently, the MESIF were rinsed repeatedly in fresh dd$H_2O$ using the same mechanical treatment until the pH of the rinse water reached pH 7. The washed MESIF were then dried at 90 °C for at least 6 h. For silanization, a 2% (v/v) solution of either 3-aminopropyltriethoxysilane (APTES) or 3-

glycidyloxypropyltrimethoxysilane (GPTS) in toluene was prepared. The activated MESIF were incubated in the respective silane solution at 80 °C and 300 rpm for 24 h. Following incubation, the samples were washed three times with toluene and subsequently three times with dichloromethane under continuous stirring to remove unbound silane. Residual solvent was removed using a rotary evaporator (Laborota 4000 – efficient, Heidolph, Germany) at 40 °C under reduced pressure. For GPTSh modification, the epoxy groups of the GPTS functionalized MESIF were opened under acidic conditions by treating the samples in a 1:1 (v/v) solution of hydrochloric acid and ddH$_2$O for 25 min.

1.3.2. Material modification with PEG

PEG modification of the MESIF was carried out according to the protocol described by Gökaltun *et al.*[3] A 0.25% (w/w) PDMS–PEG block copolymer (DBE-712, Gelest, USA) was added to the PDMS prepolymer and thoroughly mixed prior to vacuum treatment. The subsequent fabrication of the MESIF was then performed following the standard preparation protocol described above.

The chemically modified MESIF were assembled into chips and sterilized by autoclaving at 120 °C for 21 min and 1 bar overpressure.

1.3.3. Catalase functionalization

The thermostable catalase TaCAT from *Thermoanaerobaculum aquaticum* was selected because of its central role in microbial oxidative stress defense through the decomposition of reactive oxygen species (ROS) and modulation of oxygen availability.[4] To enable site specific immobilization, TaCAT was fused to a SpyTag (ST) peptide (TaCAT-ST-His, while surface attachment was mediated by SpyCatcher (SC). .[5] Plasmids encoding SpyCatcher (His SC) and catalase (TaCAT ST His; Table S2) were designed and cloned into pET expression vectors carrying an ampicillin resistance marker. Insertion of larger DNA fragments (>100 bp) was achieved using Gibson assembly according to established protocols.[6] Following cloning, reaction mixtures were treated with DpnI to degrade residual template plasmid DNA prior to transformation into *E. coli* DH5α. Plasmid isolation was performed using a ZR Plasmid Miniprep–Classic kit (Zymo Research, Germany) in accordance with the manufacturer's guidelines. Correct construct assembly and sequence fidelity were confirmed by commercial sequencing (LGC Genomics, Germany). Primers were purchased from Sigma-Aldrich, and synthetic DNA fragments were obtained via the GeneArt Strings service (Thermo Fisher Scientific, USA). Recombinant protein production was performed as described previously.[7,8] Briefly, *E. coli* BL21 (DE3) cells harboring the respective plasmids were cultured overnight in a total volume of 2.5 L and protein expression was induced with 0.1 mM IPTG. Cultures expressing the catalase were additionally supplemented with 200 µM 5-Aminolevulinic acid. Cells were collected by centrifugation (10,000 × g, 10 min, 4 °C) and resuspended in NPi-10 buffer (50 mM NaH$_2$PO$_4$, 300 mM NaCl, 10 mM imidazole, pH 8.0). Cell disruption was achieved by ultrasonication (Sonopuls17 MS73, Bandelin, Germany), followed by removal of insoluble material via high-speed centrifugation (45,000 × g, 1 h, 4 °C). The clarified lysate was filtered through a 0.45 µM PVDF membrane and applied to a Ni-NTA affinity column (HisTrap FF, 5 mL, GE Healthcare) connected to an ÄKTA Pure chromatography system (Akta Prime Plus, GE Healthcare, USA). After equilibration and washing with NPi-10 buffer, bound His-tagged proteins were eluted using an imidazole gradient up to 500 mM (NPi-500 buffer: 50 mM NaH$_2$PO$_4$, 300 mM NaCl, 300 mM Imidazol, pH 8,0). Finally, buffer exchange to PBS (150 mM NaCl, 23 mM KH$_2$PO$_4$, 77 mM K$_2$HPO$_4$, pH 7.5) was carried out using centrifugal filtration devices (10 kDa MWCO, GE Healthcare). The purified proteins were analyzed by discontinuous SDS–polyacrylamide gel electrophoresis (SDS-PAGE) using Laemmli midi-gels, followed by Coomassie staining. Molecular weights were estimated by comparison with a prestained broad-range protein ladder (Color Prestained Protein Standard, New England Biolabs, USA).

For protein immobilization, GPTS modified MESIF were placed in a fluidic chamber (Fig. S1b) and incubated with 350 µL of a 250 µM SC solution in PBS for 24 h to enable covalent immobilization to the surface, due to epoxy–amine coupling with lysine residues. Following incubation, the MESIF were rinsed with 3 mL PBS to remove unbound protein. Remaining reactive epoxide groups on the surface were subsequently blocked by treatment with 1 M ethanolamine for 3 h. After an additional washing step with 3 mL PBS, a 250 µM TaCAT-ST solution in PBS was introduced into the MESIF and incubated for 24 h to allow the formation of the intermolecular isopeptide bond with the SC. Finally, the MESIF were rinsed with 3 mL $ddH_2O$ to remove residual unbound TaCAT-ST. The concentrations of SC and TaCAT in the collected effluent were quantified to determine immobilization efficiency.

1.3.4. ODN functionalization

For oligonucleotide (ODN) functionalization, GPTS modified MESIF were used. The MESIF were placed into the fluidic chamber and saturated with 350 µL of a 100 nm solution of 5'-Amine-$C_{12}$-$T_8$-$(CAT)_8$-3' in PBS (150 mM NaCl, 23 mM $KH_2PO_4$, 77 mM $K_2HPO_4$, pH 7.5). The oligonucleotide solution was incubated overnight, allowing covalent epoxy–amine coupling, followed by rinsing the next day with 3 mL PBS. Subsequently, the complementary strand (5'-$(ATG)_8$-3') was introduced at a concentration of 100 nm in PBS and incubated overnight under identical conditions. Excess oligonucleotide was removed by rinsing with 3 mL $ddH_2O$ on the following day.

1.3.5. Functionalization with DNA hydrogel

To prepare the MESIF for DNA hydrogel synthesis, GPTS modified MESIF were first functionalized with an amino modified primer. For this purpose, a 100 nm solution of 5'-Amine-$C_{12}$-TTTTTTTTTTTTCCACCTACGGTGGCGGCTGA-3') in PBS was introduced into MESIF mounted in a fluidic chamber and incubated for at least 6 h. Subsequently, the MESIF were rinsed with 3 mL PBS to remove unbound primer. To prevent non-specific binding, the MESIF surfaces were blocked using denatured bovine serum albumin (BSA; 1% in PBS, preheated for 5 min at 95 °C). Blocking was performed for 6 h at room temperature, followed by rinsing with 3 mL $ddH_2O$. The synthesis of DNA hydrogels was carried out as described in detail by Moench *et al.*[9] For rolling circle amplification (RCA), a circularized DNA template was first generated. The template strand (5′-GTAGGTGGACACGCCGCATCCTACATCCAGCGTCAACGTCGGACGCGATTCATTGACCAG TCAGCCGCCACC-3′) and primer (5′-CCACCTACGGTGGCGGCTGA-3′) were mixed at final concentrations of 5 µM each in 1× TE buffer (20 mM Tris-HCl, 1 mM EDTA, pH 7.6) supplemented with 100 mM NaCl. Hybridization was performed at 25 °C with gentle shaking (400 rpm) for 1.5 h. For ligation, 100 µL of the hybridized DNA mixture was combined with 20 µL of 10× ligase buffer (New England Biolabs, USA), 78 µL $ddH_2O$, and 2 µL T4 ligase (400,000 $U{\cdot}mL^{-1}$, New England Biolabs, USA). The reaction was incubated at 25 °C for 3 h, followed by enzyme deactivation at 70 °C for 20 min. To remove residual linear DNA, 23 µL of 10× NEB buffer 1, 4 µL exonuclease I (20,000 $U{\cdot}mL^{-1}$), and 4 µL exonuclease III (100,000 $U{\cdot}mL^{-1}$) were added to 200 µL of the ligation mixture. The reaction was incubated overnight at 37 °C. Enzymes were subsequently inactivated at 95 °C for 10 min. The circularized DNA templates were purified using Vivacon ultrafiltration units (10 kDa cutoff, Sartorius, Germany) and washed five times with 0.1× TE buffer. DNA concentrations were determined using a Synergy plate reader (BioTek, USA), and samples were diluted to 1 µM. For rolling circle amplification, a total reaction volume of 350 µL (corresponding to one MESIF) was prepared consisting of 140 µL circular template (1 µM), 2.3 µL primer (1 mM), 46.6 µL dNTPs (10 mM), 23.3 µL 10× Phi29 reaction buffer (New England Biolabs, USA), 1.4 µL albumin (20 $mg{\cdot}mL^{-1}$), 113.1 µL $ddH_2O$, and 23.3 µL Phi29 polymerase (10,000 $U{\cdot}mL^{-1}$). The RCA mixture was introduced into the MESIF and incubated at 30 °C for 72 h to enable hydrogel formation. Following incubation, the MESIF were rinsed with 3 mL $ddH_2O$ to remove residual reaction components.

##### 1.4. Validation of material environments

1.4.1. Fluorescamine assay

Successful silanization of the MESIF was verified using a fluorescamine assay targeting surface exposed primary amino groups. A 3.5 mM fluorescamine solution in acetone was introduced into APTES modified and unmodified control MESIF. Reaction of fluorescamine with primary amines generated a highly fluorescent product exhibiting spectral properties similar to those of blue fluorescent protein. Fluorescence intensity was recorded at 480 nm using a fluorescence imaging system (Fusion FX, Vilber, France) equipped with an F 500 filter. The effectiveness of surface functionalization was evaluated by comparing fluorescence intensities between APTES modified and control samples.

##### 1.4.2. Catalase activity

Catalase activity was assessed based on the protein's ability to decompose hydrogen peroxide ($H_2O_2$) into water ($H_2O$) and oxygen ($O_2$). The decrease in $H_2O_2$ concentration was monitored spectrophotometrically by measuring absorbance at 240 nm. For quantitative analysis, calibration curves correlating absorbance to $H_2O_2$ concentration were recorded alongside each set of measurements.

For activity measurements catalase (100 nM) was diluted in PBS. Aliquots of 50 µL catalase solution and 200 µL $H_2O_2$ solution of varying concentrations were combined in a 96-well plate. Absorbance at 240 nm was recorded immediately using a Synergy plate reader (BioTek, USA) over a 60 min.

For activity measurements of immobilized catalase, MESIF were functionalized according to section 1.3.2 with 250 µM catalase were injected with 1 mL 73 mM $H_2O_2$ at a flow rate of 20 $\mu L \cdot min^{-1}$. Subsequently, the MESIF were rinsed with 1 mL PBS at a flow rate of 50 $\mu L \cdot min^{-1}$. The collected effluent fractions were analyzed by measuring their volume and determining the residual $H_2O_2$ concentration via absorbance at 240 nm.

##### 1.4.3. ODN hybridization analysis

For oligonucleotide hybridization studies, GPTS modified MESIF were placed in a fluidic chamber (Fig. S1b) and functionalized according to the previously described ODN protocol (Section 1.3.2). Fluorescently labeled oligonucleotides were used to monitor hybridization. The amino modified strand (5′-Amine-C12-T8-$(ATG)_4$-Cy3-3′) was labeled with Cy3, while the complementary strand (5′-Cy5-$(CAT)_4$-3′) carried a Cy5 label.

Fluorescence imaging was performed using both a fluorescence imaging system and confocal laser scanning microscopy (CLSM, LSM 880, Zeiss, Germany). Measurements included fluorescence intensity after immobilization of the Cy3-labeled strand (Fluorescence-Imaging-System: $\lambda_{ex}$ = 580 nm, F-550, 100ms; CLSM: $\lambda_{ex}$ = 561 nm $\lambda_{em}$ = 567 nm), Cy5 after hybridization (Fluorescence-Imaging-System: $\lambda_{ex}$ = 640 nm, F-650, 100ms; CLSM: $\lambda_{ex}$ = 633 nm $\lambda_{em}$ = 670 nm), quenching of Cy3 fluorescence upon hybridization with the complementary strand, and Förster resonance energy transfer (FRET), detected via Cy3 excitation and Cy5 emission (Fluorescenc -Imaging-System: $\lambda_{ex}$ = 580 nm, F-650, 100ms; CLSM: $\lambda_{ex}$ = 561 nm, $\lambda_{em}$ = 670 nm).

##### 1.4.4. DNA hydrogel

For visualization of the DNA hydrogel 500 nM Cy5 labeled dUTPs were spiked in the RCA mix prior to filling the mix into the MESIF in the fluidic chamber. For control, RCA mixtures without polymerase were used. The MESIF were incubated for 72 h at 30 °C in the dark and then gently rinsed with $ddH_2O$. Fluorescence images to confirm hydrogel formation in the MESIF were taken before and after rinsing using a fluorescence gel scanner ($\lambda_{Ex}$ = 640 nm, $\lambda_{Em}$ = 700 nm, 100 ms). A more detailed distribution of fluorescence from the Cyanine 5-labeled dUTPs and Cyanine 3-labeled oligonucleotides within the MESIF was obtained using CLSM (Cy5: $\lambda_{Ex}$ = 633 nm, $\lambda_{Em}$ = 670 nm).

Quantification of DNA hydrogel within the MESIF required extraction of DNA from the porous matrix. Consistent with the microbial DNA extraction procedure described in Section 1.8.3, the DNeasy PowerSoil Pro Kit (QIAGEN, Germany) was used according to the manufacturer's instructions, with minor modifications. For hydrogel DNA quantification, the entire MESIF was processed by mechanically compressing the material in CD1 lysis buffer to facilitate DNA release from the porous structure, followed by bead beating (two cycles of 1 min at 5.0 $m \cdot s^{-1}$; MP Biomedicals, USA). Extracted DNA was quantified fluorometrically using the Qubit™ dsDNA HS Assay Kit (Thermo Fisher Scientific, Germany) and by quantitative PCR (Section 1.8.4).

#### 1.5. Model organisms and fluorescent reports

In this study, three model organisms were used: *Escherichia coli* K12 MG1655 (*E. coli*), *Bacillus subtilis* 1012 wild type (*B. subtilis*), and *Shewanella oneidensis* MR-1 (*S. oneidensis*).

Plasmids utilized in *E. coli* were based on a pMK backbone carrying the arabinose-inducible promoter $P_{BAD}$ and encoded either eGFP, mRFP1, or mTagBFP2 (see Table S2), cloned as described above (Section S1.3.2). An RGB-S reporter plasmid was available in-house.[10] For *S. oneidensis*, mRFP1 in a pMK backbone was used as a fluorescence reporter. The plasmid used for *B. subtilis*, encoding folding reporter GFP, was obtained commercially (pHT10-gfp+ (PBS005C), Boca Scientific, USA). To ensure consistent antibiotic selection across all constructs, chloramphenicol resistance cassettes were replaced with kanamycin resistance (knt) cassettes.

Transformation of *E. coli* was performed by heat shock. An amount of 100 ng plasmid DNA was added to 60 µL competent cells, incubated on ice for 5 min, subjected to heat shock at 42 °C for 45 s, and immediately placed on ice again for 2 min. Cells were then recovered in 200 µL SOC medium (5 $g{\cdot}L^{-1}$ yeast extract, 20 $g{\cdot}L^{-1}$ tryptone, 10 mM NaCl, 2.5 mM KCl, 10 mM $MgCl_2$, 10 mM $MgSO_4$, 20 mM glucose) for at least 1.5 h before plating on LB agar containing 50 $\mu g{\cdot}mL^{-1}$ kanamycin.

Transformation of *S. oneidensis* and *B. subtilis* was carried out by electroporation. In both cases, 100 ng plasmid DNA was mixed with competent cells and transferred into chilled 1 mm electroporation cuvettes. *S. oneidensis* was electroporated at 1.2 kV, while *B. subtilis* cells were pulsed at 2.0 kV (Eporator, Eppendorf, Germany). Following electroporation, *S. oneidensis* cells were recovered in 200 µL SOC medium, and *B. subtilis* cells in LBMS medium (10 $g{\cdot}L^{-1}$ NaCl, 10 $g{\cdot}L^{-1}$ tryptone, 5 $g{\cdot}L^{-1}$ yeast extract, 0.5 M mannitol, 0.38 M sorbitol) for 3 h. Cells were subsequently plated on LB agar supplemented with kanamycin (50 $\mu g{\cdot}mL^{-1}$ for *S. oneidensis* and 20 $\mu g{\cdot}mL^{-1}$ for *B. subtilis*).

For cultivation, *E. coli* strains harboring the RGB-S plasmid were grown in LB-medium (5 $g{\cdot}L^{-1}$ yeast extract, 10 $g{\cdot}L^{-1}$ tryptone, 10 $g{\cdot}L^{-1}$ NaCl) supplemented with 50 $\mu g{\cdot}mL^{-1}$ kanamycin at 37 °C. In all other experiments, kanamycin was used at a concentration of 40 $\mu g{\cdot}mL^{-1}$, and cultures were additionally supplemented with 1 mM arabinose and 100 µM IPTG for induction and cultivated at 30 °C. For all experiments, freshly transformed cells were used. Overnight precultures were prepared and subsequently diluted to an optical density ($OD_{600}$) of 0.2 prior to inoculation of the respective experiments.

###### 1.5.1. Cultivation in liquid culture

Liquid cultivations were performed in 96-well microtiter plates using a working volume of 100 µL per well at 30 °C under orbital shaking with a 3 mm shaking amplitude in a Synergy plate reader (BioTek, USA). Inoculation was performed with 20 µL of a bacterial suspension ($OD_{600}$ 0.2). For quantitative comparison, fluorescence intensities were normalized to the highest fluorescence value measured within the respective experiment or to respective monoculture for the fluorescent consortia.

###### 1.5.2. Cultivation in MESIF chips

MESIF chips were saturated with 1 mL growth medium via a syringe and inoculated with 20 µL of a bacterial suspension ($OD_{600}$ 0.2). Chips were incubated in sealed Petri dishes containing $ddH_2O$ to maintain humidity and prevent evaporation.

Fluorescence images were acquired daily (Table S3) and selected samples were additionally analyzed by CLSM at defined time points. To facilitate comparison between material conditions and time points, fluorescence intensities were normalized to the maximum fluorescence intensity observed for the respective reporter within an experiment or to respective monoculture for the fluorescent consortia.

##### 1.6. Transcriptomic analysis

###### 1.6.1. RNA extraction and library preparation

For transcriptomic analysis of fluorescent consortia, RNA was extracted 7 days post inoculation using the Quick-RNA™ Fungal/Bacterial MiniPrep Kit (Zymo Research, USA) according to the manufacturer's protocol with minor adjustments. MESIF were halved; one half was processed by suspending it in 800 µL RNA lysis buffer and mechanically compressed with sterile forceps, while the other half was stored at -80 °C for future use. Mechanical disruption was carried out by bead-beating (one cycle of 1 min at 5.0 $m{\cdot}s^{-1}$; MP Biomedicals, USA). Following extraction, 1 µL RNasin® Ribonuclease Inhibitor (Promega, USA) was added, and samples were stored at −80 °C. rRNA depletion was performed using the NEBNext rRNA Depletion Kit (Bacteria) (New England

Biolabs, USA). RNA quality and depletion efficiency were verified using a Bioanalyzer (RNA 600 Nano Kit, Agilent, USA). Library preparation was carried out with the NEBNext Ultra II Directional RNA Library Prep Kit (Illumina, New England Biolabs, USA) using 13 PCR cycles, and barcoding was performed with NEBNext Multiplex Oligos for Illumina. Library size and quality were assessed using the Bioanalyzer High Sensitivity DNA Kit (Agilent, USA), and concentrations were adjusted to 2 nM using the Qubit™ dsDNA HS Assay Kit (Thermo Fisher Scientific, Germany). Libraries were pooled and sequenced on an Illumina NextSeq 1000 platform using a P2 kit (300 cycles).

###### 1.6.2. Data analysis

Raw sequencing reads were quality-filtered and adapter-trimmed using fastp,[11] followed by read merging with FLASH.[12] Residual rRNA was removed using SortMeRNA.[13] Filtered reads were mapped to a combined reference genome using BBMap,[14] comprising concatenated genome sequences and corresponding annotations (FASTA/GFF) obtained from NCBI[15] (*S. oneidensis*: GCF_000146165.2; *B. subtilis*: GCF_000009045.1; *E. coli*: GCF_000005845.2). Resulting alignments were filtered using SAMtools for mapping quality (MAPQ ≥ 30), retaining only primary alignments.[16] Gene-level counts, defined as the raw counts of the corresponding sequencing gene, were generated using featureCounts with fractional assignment of multi-mapping reads.[17] Functional annotation was performed using eggNOG-mapper,[18] and pathway information was retrieved from KEGG via the KEGGREST package.[19] For analyses based on Clusters of Orthologous Groups (COG) functional categories, genes were assigned to all annotated COG categories. Consequently, genes associated with multiple COG categories contributed to each corresponding functional category. Data was analyzed in R (v4.4.2). Differential expression analysis was conducted separately for each organism using DESeq2,[20] applying a generalized linear model (~material) with the native material as the reference level. Genes with low counts (<10 total reads) were excluded. Likelihood ratio tests (LRT) against a reduced model (~1) were used to identify genes significantly affected by experimental conditions and significance was assessed using FDR-adjusted p-values. To compare the relative contribution of individual consortium members to the overall metatranscriptome, transcript abundances were normalized separately for each species. Gene counts were first normalized by gene length and subsequently converted to transcripts per million (TPM) using species specific scaling factors. For visualization of global transcriptional responses, read counts were normalized using DESeq2 variance stabilizing transformation (VST). Genes were assigned to B. subtilis, S. oneidensis, or E. coli based on species specific gene identifiers. Principal component analyses were performed separately for each organism to avoid clustering driven by interspecies transcriptomic differences. For each species, the 500 genes exhibiting the highest variance across all samples were selected, and PCA was calculated using the transformed expression values.

##### 1.7. Environmental sampling

Two experimental approaches were applied: a short-term in situ incubation conducted in a wastewater treatment plant (hereafter "WWTP"), and a long-term incubation performed under controlled laboratory mesocosm conditions (hereafter "mesocosm").

###### 1.7.1. WWTP cultivation

MESIF chips were fabricated as described above (Section S1.2 and S1.3).[1] Prior to incubation, chips were loaded with autoclaved and sterile filtered wastewater, taken from the tank one day prior inoculation. For in situ incubation, the chips were placed inside perforated Falcon tubes (nine openings distributed along the tube body and five in the cap) to facilitate exchange with the surrounding wastewater. One Falcon tube contained one replica of each material environment.

The tubes were secured within a protective cage and submerged in the chemical wastewater basin WP06 B01.1 at KIT Campus North (49.096683, 8.432755). The incubation period lasted 21 days, with samples collected at four time points (days 1, 3, 7 and 21.

###### 1.7.2. Mesocosm cultivation

For long-term sampling, a laboratory-scale mesocosm system was established to enable continuous cultivation of MESIF chips for up to 100 days. MESIF chips were prepared and supplemented as described above.[1] The mesocosm consisted of a 60 L container filled with 40 L of chemical wastewater obtained from the same WWTP tank used for the in situ experiments. The mesocosm was maintained at room temperature under constant stirring (180 rpm) and covered with plastic foil to reduce evaporation. One replica of a MESIF chip of each material environment was were placed in a mesh bag. Samples were collected at five time points (days 1, 3, 7, 21 and 100).

##### 1.8. DNA extraction and library preparation

Genomic DNA (gDNA) was extracted using the DNeasy PowerSoil Pro Kit (QIAGEN, Germany) according to the manufacturer's protocol with minor adjustments. For MESIF samples were divided into two halves. One half was immediately stored at −20 °C for future use, while the other half was processed by mechanically compressing the material in CD1 lysis buffer to release biomass embedded within the matrix. For control 2 mL liquid wastewater samples were centrifuged (10,000 × g, 10 min) and the supernatant discarded. The resulting pellet was resuspended in CD1 buffer and processed following the standard extraction procedure. Mechanical disruption was carried out by bead-beating (two cycles of 1 min at 5.0 $m \cdot s^{-1}$; MP Biomedicals, USA). DNA quantity was assessed fluorometrically using the Qubit™ dsDNA HS Assay Kit (Thermo Fisher Scientific, Germany). The gDNA samples were stored at -20 °C until further use.

Microbial community composition was assessed using 16S rRNA gene amplicon sequencing and shotgun metagenomics.

For amplicon sequencing, the prokaryotic 16S rRNA V3 region was targeted using the degenerate primers 341bf (ACACTCTTTCCCTACACGACGCTCTTCCGATCTCCTACGGGNGGCWGCAG) and 518r (GACTGGAGTTCAGACGTGTGCTCTTCCGATCTWTTACCGCRGCTGCTGG), which included Illumina TruSeq adapter sequences. PCR amplification was performed in 25 µL reactions containing 0.04 $U \cdot \mu L^{-1}$ Q5 High-Fidelity DNA Polymerase, 1× Q5 reaction buffer, 1× high GC enhancer, 0.25 µM of each primer, 0.2 mM dNTPs, and 100 ng template DNA. Amplification followed a touchdown protocol (95 °C, 2 min; 4 cycles of 95 °C for 30 s, annealing from 56–52 °C for 40 s, 72 °C for 40 s; 20 cycles of 95 °C for 30 s, 51 °C for 40 s, 72 °C for 40 s; final extension at 72 °C for 5 min). PCR products were purified using a Clean & Concentrate-5 Kit (Zymo Research, Germany) according to the manufacturer's instructions.

Sequencing libraries were prepared from purified genomic DNA (shotgun metagenomics) and amplicon PCR products using the NEBNext Ultra II FS DNA Library Prep Kit (New England Biolabs, Germany) according to the manufacturer's instructions for DNA inputs ≥100 ng. For shotgun libraries, DNA was enzymatically fragmented for 14 min, whereas amplicon libraries were prepared without additional fragmentation. Adapter ligation and indexing were performed using NEBNext Multiplex Oligos for Illumina with six PCR cycles. Library fragment size and quality were assessed using a Bioanalyzer (High Sensitivity DNA Kit, Agilent, USA), and concentrations were normalized to 2 nM using the Qubit™ dsDNA HS Assay Kit (Thermo Fisher Scientific, Germany). Libraries were pooled and subjected to paired-end sequencing on an Illumina NextSeq 1000 platform using a P2 kit (300 cycles).

##### 1.9. qPCR DNA hydrogel quantification

Quantification of RCA-derived DNA hydrogel material was performed by quantitative PCR (qPCR) following a previously established protocol.[21] The primer and probe sequences used are listed in Table S1. Briefly, calibration curves were generated for each linear DNA template using standard solutions ranging from 156.3 pM to 20 nM. Prior to qPCR analysis, samples and standards were thoroughly mixed and subjected to 5 s of ultrasonic treatment to fragment the high-molecular-weight DNA concatemers generated during rolling circle amplification. For qPCR reactions, a master mix was prepared containing 1× qPCR buffer (16 mM $(NH_4)_2SO_4$, 67 mM Tris-HCl, 2.5 mM KCl, 2.5 mM $MgCl_2$, and 0.01% (v/v) Tween-20), 200 µM of each dNTP, 1 µM forward primer, 1 µM reverse primer, 0.2 µM hydrolysis probe and Taq DNA polymerase (50 $U \cdot mL^{-1}$; New England Biolabs, USA). Reactions were assembled by combining 20 µL of the qPCR master mix with 1.5 µL of sample or calibration standard. All measurements were performed in technical triplicate using a real-time PCR system (Bio-Rad, USA). DNA concentrations were determined from the corresponding calibration curves by linear regression of cycle threshold (Ct) values against the logarithm of the standard concentrations. Hydrogel-derived DNA was quantified using a hydrogel-sequence-specific qPCR assay. Concentrations obtained from the standard curve were corrected for sample dilution, converted to mass concentrations using the molecular weight of the target amplicon, and expressed as a percentage of the total DNA concentration measured in the corresponding DNA extract.

##### 1.10. Data analysis for amplicon data

Amplicon sequencing data were processed using QIIME 2 (version 2025.10.1).[22] Raw paired-end reads were first quality-filtered and adapter-trimmed using fastp,[11] followed by read merging with FLASH to obtain full-length amplicon sequences.[12] The merged reads were subsequently imported into QIIME 2, where residual adapter and primer sequences were removed using cutadapt.[23] Denoising and inference of amplicon sequence variants (ASVs) were performed using the DADA2 algorithm.[24] Chimeric sequences were identified and removed using VSEARCH.[25] Representative ASV sequences were aligned with MAFFT via the q2-alignment plugin, and phylogenetic trees were generated using FastTree2 through the q2-phylogeny plugin.[26] Taxonomic assignment of ASVs was conducted within QIIME 2 using Naïve Bayes classifiers trained on the SILVA reference database (release 138.2).[27,28]

Alpha and beta diversity analyses were performed in R (v4.4.2) using the qiime2R,[29] phyloseq,[30] and microbiome packages.[31] Alpha diversity was assessed using Shannon diversity. Beta diversity was evaluated based on Bray–Curtis dissimilarities with PERMANOVA using the vegan package.

Analyses of material effects were restricted to the seven replicated conditions (native, APTES, GPTSh, PEG, catalase, ODN, and DNA hydrogel). Trajectories visualized as Principal component analysis (PCA) was performed on transformed ASV count tables using a centered log-ratio (CLR) and subsequent community dissimilarity quantified as Euclidean distance in CLR space, corresponding to Aitchison distance. The contribution of material environments to variation in community composition was quantified separately for each timepoint using permutational multivariate analysis of variance (PERMANOVA) based on Aitchison distances using the vegan package. [32] and visualized as $R^2$. Homogeneity of multivariate dispersion among materials was assessed separately at each sampling time using distance-to-centroid analysis (betadisper) followed by permutation tests. To assess temporal changes in the magnitude and configuration material specific centroids were calculated directly in CLR and pairwise Aitchison distances were calculated to characterize the overall magnitude of compositional separation. Temporal preservation of the relative configuration of material specific communities was assessed by comparing matrices of pairwise habitat-centroid distances between sampling times using Spearman Mantel correlations. Statistical significance was assessed by permutation of material environment labels.

##### 1.11. Data analysis for shotgun metagenomic data

Raw sequencing reads were subjected to quality control and adapter removal using fastp.[11] Filtered reads were subsequently merged with FLASH.[12] Metagenomic assemblies were generated using MEGAHIT.[33] To quantify contig abundance, quality-filtered reads were aligned back to their respective assemblies with CoverM,[34] and resulting alignment files were further processed using samtools.[16] The resulting coverage estimates were subsequently used for taxonomic, functional, and genome-resolved downstream analyses.

Taxonomic characterization of microbial communities was carried out by recovering phylogenetic marker genes from assembled contigs using MDMcleaner.[35] Extracted small subunit and large subunit rRNA gene sequences were classified taxonomically with SINA against the SILVA database.[36,37] Relative abundance profiles were generated by mapping reads back to the identified marker genes using CoverM.

For functional characterization, open reading frames (ORFs) were predicted from assembled contigs with Prodigal.[38] Functional annotation was then conducted in metagenomic mode using eggNOG-mapper.[18] Annotation outputs were integrated with contig coverage information to produce abundance-weighted functional profiles, including COG categories,[39] KEGG orthologs,[40–42] KEGG pathways and modules, as well as Pfam domains.[43] These datasets were subsequently used to evaluate the metagenomic potential of the microbial communities. To investigate the distribution of selected ecological and physiological functions across material environments, genes were assigned to manually curated functional categories representing biofilm formation, quorum sensing, motility, c-di-GMP signaling, competence, DNA processing and metabolism, mobile genetic elements, developmental processes, persistence, stress responses, respiration and metabolic functions. Functional assignments were based on combinations of gene identifiers, annotation keywords and Pfam domain annotations retrieved from eggNOG-mapper outputs. As individual genes can possess multiple functions or annotation terms, a single gene could contribute to more than one functional category when it fulfilled the respective assignment criteria. Functional categories therefore represent overlapping functional profiles rather than mutually exclusive gene sets.

Genome-resolved metagenomic analysis employed a multi-binning workflow integrating MaxBin2,[44] MetaBAT2,[45] and CONCOCT,[46] followed by bin refinement using DAS Tool.[47] Binning procedures were applied to individual assemblies using coverage-guided approaches. Recovered genome bins were dereplicated with dRep,[48] at 95% average nucleotide identity (ANI; fastANI), retaining genomes with ≥50% completeness and ≤5% contamination and curated using MDMcleaner[35]. Genome quality was assessed with CheckM2[49] and taxonomic assignments were obtained using GTDB Tk. [50] MAG abundance estimates were obtained through read mapping using CoverM.

##### 1.12. Detection and characterization of candidate DNA hydrogel–microbial DNA junctions

To identify candidate sequence junctions between DNA hydrogel template sequences and microbial genomic DNA, paired-end sequencing reads were screened for chimeric molecules containing both DNA hydrogel template and microbial sequences. Reads were aligned against a DNA hydrogel template reference and the corresponding sample-specific microbial assembly using Bowtie2 in very-sensitive-local mode.[51] Candidate chimeric reads were required to contain at least 25 bp matching the DNA hydrogel template and at least 25 bp matching microbial genomic DNA (gDNA), with the microbial alignment exhibiting a mapping quality (MAPQ) ≥ 20. Candidate reads were retained only when their paired mate also had MAPQ ≥ 20, both reads mapped to the same microbial contig, the pair was properly paired, its orientation matched the predominant library orientation, and its template span fell within the empirical 1st-99th percentile range. For downstream analysis, only reads in which the DNA hydrogel template and microbial segments were directly adjacent or overlapping were considered. Junction breakpoints were inferred at the boundary between the microbial and DNA hydrogel template alignments, or, in the case of overlap, right at the beginning of the overlapping region. Candidate junctions were subsequently rescreened across all paired-end FASTQ datasets using Bowtie2 in very-sensitive-local mode with reporting of all alignments (--very-sensitive-local -a). Supporting reads were required to span the inferred breakpoint with at least 25 aligned bases on each side and to exhibit 100% sequence identity across the resulting 50 bp junction-spanning region. PCR duplicates were removed by collapsing reads with identical sequences that mapped to the same coordinates on the reference contig. The number of remaining independent supporting read pairs was then determined and used to assess junction occurrence across the environmental datasets. The microbial-mapping portions of candidate junction-supporting reads were assigned to assembled metagenomic contigs and annotated with Bakta,[52] eggNOG-mapper and BLASTp to characterize the genomic context surrounding candidate DNA hydrogel–microbial DNA junctions. Genes located within ±2 kb of the inferred microbial breakpoint were manually assigned to broad functional categories. Mobile-element- or phage-associated genes were identified by keyword-based screening of functional annotations. Genes annotated as transposases, integrases, recombinases, phage- or prophage-associated proteins, virion proteins, terminases, tail-associated proteins, endolysins, homing endonucleases or Cro repressors were classified as mobile-element- or phage-associated. Further categories included DNA replication/recombination, transport, regulation/signaling, cell surface/secretion, metabolism/other, and hypothetical/unknown functions. Genes were assigned to all applicable functional categories based on their annotations; consequently, individual genes could contribute to multiple categories and category counts are not mutually exclusive.

##### 1.13. Data and statistical Analysis

Data were expressed as means ± standard deviation, with at least three biological replicates per condition ($n \geq 3$). Log transformation was performed as $\log_2(x + 0.001)$ or $\log_{10}(x + 0.01)$to avoid undefined values for zero measurements.

Statistical analyses were performed in R (v4.4.2). Fluorescence trajectories were analyzed using linear mixed effects models implemented in the *lme4* package in R. [53] Material environment, cultivation time, and their interaction were included as fixed effects, while individual MESIF chips were treated as a random effect to account for repeated measurements of the same biological replicate over time. Significance of fixed effects was assessed using Type III ANOVA with Satterthwaite approximation of degrees of freedom as implemented in the lmerTest package. [54] Pairwise comparisons between material conditions were performed using estimated marginal means with Benjamini-Hochberg correction for multiple testing. A significance threshold of $p < 0.05$ was applied throughout.

Correlations were assessed using Spearman's rank correlation coefficient.

##### 1.14. Data availability

Metagenomic and transcriptomic sequencing reads generated in this study have been submitted to RADAR (DOI 10.35097/yhdzt8n0gf4j8p2q).

#### 2. Supplementary Figures

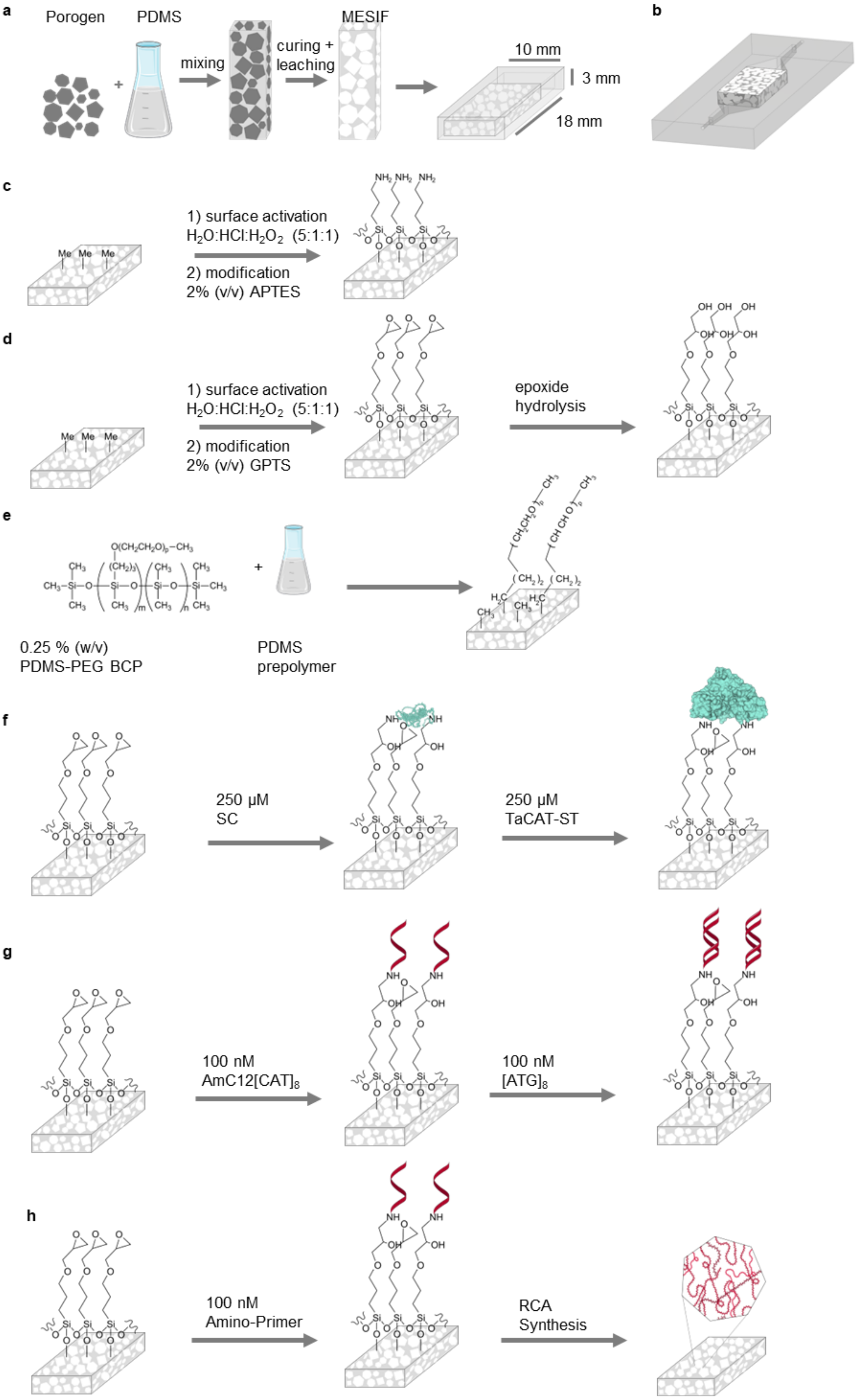


**Fig. S1 | Fabrication and functionalization of MESIF materials. (a)** Fabrication of macroporous elastomeric silicone foam (MESIF) by mixing a sacrificial porogen with polydimethylsiloxane (PDMS) prepolymer, followed by curing and porogen leaching to form an interconnected pore network. **(b)** Integration of the MESIF scaffold into the cultivation device. **(c)** Surface activation and silanization with (3-aminopropyl)triethoxysilane (APTES) to introduce primary amino groups. **(d)** Silanization with (3-glycidyloxypropyl)trimethoxysilane (GPTS), followed by epoxide hydrolysis to generate hydroxyl-functionalized interfaces (GPTSh). **(e)** Incorporation of a PDMS–poly(ethylene glycol) (PEG) block copolymer into the PDMS prepolymer to generate PEG-functionalized MESIF. **(f)** Immobilization of SpyCatcher (SC) on GPTS-functionalized MESIF, followed by covalent capture of SpyTag-functionalized catalase (TaCAT–ST). **(g)** Covalent attachment of amino-modified oligodeoxynucleotides (ODNs) to GPTS-functionalized MESIF, followed by hybridization with complementary DNA strands to generate surface-bound DNA duplexes. **(h)** Immobilization of amino-modified DNA primers on GPTS-functionalized MESIF, followed by rolling-circle amplification (RCA) to generate an extended DNA hydrogel within the MESIF pore network. Schematics are not drawn to scale.

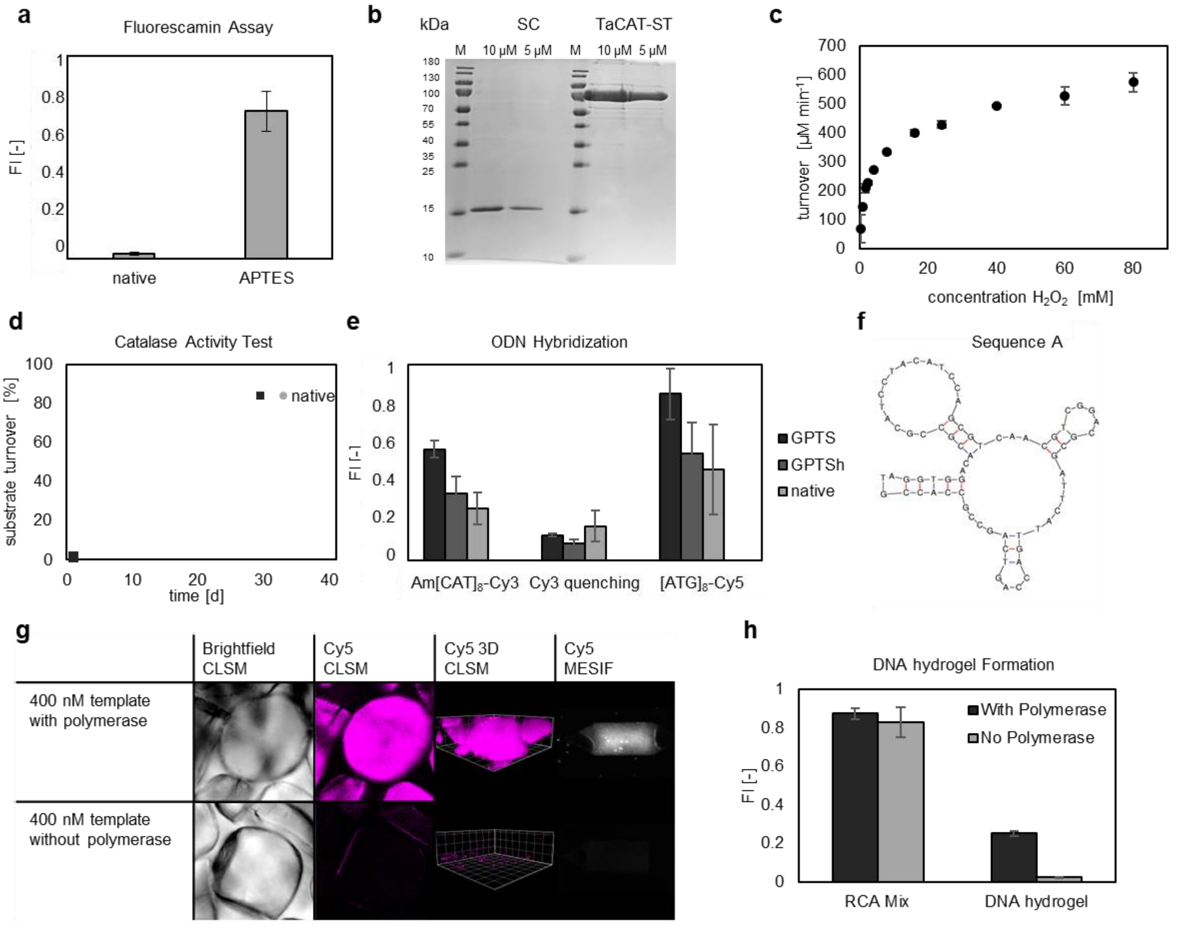


**Fig. S2 | Verification of MESIF functionalization and biomolecular activity.** Unless stated otherwise, experiments were performed in triplicate (n = 3), and quantitative data are presented as mean ± SD. **(a)** Fluorescamine assay comparing native and APTES-functionalized MESIF, with increased fluorescence indicating the presence of accessible primary amino groups following silanization. **(b)** SDS–PAGE analysis of purified SpyCatcher (SC; 15.4 kDa) and SpyTag-functionalized catalase (TaCAT–ST; 84.4 kDa) used for enzyme immobilization. **(c)** Michaelis–Menten analysis of purified TaCAT–ST (62 nM) based on hydrogen peroxide ($H_2O_2$) turnover, yielding $K_M = 5.0 \pm 0.5$ mM and $k_{cat} = 239.0 \pm 9.1\ s^{-1}$. **(d)** Time-dependent $H_2O_2$ conversion by catalase-functionalized MESIF compared with native MESIF controls over 40 days, demonstrating retained enzymatic activity following immobilization. **(e)** Fluorescence-based verification of oligodeoxynucleotide (ODN) immobilization and hybridization on GPTS-, GPTSh- and native MESIF. Cy3 fluorescence from the immobilized amino-modified anchor strand (Am-[CAT]$_8$-Cy3), Cy3 quenching following addition of the complementary strand and Cy5 emission upon Cy3 excitation as a FRET-associated hybridization signal are shown for the indicated surfaces. **(f)** Predicted secondary structure of the 72-nucleotide circular DNA template containing three self-complementary, hairpin-forming domains used for rolling-circle amplification (RCA). **(g)** Representative bright-field and Cy5 fluorescence microscopy images of MESIF following RCA reactions performed with or without DNA polymerase, showing fluorescence-associated DNA structures within individual pores and at the whole-MESIF scale after 72 h of reaction and subsequent rinsing. **(h)** Fluorescence intensities of RCA reaction mixtures containing Cy5-labelled dUTP (0.5 µM) and the resulting MESIF-associated DNA material after polymerization and rinsing, comparing reactions with and without DNA polymerase.

**Extended Description | Fabrication, chemical diversification and biomolecular validation of MESIF (Supplementary Figs. S1–S2)**

The macroporous elastomeric silicone foam (MESIF) platform was designed to provide a common three-dimensional scaffold in which the extracellular material environment could be systematically varied. MESIF was fabricated from polydimethylsiloxane (PDMS) using sacrificial salt crystals as porogens, followed by curing and salt leaching to generate an interconnected pore network (Supplementary Fig. S1a). The resulting scaffolds were integrated into PDMS cultivation devices comprising a medium reservoir and an environmental contact region (Fig. 1a; Supplementary Fig. S1b). This architecture enabled cultivation of defined microbial populations and exposure to environmental microbiomes within a shared porous material platform. Detailed fabrication and assembly procedures are provided in Supplementary Methods Sections 1.2 and 1.3.

Seven material environments were established by modifying the PDMS scaffold or introducing biomolecular components (Fig. 1b; Supplementary Fig. S1c–h). Surface activation followed by silanization with (3-aminopropyl)triethoxysilane (APTES) generated interfaces presenting primary amino groups, whereas treatment with (3-glycidyloxypropyl)trimethoxysilane (GPTS) followed by epoxide hydrolysis produced hydroxyl-functionalized GPTSh interfaces (Supplementary Fig. S1c,d).[2] In contrast, PEG modification was achieved by incorporating a PDMS–PEG block copolymer into the prepolymer before scaffold fabrication (Supplementary Fig. S1e).[3] The presence of accessible amino groups following APTES functionalization was supported by a pronounced increase in fluorescamine-associated fluorescence relative to native MESIF (Supplementary Fig. S2a). Together, these modifications provided chemically distinct extracellular interfaces without changing the underlying MESIF cultivation concept.

The biomolecular environments introduced additional functional and structural diversity. Catalase-functionalized MESIF was prepared by immobilizing SpyCatcher on GPTS-modified surfaces, followed by covalent capture of SpyTag-fused catalase from *Thermoanaerobaculum aquaticum* (TaCAT–ST; Supplementary Fig. S1f).[5] SDS–PAGE confirmed the purified protein components, while hydrogen peroxide turnover established catalytic activity of TaCAT–ST, with a Michaelis constant of 5.0 ± 0.5 mM and a turnover number of 239.0 ± 9.1 $s^{-1}$ (Supplementary Fig. S2b,c). Catalase-functionalized MESIF also exhibited hydrogen peroxide conversion during repeated measurements over 40 days, unlike native controls (Supplementary Fig. S2d). These observations support the presence of a functionally active enzyme-associated material environment, although the resulting local oxygen and peroxide concentrations during microbial cultivation were not directly determined.

Two distinct DNA-containing environments were generated using GPTS-based surface attachment. For ODN-functionalized MESIF, amino-modified oligonucleotides were covalently attached through epoxy–amine coupling and subsequently hybridized with complementary strands, producing surface-associated DNA duplexes (Supplementary Fig. S1g). Fluorescence measurements using labelled oligonucleotides supported immobilization and hybridization through changes in Cy3 fluorescence and a FRET-associated Cy5 signal following complementary-strand addition (Supplementary Fig. S2e). Thus, ODN functionalization presented relatively short DNA sequences at the material interface without requiring polymerase-driven DNA synthesis.

By contrast, DNA hydrogel-functionalized MESIF was prepared by immobilizing amino-modified primers on GPTS surfaces, followed by rolling-circle amplification (RCA) using a circular DNA template and Phi29 DNA polymerase (Supplementary Fig. S1h).[9] The template design included self-complementary domains capable of forming secondary structures within the amplified DNA product (Supplementary Fig. S2f). After 72 h of RCA and subsequent rinsing, Cy5-associated fluorescence was observed within MESIF pores and at the whole-scaffold scale, with substantially stronger retained fluorescence in polymerase-containing reactions than in polymerase-free controls (Supplementary Fig. S2g,h). These results support polymerase-dependent formation and retention of extended DNA material within the porous scaffold. The fluorescence-based characterization does not, by itself, establish the mechanical properties or exact spatial connectivity of the DNA network.

The distinction between the two DNA-containing environments is central to the experimental design. ODN-functionalized MESIF presents short, surface-tethered DNA duplexes, whereas RCA produces an extended DNA-rich material within the pore network (Fig. 1b; Supplementary Fig. S1g,h and S2e–h). These environments therefore differ not only in DNA presentation but also in polymer architecture, local DNA distribution and preparation chemistry. Comparisons between them can reveal whether microbial phenotypes depend on the form in which extracellular DNA is presented, but cannot isolate any single structural or chemical variable without additional controls.

Collectively, the fabrication and validation experiments establish a common porous MESIF platform incorporating chemically and biomolecularly distinct material environments. They provide the material-science basis for the subsequent analyses of growth-associated fluorescence, stress-reporter activity, species-specific transcriptional states and microbial community succession (Figs. 2–6), while distinguishing the experimentally verified functionalizations from mechanistic explanations of their biological effects.

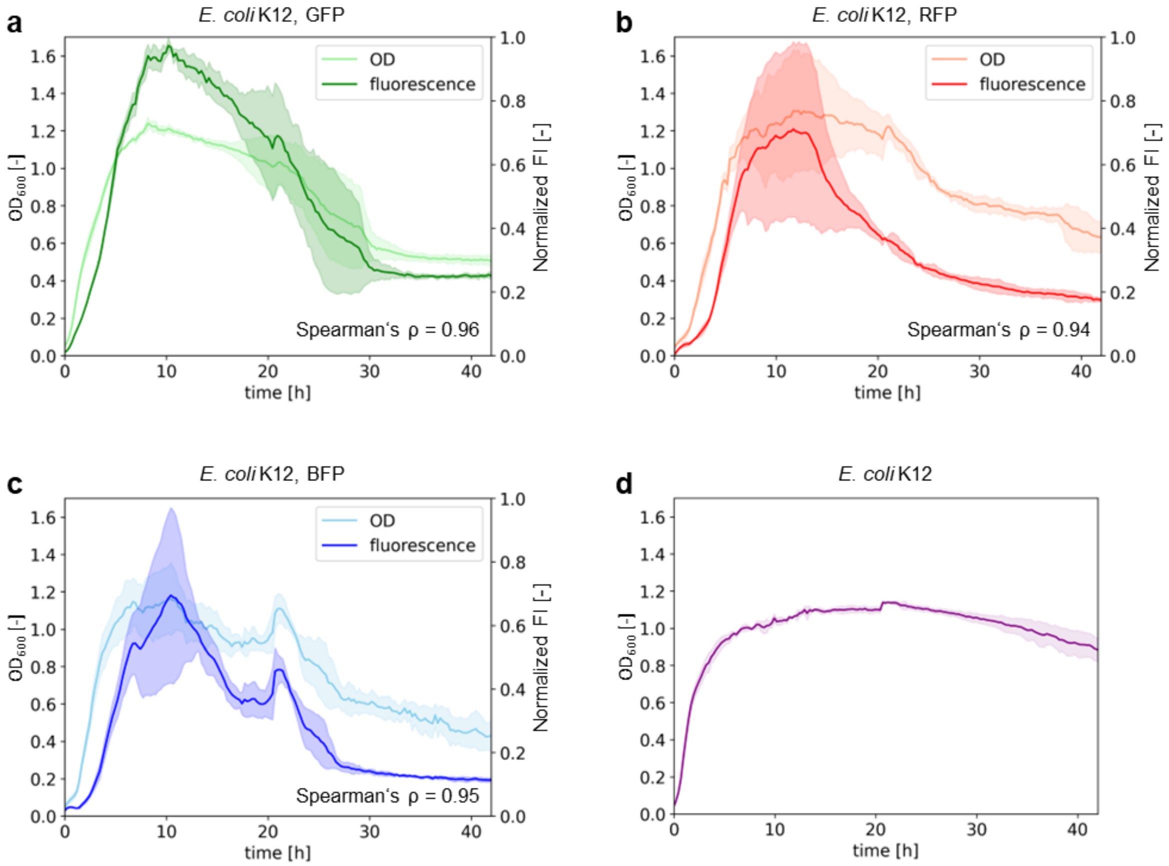


**Fig. S3 | Growth and fluorescence dynamics of fluorescent *E. coli* reporter strains in liquid culture.** *Escherichia coli* K-12 MG1655 strains expressing GFP, RFP or BFP were cultivated in liquid culture at 30 °C. **(a–c)** Time-resolved optical density at 600 nm ($OD_{600}$; light-coloured curves, left axis) and normalized fluorescence intensity (FI; dark-coloured curves, right axis) for GFP-expressing (a), RFP-expressing (b) and BFP-expressing (c) strains. Spearman correlation coefficients between $OD_{600}$ and fluorescence intensity were ρ = 0.96, 0.94 and 0.95, respectively. Fluorescence intensities were normalized to the maximum value measured for each reporter strain. **(d)** Growth curve of plasmid-free *E. coli* K-12 MG1655 cultivated without kanamycin. Lines represent means and shaded areas indicate standard deviations from n = 3 independent liquid cultures.

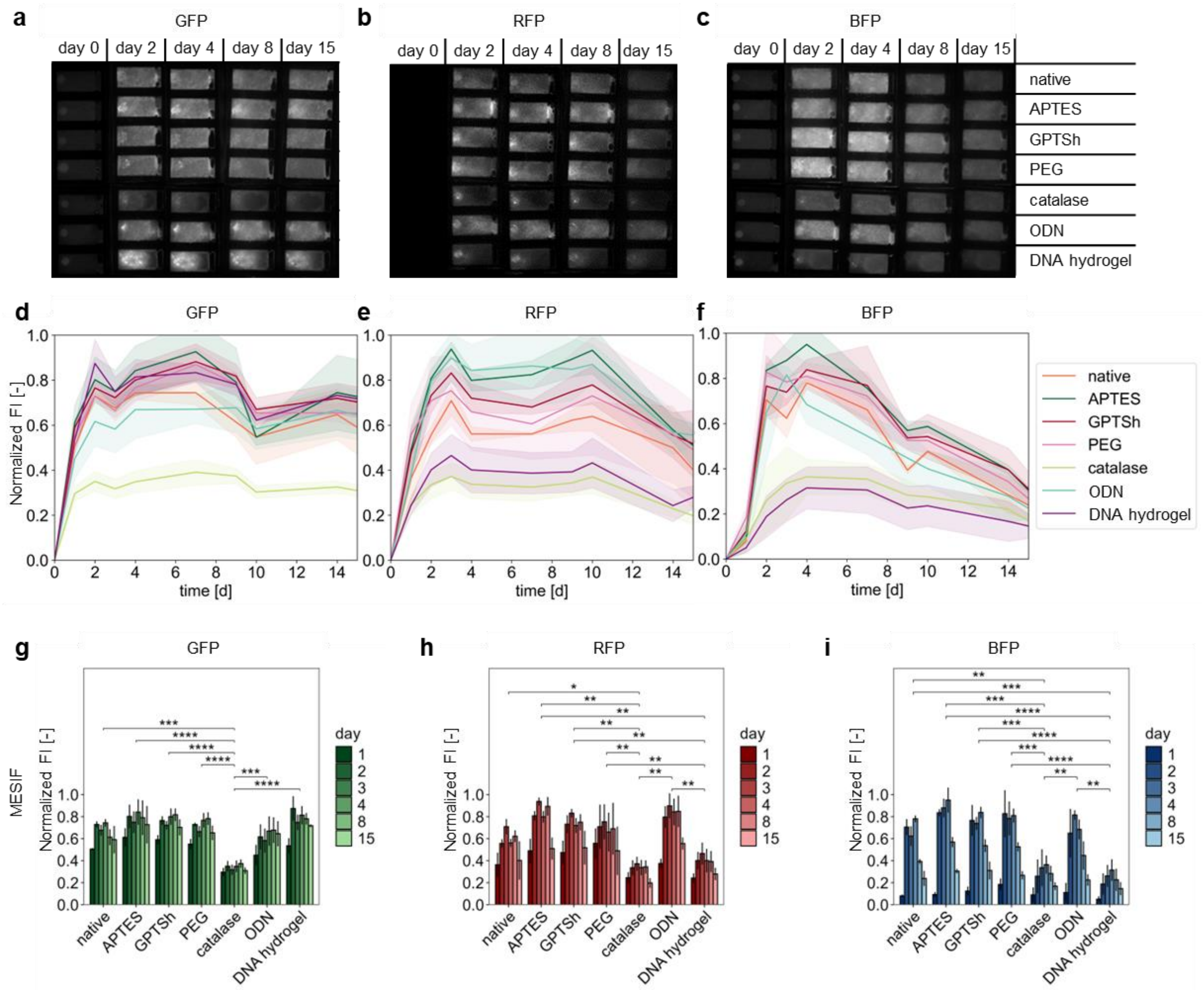


**Fig. S4 | Material-dependent fluorescence dynamics of *E. coli* reporter strains during MESIF colonization.** *Escherichia coli* K-12 MG1655 strains expressing GFP, RFP or BFP were cultivated for 15 days at 30 °C in native, APTES-, GPTSh-, PEG-, catalase-, ODN- or DNA hydrogel-functionalized MESIF. **(a–c)** Representative whole-MESIF fluorescence images acquired on days 0, 2, 4, 8 and 15 for GFP-expressing (a), RFP-expressing (b) and BFP-expressing (c) strains. **(d–f)** Longitudinal whole-MESIF fluorescence intensities (FI) for GFP (d), RFP (e) and BFP (f) across the seven material environments. Lines indicate means and shaded areas represent standard deviations. **(g–i)** Fluorescence intensities resolved by material environments and cultivation day for GFP (g), RFP (h) and BFP (i). Bars indicate means and error bars represent standard deviations. All quantitative data were obtained from n = 3 independent MESIF chips per material condition, repeatedly measured over the cultivation period. Fluorescence intensities were normalized to the maximum value measured for the respective reporter strain. Statistical significance was assessed using linear mixed-effects models with material condition, cultivation time and their interaction as fixed effects and MESIF chip as a random effect. Pairwise comparisons between material conditions were performed using estimated marginal means with Benjamini–Hochberg correction for multiple testing. Significance levels are indicated as $p < 0.05$ (*), $p < 0.01$ (**), $p < 0.001$ (***), and $p < 0.0001$ (****).

**Extended Description | Material-dependent growth-associated fluorescence of *E. coli* in MESIF (Supplementary Figs. S3–S4)**

To establish fluorescence as a readout for monitoring bacterial colonization of MESIF, we first characterized *Escherichia coli* K-12 MG1655 strains expressing GFP, RFP or BFP in liquid culture at 30 °C. Fluorescence intensity correlated strongly with optical density for all three strains, with Spearman correlation coefficients of $\rho = 0.96$, 0.94 and 0.95, respectively (Supplementary Fig. S3a–c). However, the fluorescence and optical-density trajectories were not identical, and their temporal relationships differed among reporters. The growth profile of plasmid-free *E. coli* provided an additional reference for cultivation under these conditions (Supplementary Fig. S3d). Together, these observations support the use of reporter fluorescence as a growth-associated readout, while emphasizing that fluorescence cannot be interpreted as a direct or universally proportional measure of cell number or biomass.

When transferred to MESIF, the three reporter strains exhibited distinct material-dependent fluorescence profiles over 15 days of cultivation (Fig. 2a–c; Supplementary Fig. S4a–i). The complete longitudinal measurements revealed an initial increase in fluorescence followed by reporter- and material-specific trajectories (Supplementary Fig. S4d–f), whereas the corresponding time-resolved comparisons documented differences among the seven environments throughout cultivation (Supplementary Fig. S4g–i). Catalase-functionalized MESIF was associated with low fluorescence across all three strains. In contrast, the DNA hydrogel environment produced divergent reporter-specific responses. GFP-associated fluorescence remained relatively prominent, whereas RFP- and BFP-associated fluorescence were comparatively weak (Fig. 2a–c; Supplementary Fig. S4g–i). These differences caution against interpreting any single fluorescent reporter as a universal measure of bacterial growth across chemically distinct extracellular environments.

Three-dimensional confocal microscopy additionally revealed material-dependent distributions of fluorescent *E. coli* within individual MESIF pores (Fig. 2d). Fluorescence-associated structures varied across material conditions and reporter strains, providing spatial context for the whole-MESIF measurements. However, these reconstructions are representative observations rather than direct quantitative measurements of viable biomass or pore occupancy. The observed differences may reflect a combination of bacterial colonization, reporter expression and fluorescence-related properties, which cannot be separated by these measurements alone.

These considerations are particularly important for interpreting the subsequent physiological experiments. The RGB-S system combines SOS-, RpoS- and RpoH-responsive fluorescent modules to monitor stress-associated transcriptional activity (Supplementary Fig. S5a,b).[10] Because the single-reporter strains themselves exhibit material-dependent fluorescence, absolute RGB-S signals cannot be attributed exclusively to changes in stress-responsive promoter activity. Accordingly, the RGB-S measurements were evaluated both as longitudinal fluorescence profiles (Fig. 3a–c; Supplementary Fig. S6d–f) and relative to the corresponding single-reporter signals under matched material and cultivation-time conditions (Fig. 3d–f). This comparison provides a more informative basis for identifying material-associated physiological reporter states, while retaining the limitations inherent to fluorescence-based measurements.

Together, the single-reporter experiments establish a fluorescence-based framework for assessing *E. coli* responses within MESIF. The contrasting GFP, RFP and BFP profiles under DNA hydrogel conditions are especially informative: they demonstrate that this extracellular environment does not simply increase or suppress all fluorescent signals uniformly, motivating the subsequent investigation of stress-associated responses within the same material environment.

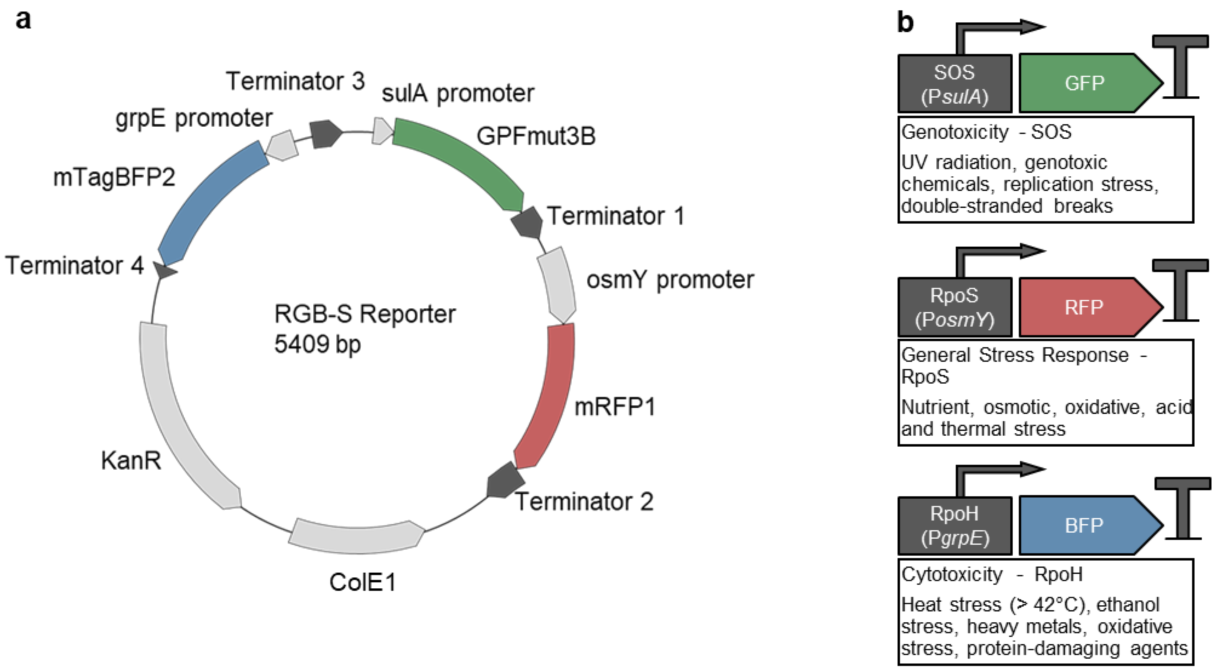


**Fig. S5 | Genetic architecture and stress-response modules of the RGB-S reporter plasmid. (a)** Schematic map of the 5,409-bp RGB-S reporter plasmid comprising three stress-responsive transcriptional reporter modules, a kanamycin-resistance marker (KanR) and a ColE1 origin of replication. **(b)** Schematic representation of the three promoter–reporter modules: the SOS-responsive *sulA* promoter (PsulA) driving GFPmut3B expression, the RpoS-dependent *osmY* promoter (PosmY) driving mRFP1 expression, and the RpoH-dependent *grpE* promoter (PgrpE) driving mTagBFP2 expression. These modules enable fluorescence-based monitoring of SOS-associated genotoxic stress, RpoS-mediated general stress and RpoH-associated proteotoxic stress responses, respectively. Adapted from Zoheir *et al.*[10]

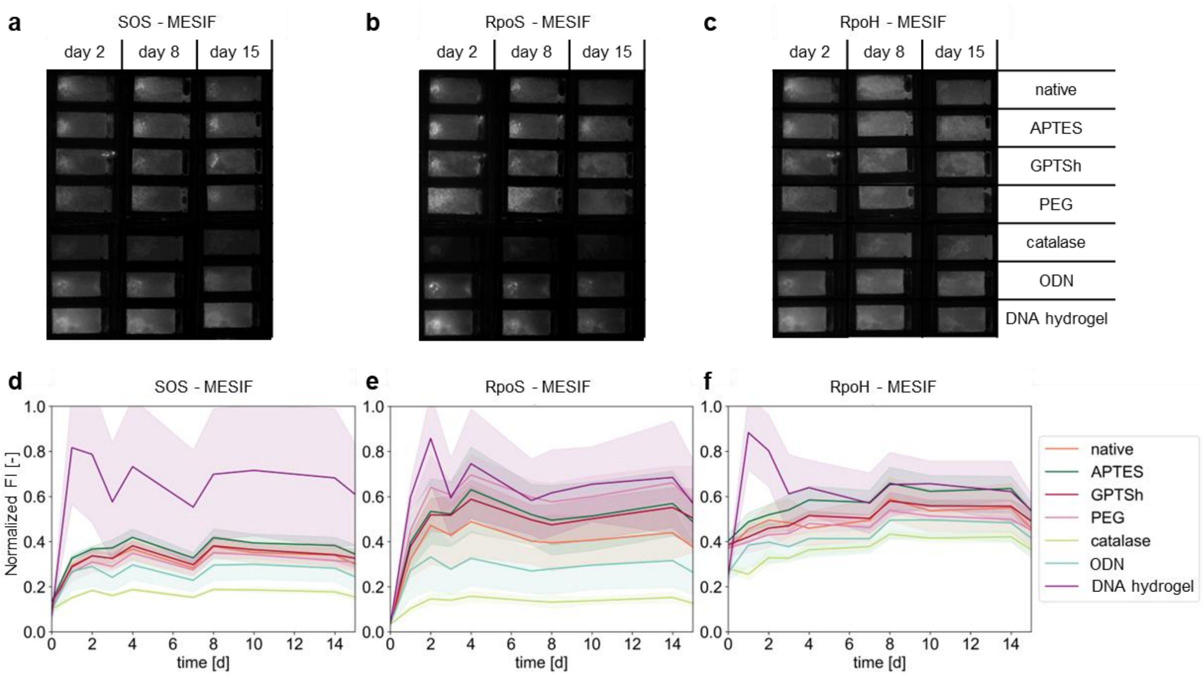


**Fig. S6 | Temporal dynamics of RGB-S stress-reporter fluorescence across MESIF material environments.** *Escherichia coli* K-12 MG1655 carrying the RGB-S reporter plasmid was cultivated for 15 days in native, APTES-, GPTSh-, PEG-, catalase-, ODN- or DNA hydrogel-functionalized MESIF. **(a–c)** Representative whole-MESIF fluorescence images acquired on days 2, 8 and 15 for the SOS-associated GFP (a), RpoS-associated RFP (b) and RpoH-associated BFP (c) reporter channels. **(d–f)** Corresponding longitudinal whole-MESIF fluorescence intensities (FI) for the SOS (d), RpoS (e) and RpoH (f) reporter channels across the seven material environments. Lines indicate means and shaded areas represent standard deviations from n = 3 independent MESIF chips per material condition.

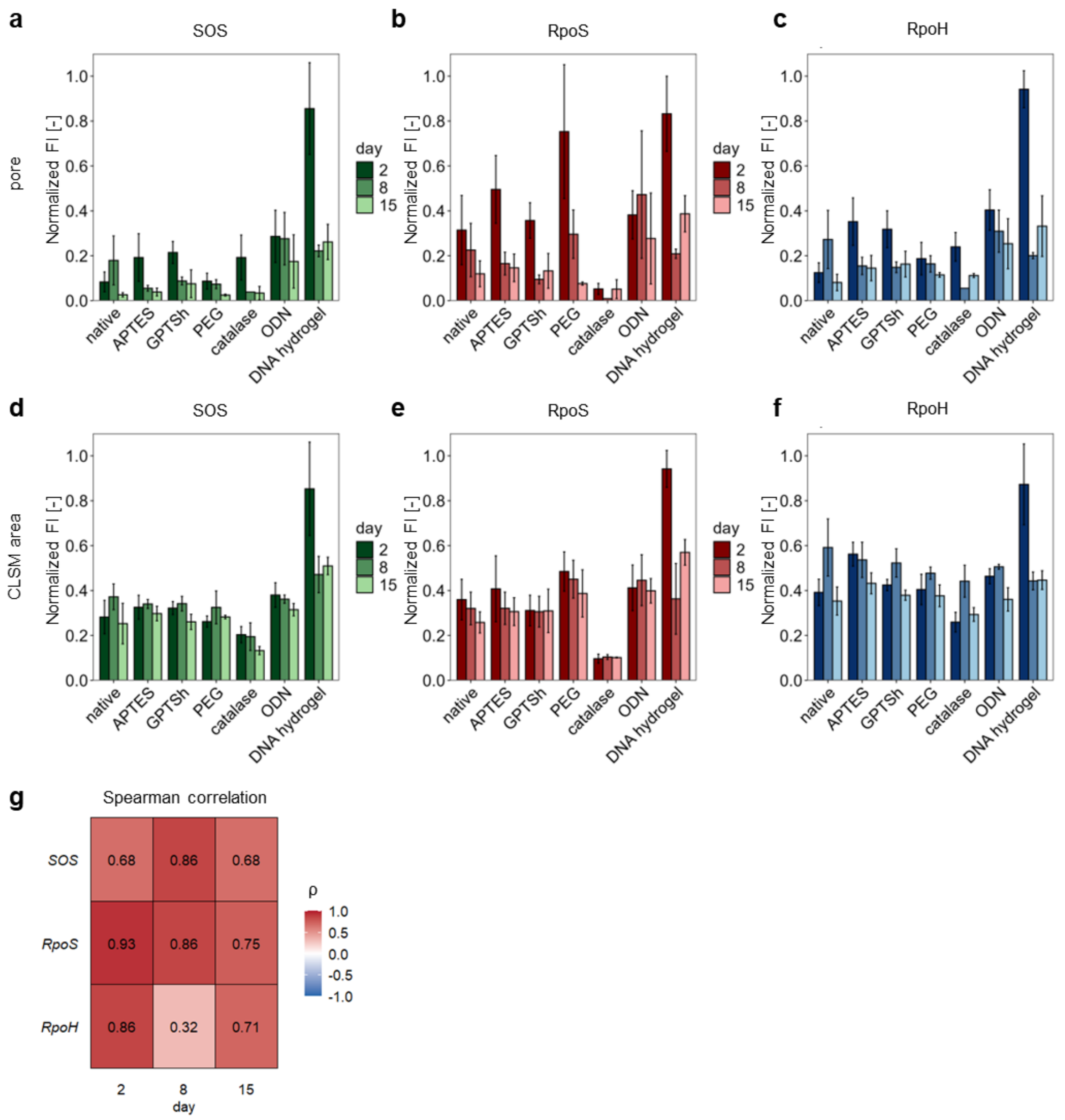


**Fig. S7 | Comparison of CLSM- and whole-MESIF-based quantification of RGB-S stress-reporter fluorescence.** *Escherichia coli* K-12 MG1655 carrying the RGB-S reporter plasmid was cultivated in native, APTES-, GPTSh-, PEG-, catalase-, ODN- or DNA hydrogel-functionalized MESIF. The reporter comprises three fluorescence channels associated with SOS-mediated genotoxic stress (GFP), RpoS-mediated general stress (RFP) and RpoH-associated proteotoxic stress (BFP). **(a–c)** Fluorescence intensities (FI) quantified from confocal laser scanning microscopy (CLSM) images of individual MESIF pores on days 2, 8 and 15 for the SOS (a), RpoS (b) and RpoH (c) reporter channels. Bars indicate means and error bars represent standard deviations across multiple individual pore images acquired from a single representative MESIF chip per material condition and time point. **(d–f)** Corresponding fluorescence intensities quantified from regions of interest in whole-MESIF fluorescence images matched to the pore regions examined by CLSM, for SOS (d), RpoS (e) and RpoH (f). Bars indicate means and error bars represent standard deviations from n = 3 independent MESIF chips per material condition. No inferential statistical comparisons between material conditions were performed for the CLSM- or ROI-based bar-chart measurements owing to limitations of the imaging replication and longitudinal analysis. **(g)** Spearman correlation coefficients (ρ) comparing CLSM-derived and ROI-based whole-MESIF fluorescence intensities separately for each reporter channel and imaging day (days 2, 8 and 15), using material-specific mean fluorescence intensities.

**Extended Description | Material-dependent stress-reporter responses in *E. coli* (Supplementary Figs. S5–S7)**

To investigate whether engineered extracellular material environments influence bacterial physiology beyond growth-associated fluorescence, we employed the previously established RGB-S reporter system in *Escherichia coli* K-12 MG1655.[10] The plasmid combines three transcriptional modules: the SOS-responsive *sulA* promoter driving GFPmut3B, the RpoS-dependent *osmY* promoter driving mRFP1, and the RpoH-dependent *grpE* promoter driving mTagBFP2 (Supplementary Fig. S5a,b). These modules monitor promoter activities associated with DNA damage and replication stress, general stress adaptation, and proteotoxic stress, respectively.[55–57] Because these regulatory pathways can respond to overlapping physiological challenges, fluorescence reflects stress-associated transcriptional activity rather than direct evidence of specific molecular damage.

Longitudinal whole-MESIF imaging revealed pronounced material- and pathway-dependent differences in reporter fluorescence over 15 days (Fig. 3a–c; Supplementary Fig. S6a–f). DNA hydrogel-functionalized MESIF exhibited particularly elevated SOS-associated fluorescence, sustained RpoS-associated signals and an early increase in RpoH-associated fluorescence relative to several other material environments (Fig. 3a–c; Supplementary Fig. S6d–f). By contrast, catalase-functionalized MESIF generally showed comparatively low absolute reporter fluorescence, whereas the remaining environments exhibited distinct pathway-specific trajectories. These patterns indicate that material-associated physiological responses cannot be described by a uniform stress-reporter profile.

Absolute fluorescence, however, also depends on the number and fluorescence-associated state of cells within MESIF. To distinguish stress-associated reporter responses from material-dependent differences in growth-associated fluorescence, RGB-S fluorescence intensities were normalized to mean fluorescence intensities obtained from independent single-reporter growth experiments, matched by reporter channel, material condition and cultivation day. The resulting fold changes were subsequently averaged across the cultivation period (Fig. 3d–f). The DNA hydrogel remained associated with elevated relative SOS-, RpoS- and RpoH-related signals after normalization, supporting a distinctive physiological reporter state that is not explained solely by differences in absolute fluorescence. Other materials exhibited more selective relative responses, including PEG-associated RpoS and catalase-associated RpoH signals (Fig. 3e,f). Although this normalization strengthens the physiological interpretation, it does not establish the molecular cause of reporter activation or eliminate all possible differences between reporter strains.

Spatially resolved CLSM revealed heterogeneous stress-reporter fluorescence within individual MESIF pores on days 2, 8 and 15 (Fig. 3g; Supplementary Fig. S7a–c). Particularly prominent early fluorescence was observed in DNA hydrogel-functionalized pores, while the relative contributions of the three reporter channels changed over time (Fig. 3g). Comparison with fluorescence quantified from corresponding regions of interest in whole-MESIF images showed broadly concordant material-associated patterns, although the degree of agreement depended on reporter channel and cultivation day (Spearman correlations calculated from material-specific mean fluorescence intensities; Supplementary Fig. S7d–g). Thus, the spatial measurements complement the longitudinal whole-MESIF analysis but should not be treated as quantitatively interchangeable.

Together, these results identify DNA hydrogel-functionalized MESIF as an extracellular environment associated with a pronounced, multi-pathway stress-reporter phenotype in *E. coli*. This phenotype provides a physiological context for the reduced *E. coli*-associated fluorescence subsequently observed in the DNA hydrogel environment during monoculture and consortium cultivation (Fig. 4c,f; Supplementary Fig. S9c,f,i,l). The relationship remains associative: neither the reporter measurements nor the spatial imaging establish a specific DNA-dependent stress mechanism. Instead, they motivate the subsequent investigation of how the same DNA-rich material environment is associated with divergent responses among different microbial species.

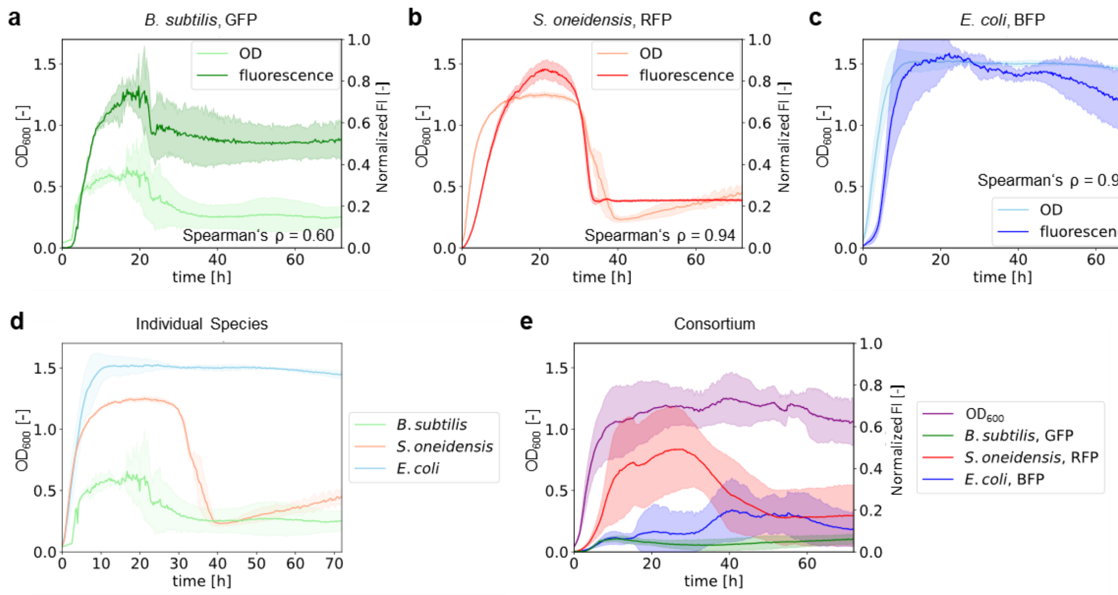


**Fig. S8 | Validation of species-specific fluorescence reporters and temporal dynamics of a three-species consortium in liquid culture.** GFP-expressing *Bacillus subtilis* 1012, RFP-expressing *Shewanella oneidensis* MR-1 and BFP-expressing *Escherichia coli* K-12 MG1655 were cultivated individually or as an initially equal-ratio three-species consortium in liquid culture at 30 °C. **(a–c)** Time-resolved optical density at 600 nm ($OD_{600}$; light-colored curves, left y-axis) and normalized fluorescence intensity (FI; dark-colored curves, right y-axis) for monocultures of *B. subtilis* (a), *S. oneidensis* (b) and *E. coli* (c). Spearman correlation coefficients between $OD_{600}$ and fluorescence intensity were $\rho$ = 0.60, 0.94 and 0.92, respectively. **(d)** Time-resolved $OD_{600}$ of the three species cultivated separately as monocultures. **(e)** Time-resolved total $OD_{600}$ of the three-species consortium (purple curve, left y-axis) together with species-specific normalized fluorescence intensities for *B. subtilis* (green), *S. oneidensis* (red) and *E. coli* (blue; right y-axis). Fluorescence intensities were normalized separately for each species by dividing each measurement by the maximum fluorescence intensity recorded for that species during monoculture cultivation. This normalization enables comparison of temporal fluorescence responses within each reporter system but does not provide a direct quantitative measure of relative species abundance or biomass in the consortium. Lines represent means and shaded areas indicate standard deviations from three independent biological cultures (n = 3).

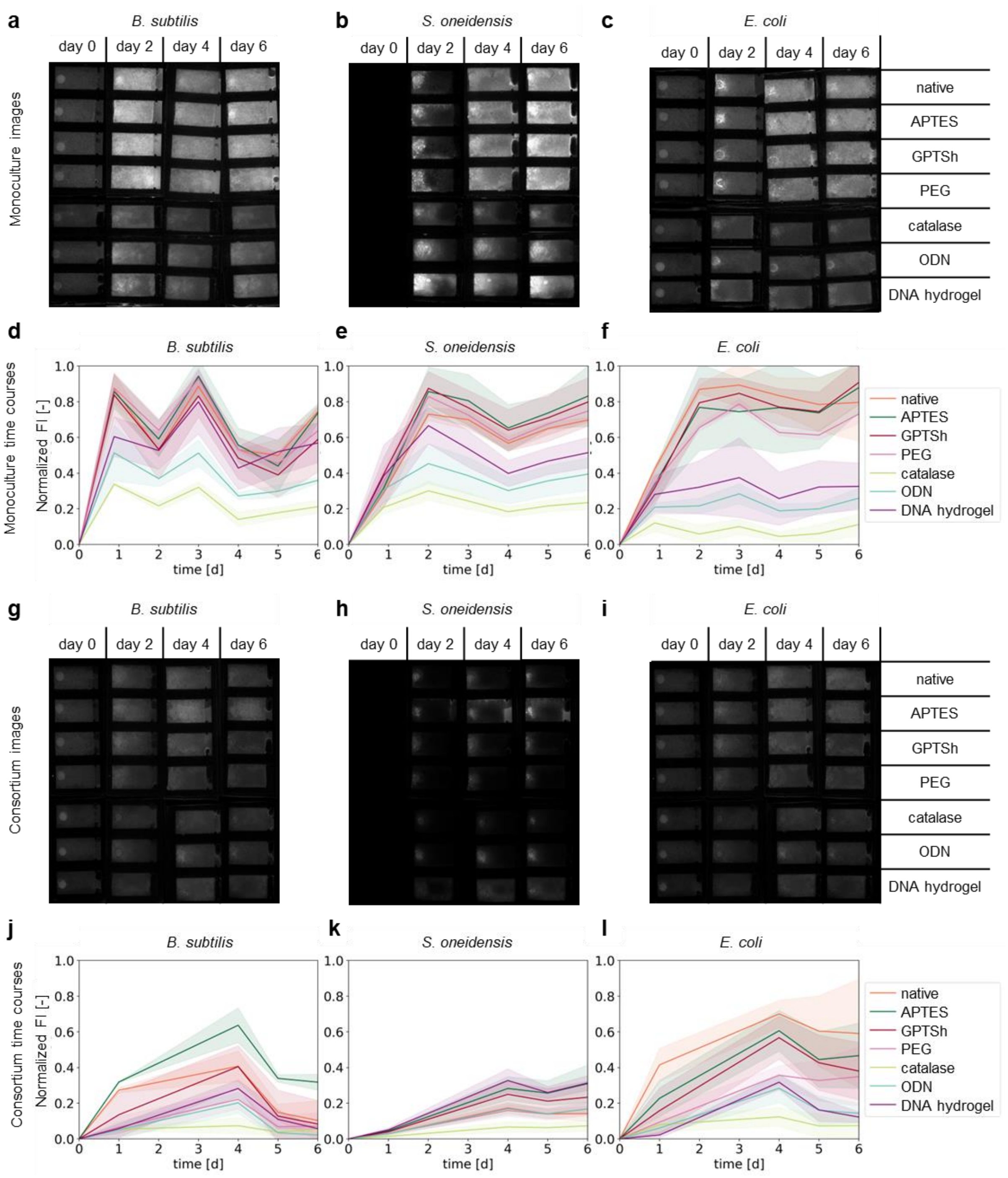


**Fig. S9 | Material- and culture-context-dependent fluorescence dynamics of individual species and a three-species consortium in MESIF.** GFP-expressing *Bacillus subtilis* 1012, RFP-expressing *Shewanella oneidensis* MR-1 and BFP-expressing *Escherichia coli* K-12 MG1655 were cultivated individually or as an initially equal-ratio three-species consortium in MESIF presenting native, APTES-, GPTSh-, PEG-, catalase-, ODN- or DNA hydrogel-functionalized material environments. **(a–c)** Representative whole-MESIF fluorescence images acquired on days 0, 2, 4 and 6 for monocultures of *B. subtilis* (a), *S. oneidensis* (b) and *E. coli* (c). Rows correspond to the seven material environments in the order indicated. **(d–f)** Corresponding longitudinal whole-MESIF fluorescence profiles of *B. subtilis* (d), *S. oneidensis* (e) and *E. coli* (f) from day 0 to day 6 during monoculture cultivation. **(g–i)** Representative whole-MESIF fluorescence images acquired on days 0, 2, 4 and 6 for *B. subtilis* (g), *S. oneidensis* (h) and *E. coli* (i) during consortium cultivation. **(j–l)** Corresponding longitudinal whole-MESIF fluorescence profiles of *B. subtilis* (j), *S. oneidensis* (k) and *E. coli* (l) from day 0 to day 6 during consortium cultivation. Fluorescence intensities were normalized separately for each species by dividing each measurement by the maximum fluorescence intensity observed for that species during monoculture cultivation. The normalized fluorescence values enable comparison of material-associated temporal responses within a species but do not directly quantify relative species abundance or biomass. Lines represent means and shaded areas indicate standard deviations from three independent MESIF chips per material environment and culture condition (n = 3), repeatedly measured over time. Statistical analysis of the longitudinal fluorescence data using linear mixed-effects models is described in Methods; no significance comparisons are displayed in this figure.

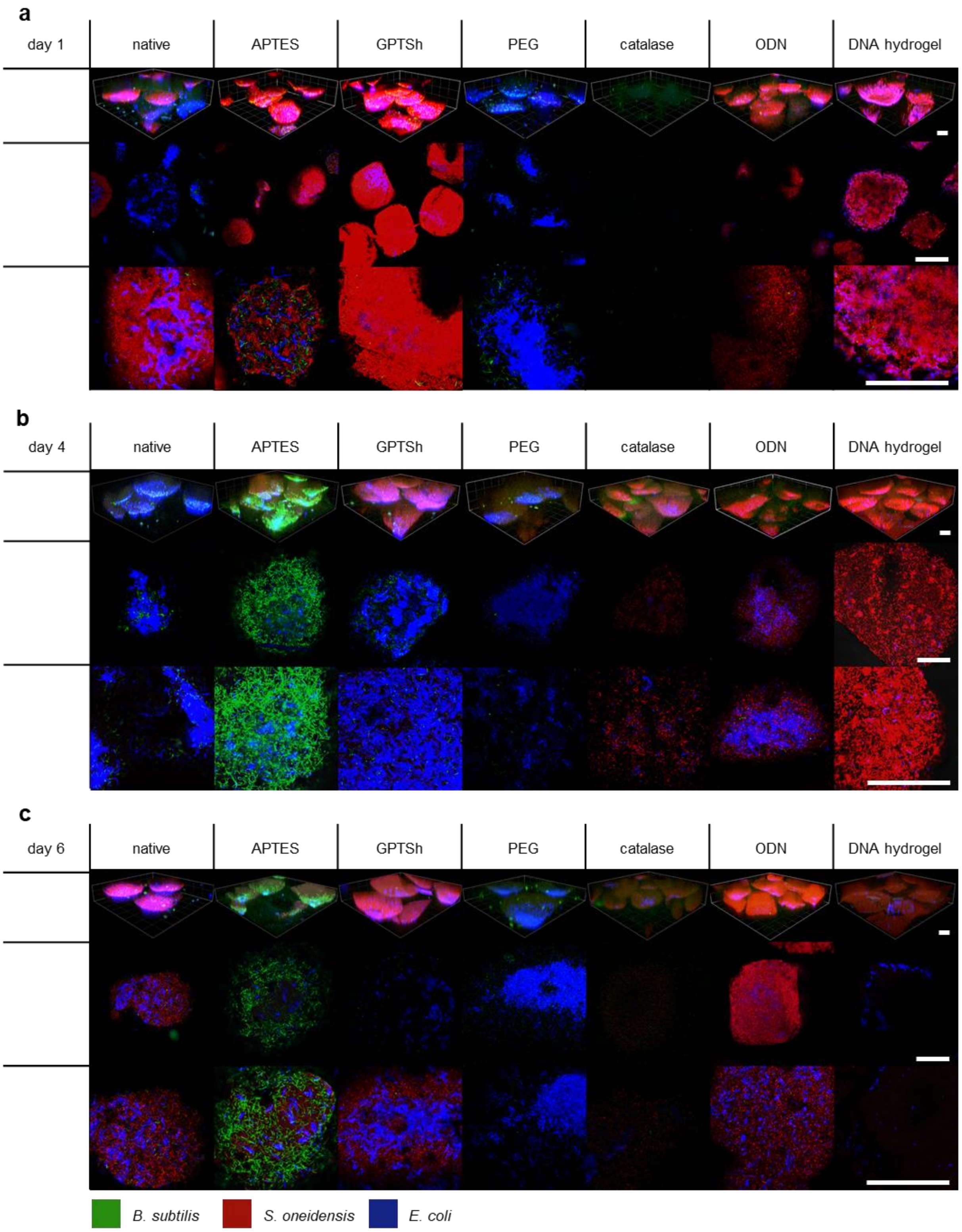


**Fig. S10 | Material- and time-dependent spatial organization of a three-species microbial consortium within MESIF pores.** GFP-expressing *Bacillus subtilis* 1012 (green), RFP-expressing *Shewanella oneidensis* MR-1 (red) and BFP-expressing *Escherichia coli* K-12 MG1655 (blue) were cultivated as an initially equal-ratio three-species consortium in MESIF presenting native, APTES-, GPTSh-, PEG-, catalase-, ODN- or DNA hydrogel-functionalized material environments. **(a–c)** Representative confocal laser scanning microscopy (CLSM) images of the consortium within MESIF pores on day 1 (a), day 4 (b) and day 6 (c). For each material environment and time point, the upper image row shows a three-dimensional representation of multiple MESIF pores, and the two lower rows show magnified views of the local species-specific fluorescence distribution within the pores. Images display merged fluorescence channels for the three consortium members. Differences in fluorescence patterns illustrate material- and time-associated variation in the spatial organization of the consortium but do not directly quantify species abundance or spatial segregation. All scale bars represent 100 µm; their displayed lengths differ between image rows owing to differences in magnification.

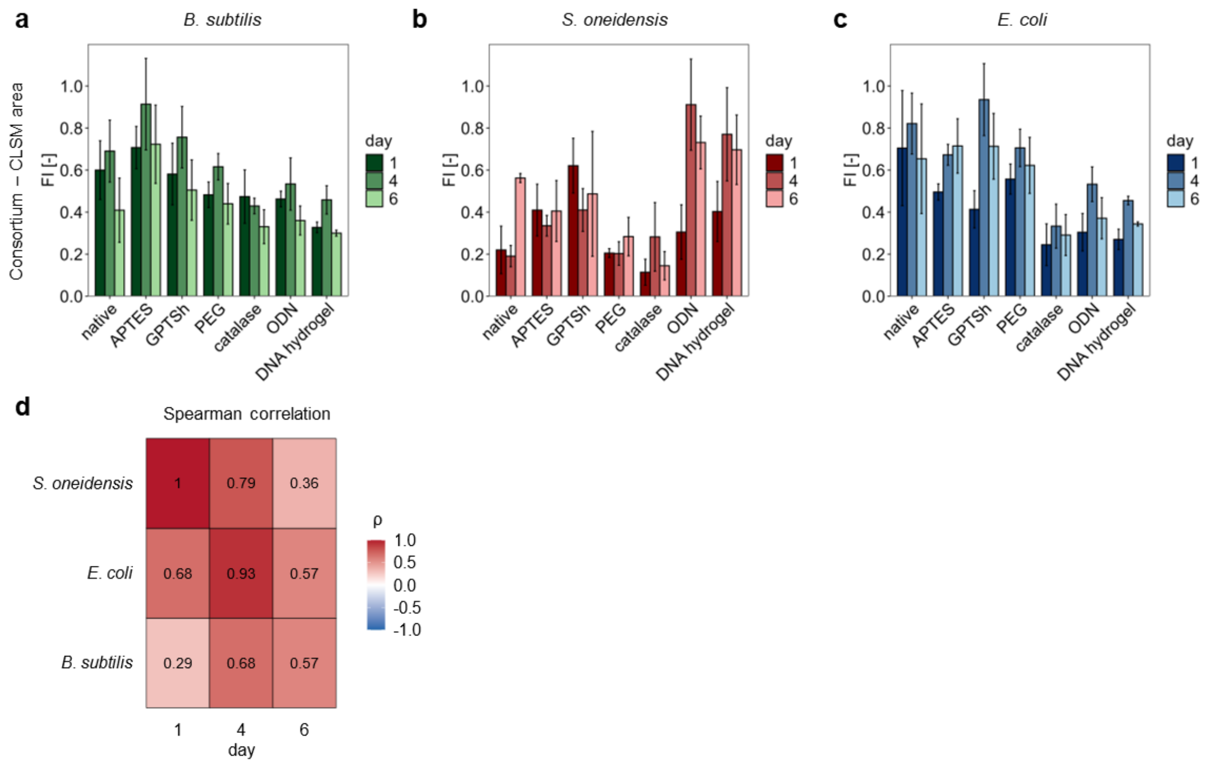


**Fig. S11 | Species- and time-dependent correspondence between whole-MESIF and spatially resolved fluorescence measurements in a three-species consortium.** GFP-expressing *Bacillus subtilis* 1012, RFP-expressing *Shewanella oneidensis* MR-1 and BFP-expressing *Escherichia coli* K-12 MG1655 were cultivated as an initially equal-ratio three-species consortium in MESIF presenting native, APTES-, GPTSh-, PEG-, catalase-, ODN- or DNA hydrogel-functionalized material environments. **(a–c)** Species-specific fluorescence intensities quantified from regions of interest (ROIs) in whole-MESIF fluorescence images corresponding to pore regions investigated by confocal laser scanning microscopy (CLSM), shown for *B. subtilis* (a), *S. oneidensis* (b) and *E. coli* (c) on days 1, 4 and 6. Fluorescence measurements were normalized separately for each species to the maximum fluorescence intensity observed for that species under monoculture conditions. Bars represent mean ± standard deviation from three independent MESIF chips per material environment (n = 3). **(d)** Spearman correlation coefficients (ρ) comparing ROI-based whole-MESIF fluorescence measurements with CLSM-derived fluorescence intensities (Fig. 4g–i), calculated separately for each species and cultivation day using mean fluorescence intensities across the seven material environments. Correlation coefficients are displayed within the heatmap cells. The degree of correspondence between the two fluorescence measurement approaches varies among species and time points; the correlations do not establish quantitative interchangeability between the methods.

**Extended Description | Species-specific temporal and spatial responses within the three-species consortium (Supplementary Figs. S8–S11)**

To establish a species-resolved readout of microbial responses in a defined multispecies community, we first characterized the relationship between fluorescence reporter signals and culture growth in liquid culture. GFP-expressing *Bacillus subtilis*, RFP-expressing *Shewanella oneidensis* and BFP-expressing *Escherichia coli* were cultivated individually, and their fluorescence trajectories were compared with optical density at 600 nm ($OD_{600}$). Fluorescence correlated positively with $OD_{600}$ for all three species, although the strength of this association differed substantially (*B. subtilis*, Spearman's $\rho = 0.60$; *S. oneidensis*, $\rho = 0.94$; *E. coli*, $\rho = 0.92$; Supplementary Fig. S8a–c). In particular, the comparatively moderate correlation for *B. subtilis* reflected differences between the temporal evolution of its fluorescence and optical-density signals. These observations support the use of fluorescence as a species-specific, growth-associated readout while emphasizing that reporter intensity is not a direct quantitative measure of biomass, viable cell number or relative species abundance.

The individual monocultures also displayed distinct growth trajectories in liquid culture (Supplementary Fig. S8d). When combined in an initially equal-ratio consortium, the organisms exhibited different fluorescence dynamics despite sharing the same liquid environment (Supplementary Fig. S8e). *S. oneidensis*-associated fluorescence increased rapidly, reached an early maximum and subsequently declined, whereas *E. coli*-associated fluorescence increased more prominently at later time points. The *B. subtilis*-associated signal remained comparatively low. These patterns demonstrate that species-specific reporter dynamics can diverge even in the absence of the engineered MESIF material environments. However, because the fluorescence signals were normalized independently for each species, differences in their amplitudes cannot be interpreted as differences in population size or as direct evidence of competitive displacement.

We next examined how these species-specific responses developed within the seven MESIF material environments. The three organisms were cultivated either individually or as an initially equal-ratio consortium in native, APTES-, GPTSh-, PEG-, catalase-, ODN- and DNA hydrogel-functionalized MESIF. Whole-MESIF fluorescence was monitored longitudinally from day 0 to day 6, with representative fluorescence images acquired on days 0, 2, 4 and 6 (Supplementary Fig. S9). Each fluorescence measurement was normalized by the maximum fluorescence intensity observed for the corresponding species during monoculture cultivation. This species-specific normalization enabled comparisons of material-associated fluorescence profiles between monoculture and consortium cultivation within the same reporter system, but did not permit direct quantitative comparisons of species abundance. The complete temporal profiles complement the time-averaged fluorescence measurements presented in Fig. 4a–f. Longitudinal measurements from independent MESIF chips were analyzed using the mixed-effects framework described in Methods, accounting for repeated measurements of the same chips over time.

In monoculture, all three organisms exhibited pronounced material-associated variation in fluorescence, although the temporal patterns differed between species (Fig. 4a–c and Supplementary Fig. S9a–f). *B. subtilis* displayed non-monotonic fluorescence trajectories, with prominent early peaks and subsequent changes in signal intensity under several material conditions (Supplementary Fig. S9a,d). *S. oneidensis* exhibited a rapid initial increase in fluorescence across several environments, followed by a transient decline and, in some cases, renewed signal accumulation towards day 6 (Supplementary Fig. S9b,e). By contrast, *E. coli* showed comparatively sustained fluorescence in native and several chemically functionalized MESIF environments, whereas catalase- and DNA hydrogel-functionalized materials were associated with lower signals (Supplementary Fig. S9c,f). These patterns demonstrate that the material-dependent fluorescence phenotypes summarized in Fig. 4a–c arise from distinct temporal trajectories rather than from uniform changes in signal intensity throughout cultivation.

The *E. coli* response was particularly relevant to the preceding single-species experiments. In the present monoculture series, DNA hydrogel-functionalized MESIF again exhibited comparatively low *E. coli*-associated fluorescence, qualitatively consistent with the material-dependent response identified using the independent *E. coli* reporter system described earlier in the manuscript (Fig. 4c and Supplementary Fig. S9c,f). This agreement across experimental series supports the reproducibility of the observed fluorescence phenotype, although the two reporter systems and experimental designs should not be interpreted as quantitatively interchangeable. During consortium cultivation, *E. coli* fluorescence remained relatively high in native MESIF and several chemically functionalized environments, while catalase, ODN and DNA hydrogel conditions generally exhibited lower signals (Fig. 4f and Supplementary Fig. S9i,l). Thus, the presence of other consortium members did not eliminate the broad material-associated pattern of the *E. coli* reporter response, although the fluorescence trajectories and their magnitudes differed from those observed in monoculture.

The response of *B. subtilis* was notably different. During monoculture cultivation, its fluorescence varied substantially over time, with several material environments supporting pronounced early signals (Supplementary Fig. S9a,d). Within the consortium, APTES-functionalized MESIF was associated with a particularly prominent increase in *B. subtilis*-derived fluorescence, reaching a maximum around day 4 before declining (Fig. 4d and Supplementary Fig. S9g,j). Other material environments generally exhibited lower consortium-associated signals. This behaviour indicates that the fluorescence phenotype of *B. subtilis* is sensitive to both material environment and culture context. However, the comparison between monoculture and consortium profiles is descriptive unless supported by a formal statistical test of the material-by-culture-context interaction. Furthermore, the fluorescence data do not establish whether the underlying changes reflect cell accumulation, reporter regulation or shifts in physiological state.

*Shewanella oneidensis* showed a third response pattern. During monoculture cultivation, relatively high fluorescence was observed in several environments, whereas catalase-functionalized MESIF generally produced lower signals (Fig. 4b and Supplementary Fig. S9b,e). In the consortium, *S. oneidensis*-associated fluorescence accumulated more gradually and reached comparatively prominent levels in several conditions, including DNA hydrogel-functionalized MESIF (Fig. 4e and Supplementary Fig. S9h,k). This response contrasted with the relatively low *E. coli*-associated fluorescence in the same DNA hydrogel environment. The comparison is particularly informative because both organisms experienced the same engineered extracellular material, yet their species-specific fluorescence profiles differed. The data therefore support a species-dependent response to the material environment, without establishing whether the divergence results from direct interspecies interactions, different intrinsic material sensitivities or other physiological factors.

This contrast is consistent with the broader observation that DNA-containing hybrid materials can elicit distinct colonization phenotypes in different bacterial species. In our previous work on rolling-circle-amplification-derived DNA hybrid matrices, *S. oneidensis* and *E. coli* exhibited different colonization responses to composites containing silica nanoparticles and carbon nanotubes.[58] In particular, favorable *S. oneidensis* colonization of a conductive SiNP/CNT–DNA composite was associated with its extracellular electron-transfer capabilities and the properties of the hybrid matrix. The present findings extend the observation of species-dependent responses to a different DNA-based material environment. However, the DNA hydrogel investigated here is not equivalent to the conductive nanocomposite used previously, and neither its electrical properties nor electron-transfer activity can be inferred from the fluorescence measurements. The earlier study therefore provides biological context rather than a demonstrated mechanism for the current DNA hydrogel-associated phenotype.

To determine how the fluorescence responses were distributed within the porous MESIF architecture, we complemented whole-MESIF measurements with spatially resolved confocal laser scanning microscopy (CLSM). Representative three-dimensional views and magnified images acquired on days 1, 4 and 6 revealed heterogeneous, material-associated fluorescence distributions within individual pores (Fig. 4j and Supplementary Fig. S10a–c). In APTES-functionalized MESIF, prominent *B. subtilis*-associated fluorescence structures were visible at later time points, particularly on day 4. PEG-functionalized MESIF frequently exhibited extensive *E. coli*-associated fluorescence, whereas GPTSh- and ODN-functionalized environments showed temporally changing combinations of species-associated signals. Catalase-functionalized MESIF displayed comparatively weak fluorescence in several representative views. These observations illustrate that differences between material environments involve not only whole-chip fluorescence trajectories but also the spatial arrangement of detectable reporter signals within pores.

The DNA hydrogel condition was especially dynamic in the CLSM images. Prominent *S. oneidensis*-associated fluorescence was visible within pores on days 1 and 4, whereas the representative day-6 views showed substantially less detectable fluorescence (Supplementary Fig. S10). This temporal change should be distinguished from the corresponding whole-MESIF fluorescence trajectories, which sample a larger material volume and may show different temporal behavior. Differences between local CLSM images and whole-MESIF measurements may reflect spatial heterogeneity, differences in sampling regions or properties of the imaging methods. The reduced fluorescence in representative late-stage images does not, by itself, establish loss of viable cells, community collapse or disappearance of *S. oneidensis* from the material.

Quantitative CLSM-derived fluorescence measurements provided a complementary view of the species-specific signals within individual pores on days 1, 4 and 6 (Fig. 4g–i). To assess how these local measurements related to larger-scale fluorescence imaging, we additionally quantified fluorescence in regions of interest (ROIs) from whole-MESIF images corresponding to the pore regions investigated by CLSM (Supplementary Fig. S11a–c). The ROI-based profiles retained material-associated differences, including relatively strong *B. subtilis* signals under APTES conditions, prominent *S. oneidensis* signals under selected ODN and DNA hydrogel conditions, and relatively high *E. coli* signals in native and several chemically functionalized environments. Nevertheless, the two measurement approaches differed in spatial sampling and should not be assumed to yield identical material rankings.

Spearman correlation analysis confirmed that the correspondence between ROI-based whole-MESIF fluorescence and CLSM-derived fluorescence depended on both species and cultivation time (Supplementary Fig. S11d). For *S. oneidensis*, the reported coefficients were $\rho = 1.00$, 0.79 and 0.36 on days 1, 4 and 6, respectively. The corresponding values were $\rho = 0.68$, 0.93 and 0.57 for *E. coli*, and $\rho = 0.29$, 0.68 and 0.57 for *B. subtilis*. Thus, strong agreement was observed for selected species–time combinations, whereas others showed only weak or moderate correspondence. These correlations, calculated from material-specific mean fluorescence intensities separately for each species and cultivation day, characterize the consistency of material-associated fluorescence patterns between measurement approaches. They do not establish quantitative interchangeability, nor do they independently demonstrate a particular spatial mechanism.

Taken together, the longitudinal and spatially resolved fluorescence measurements demonstrate that the three consortium members exhibit distinct material-associated phenotypes across both time and pore space. Importantly, the same extracellular material environment was not associated with an equivalent fluorescence response in all three species. The divergence between *S. oneidensis* and *E. coli* under DNA hydrogel conditions, the prominent *B. subtilis* response in APTES-functionalized MESIF and the heterogeneous CLSM patterns collectively illustrate how material environments can be associated with different trajectories of microbial organization in a defined consortium. These observations provide the physiological and spatial context for the species-resolved metatranscriptomic analysis presented in Extended Data Fig. 1 and Supplementary Fig. S12, which examines whether the observed fluorescence phenotypes are accompanied by corresponding differences in transcriptional state.

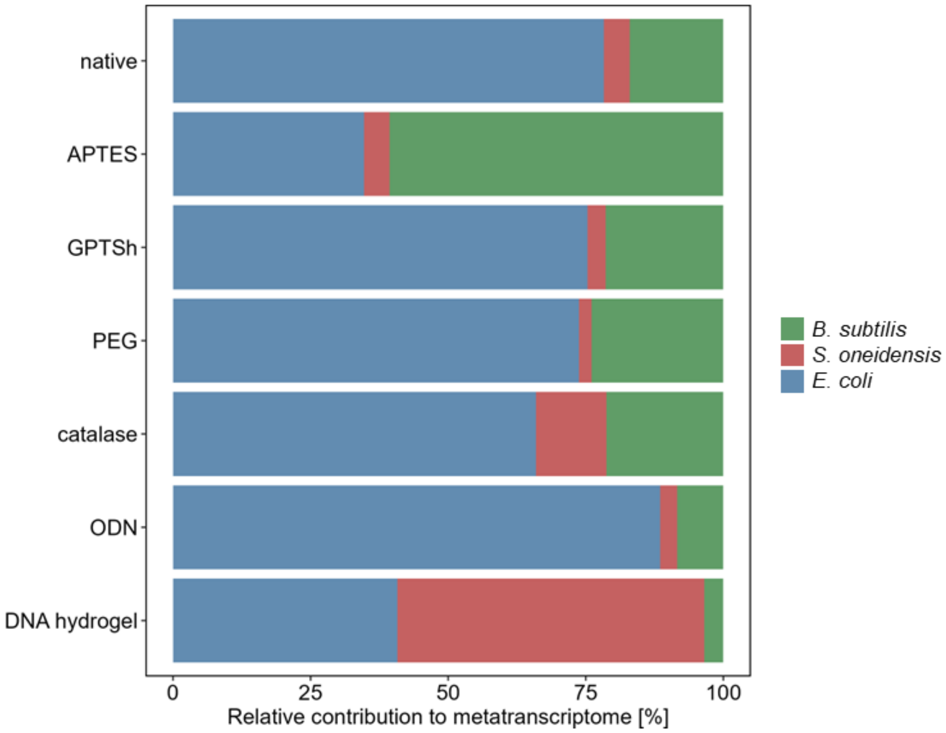


**Fig. S12 | Relative species-specific contributions to the metatranscriptome across material environments.** Metatranscriptomic analysis of a three-species consortium comprising *Bacillus subtilis*, *Shewanella oneidensis* and *Escherichia coli* cultivated for 7 days in native, APTES-, GPTSh-, PEG-, catalase-, ODN- and DNA hydrogel-functionalized MESIF environments. For each biological sample, transcript abundances were normalized as transcripts per million (TPM) across the complete metatranscriptome, with genes from all three species included in a common normalization. TPM values were summed across all genes assigned to each species and expressed as a percentage of the total TPM signal in that sample. Stacked bars show the mean relative contribution of each consortium member across biological replicates for each material environment. These values represent relative contributions to the recovered metatranscriptomic signal, not species abundances or absolute transcriptional activities.

**Extended Description | Material-dependent phenotypes are accompanied by species-specific transcriptional states**

To investigate the molecular responses accompanying material-dependent growth, stress physiology and community organization, we performed metatranscriptomic profiling of MESIF-associated microbial communities after seven days of cultivation. The fluorescent three-species consortium comprising *B. subtilis*, *S. oneidensis* and *E. coli* was cultivated in native, APTES-, GPTSh-, PEG-, catalase-, ODN- and DNA hydrogel-functionalized MESIF. Transcriptomes from the three species grown individually in native MESIF served as reference conditions for examining the influence of community context. Together with the physiological and spatial measurements presented in Figs. 3 and 4, these datasets enabled us to examine whether a shared extracellular material environment was associated with distinct molecular responses in individual consortium members.

Because RNA was extracted from entire MESIF chips, the metatranscriptomic data represent bulk transcriptional profiles of the recovered communities. They do not resolve spatial heterogeneity within individual pores or distinguish transcriptional contributions from different spatial subpopulations. Sequencing reads were assigned to the respective microbial species, and expression profiles were analyzed separately for each organism to avoid conflating interspecies differences with material-associated transcriptional variation. Raw gene counts were processed using the variance-stabilizing transformation implemented in DESeq2.[20] Principal component analysis (PCA) was then performed independently for each species using the 500 most variable genes across the analyzed consortium and native monoculture samples (Extended Data Fig. 1a–c). These species-resolved analyses were complemented by an assessment of the relative contributions of all three organisms to the recovered metatranscriptome, using TPM normalization across genes from all three species within each sample (Supplementary Fig. S12).

The resulting ordinations revealed distinct species-dependent patterns of transcriptional variation. For *B. subtilis*, the first principal component explained 77.8% of the variance, with DNA hydrogel-associated consortium samples occupying a markedly different region of the ordination from several other material environments (Extended Data Fig. 1a). *S. oneidensis* displayed a more distributed pattern across the first two components, which explained 28.7% and 27.4% of the variance, respectively (Extended Data Fig. 1b). In *E. coli*, the first component accounted for 58.1% of the variance, and material-associated differences were evident alongside variation between consortium and native monoculture samples (Extended Data Fig. 1c). These contrasting ordination patterns indicate that the three organisms did not exhibit a uniform global transcriptional response to the material environments. However, the PCA itself does not establish the statistical significance of individual material contrasts.

A complementary view of the consortium transcriptional profiles is provided by the relative species-specific contributions to the recovered metatranscriptome (Supplementary Fig. S12). Transcript abundances were normalized jointly across all genes from the three consortium members within each sample, and species-specific TPM values were summed to determine their relative contributions to the total metatranscriptomic signal. These contributions differed markedly among material environments. Most notably, DNA hydrogel-functionalized MESIF exhibited a substantially increased relative contribution from *S. oneidensis*, accompanied by a lower relative contribution from *E. coli* compared with several other environments. This pronounced redistribution of transcript contributions complements the divergent species-specific fluorescence phenotypes observed in Fig. 4 and identifies the DNA hydrogel as an environment associated with a distinctive consortium-level transcriptional profile. Importantly, these relative contributions do not directly quantify species abundance, absolute transcript production or per-cell transcriptional activity.

To examine the functional breadth of material-associated transcriptional responses, differential gene expression was analysed separately for each species using DESeq2 likelihood-ratio tests.[20] For each species, material-associated differential expression was assessed exclusively within consortium samples by comparing a full model containing material environment with an intercept-only model. Significantly differentially expressed genes were defined by a Benjamini–

Hochberg-adjusted p value below 0.05, without an additional fold-change cutoff. Low-abundance genes were filtered before analysis, and genes showing significant expression variation associated with material environment were assigned to Clusters of Orthologous Groups (COG) functional categories. The proportion of significant genes within each category was calculated relative to the total number of annotated genes assigned to that category for the respective species; genes assigned to multiple COG categories could contribute to more than one category (Extended Data Fig. 1d).

The resulting profiles revealed broad but heterogeneous material-associated transcriptional responses across functional categories involved in metabolism, cellular regulation, stress adaptation and environmental interactions. The distribution of significant genes differed among the three organisms, consistent with species-specific responses to the same set of material environments. Because the analysis summarizes the proportion of significant genes rather than their effect sizes or direction of regulation, these values describe the functional breadth of transcriptional variation, not the magnitude of physiological change.

The influence of community context was examined separately by comparing transcriptional profiles obtained during native monoculture and native consortium cultivation (Extended Data Fig. 1e). For each species, differential expression was assessed exclusively within native MESIF samples using DESeq2 likelihood-ratio tests comparing a full model containing cultivation condition with an intercept-only model. This analysis identified a different distribution of significant genes across functional categories from that associated with material environment. The generally smaller proportions of significant genes in this comparison are consistent with a more restricted detectable response to the change in culture context under the conditions tested. However, the numbers of significant genes in the material and culture-context analyses cannot alone establish which factor has the greater biological effect, because the statistical comparisons differ in design and potentially in power. In particular, the absence of detected significant genes for a species should not be interpreted as evidence that its transcriptional program is unaffected by the presence of other organisms.

To connect these global transcriptional patterns with physiological processes, we examined genes manually selected according to their annotated functions and biological relevance, encompassing surface-associated lifestyles, motility, developmental regulation, stress responses, respiration and metabolism (Extended Data Fig. 1f–h). These gene-level profiles provide a biological framework for interpreting material-associated transcriptional states. Importantly, the heatmaps show normalized transcript abundance rather than gene-wise statistical significance; differences in color therefore do not necessarily correspond to significant differential expression.

In *B. subtilis*, transcriptional profiles varied across genes involved in extracellular matrix formation, motility, competence, stress regulation and developmental transitions (Extended Data Fig. 1f). The matrix-associated genes *epsA*, *yqxM* and *tasA*, together with regulators including *sinI*, *sinR* and *slrR*, displayed material-dependent abundance patterns. The simultaneous representation of matrix-related and motility-associated transcripts, including *sigD* and *hag*, is consistent with the coexistence of regulatory programs associated with alternative lifestyles. Competence-associated genes, including *comA* and *comK*, and developmental regulators such as *spo0A* and sporulation-associated sigma factors were also represented. Of particular interest, the late sporulation-associated regulator *sigG* showed a comparatively prominent transcript signal in the DNA hydrogel condition. These observations provide a molecular context for the material-dependent fluorescence and growth phenotypes of *B. subtilis* observed in Fig. 4. Nevertheless, bulk transcriptional profiles cannot determine whether distinct programs occur in separate cells or establish progression to mature spores. Variation in transcripts associated with fermentative and respiratory metabolism, including *ldh* and *ackA*, further suggests that the metabolic state of *B. subtilis* is sensitive to the local material environment.

*Shewanella oneidensis* exhibited a different transcriptional repertoire, encompassing adhesion, motility, stress regulation and respiratory functions (Extended Data Fig. 1g). Transcripts associated with surface attachment included the MSHA pilus genes *mshA* and *mshB*, alongside *bpfA* and *aggA*. Their relative abundance varied among material environments, suggesting differences in the transcriptional regulation of adhesion-associated pathways. Genes involved in motility and surface-associated behavior, including *fliA*, *motA* and the c-di-GMP-associated regulators represented in the heatmap, also showed heterogeneous profiles. The DNA hydrogel environment was associated with a distinctive combination of stress-related and respiratory transcripts, including *rpoS*, *recA* and genes associated with extracellular electron transfer, such as *mtrA*, *mtrB*, *mtrC* and *omcA*. These patterns are compatible with changes in the physiological state of *S. oneidensis* under different material conditions. However, transcript abundance alone does not demonstrate electron-transfer activity or establish the presence of a specific redox gradient. Together with the divergent species-specific phenotypes observed in Fig. 4, these data identify *S. oneidensis* as a particularly informative example of how material environments may differentially influence the physiology of individual community members.

In *E. coli*, the selected transcript profiles encompassed biofilm-associated regulation, stress adaptation, DNA damage responses and respiratory metabolism (Extended Data Fig. 1h). Genes involved in curli production and surface-associated behavior, including *csgD* and *csgA*, showed variation among material environments, as did c-di-GMP-associated genes such as *ydeH* and *yhjH*. Transcripts associated with general stress (*rpoS*), proteotoxic stress (*rpoH*, *dnaK* and *ibpA*), oxidative stress (*oxyR*, *ahpC* and *sod* genes) and the SOS response (*recA*, *sulA* and *umuD*) were represented across the tested environments. These molecular profiles provide an independent transcriptional context for the stress-related phenotypes measured using the RGB-S reporter system (Fig. 3), although reporter output and bulk transcript abundance represent different levels of physiological regulation and need not correspond quantitatively. Genes associated with aerobic and lower-oxygen respiratory pathways, including *cyoA*, *cydA* and *narG*, together with metabolic genes such as *ackA*, indicate the presence of transcriptional programs compatible with metabolic flexibility. Their simultaneous detection in bulk samples does not, however, directly establish oxygen gradients or metabolite exchange within the MESIF structure.

Across the three organisms, DNA hydrogel-functionalized MESIF was associated with several distinctive transcriptional features, but these differed substantially in their functional composition. *B. subtilis* exhibited marked global transcriptomic separation and variation in developmental and matrix-associated programs; *S. oneidensis* displayed a characteristic combination of adhesion-, stress- and respiration-associated transcripts; and *E. coli* showed profiles involving surface-associated regulation and multiple stress-response pathways. This heterogeneity is consistent with the physiological and spatial differences observed independently in the defined consortium (Figs. 3 and 4). It also emphasizes that the presence of a common extracellular material does not impose an identical molecular response on all community members.

The porous MESIF architecture and the physicochemical properties of the functionalized materials may generate distinct local conditions for microbial growth. In particular, spatial differences in solute availability, oxygen supply and extracellular matrix accumulation could contribute to heterogeneous physiological states, as discussed in the context of microbial biofilms and microenvironmental gradients.[59–61] These possibilities are consistent with the coexistence of transcripts associated with different metabolic and lifestyle programs, but the present bulk RNA measurements do not directly resolve local chemical gradients or establish their causal contribution to gene expression. Similarly, the detection of competence-, DNA-processing- or stress-associated transcripts does not demonstrate uptake or genomic incorporation of extracellular DNA in DNA hydrogel materials.

Collectively, the metatranscriptomic data establish a molecular correlate of the material-dependent physiological responses observed in the defined microbial consortium. The seven material environments were associated with distinct, species-resolved transcriptional states spanning surface-associated behavior, stress regulation, development and metabolism. Notably, DNA hydrogel-functionalized MESIF was also associated with a pronounced shift in the relative contributions of individual species to the recovered metatranscriptome, particularly towards *S. oneidensis*. Rather than producing a uniform transcriptional response, engineered extracellular material environments were associated with different physiological programs in each consortium member. These findings strengthen the view that MESIF material environments can influence microbial behavior at multiple biological levels and provide a mechanistically informative, although not mechanistically conclusive, foundation for investigating material-associated community assembly in more complex environmental microbiomes.

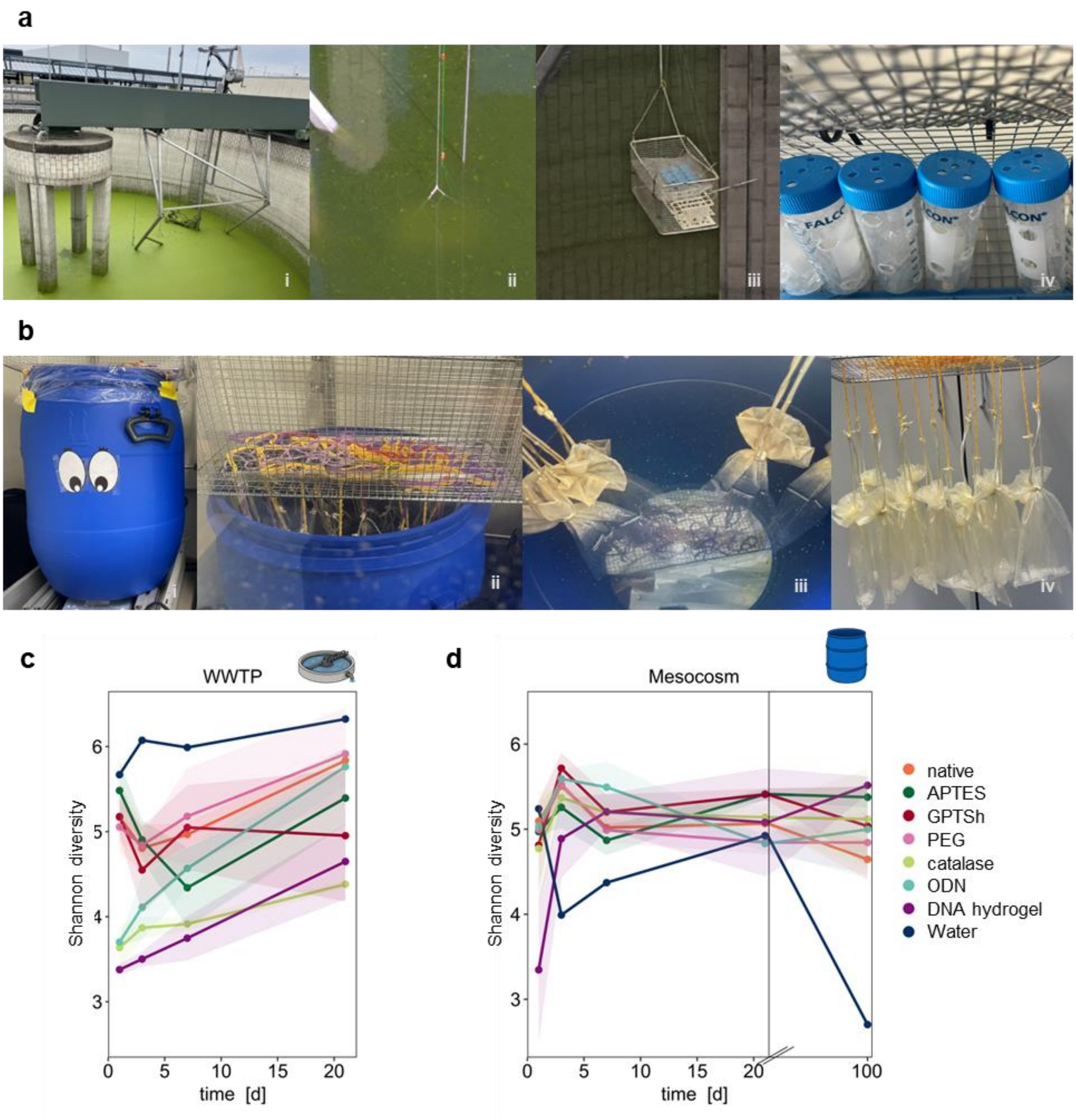


**Fig. S13 | Environmental cultivation of MESIF and temporal development of microbial alpha diversity.** MESIF presenting different material environments were exposed to complex environmental microbial communities in a chemical wastewater treatment plant (WWTP) or a wastewater-derived mesocosm. **(a)** Experimental setup for in situ cultivation in the WWTP over 1, 3, 7 and 21 days. MESIF chips were placed in perforated tubes mounted in a metal holder and submerged directly in the wastewater treatment tank, exposing the material environments to continuously changing wastewater under operational mixing conditions. Photographs (i–iv) document the cultivation tank, the deployment apparatus and the perforated tubes containing MESIF chips. **(b)** Experimental setup for long-term cultivation in a wastewater-derived mesocosm over 1, 3, 7, 21 and 100 days. MESIF chips were enclosed in mesh bags and suspended in 40 L wastewater maintained under continuous mixing at 180 rpm. Photographs (i–iv) show the mesocosm vessel and the suspended mesh-bag containing the respective MESIF chips. Before environmental exposure, MESIF chips were filled with autoclaved and sterile-filtered wastewater corresponding to the respective cultivation environment. **(c,d)** Temporal development of microbial alpha diversity in the WWTP (c) and mesocosm (d), quantified by the Shannon diversity index calculated from 16S rRNA gene amplicon sequencing data. Colors indicate native, APTES-, GPTSh-, PEG-, catalase-, ODN- and DNA hydrogel-functionalized MESIF and the corresponding water control. Lines indicate mean values and shaded regions represent ± standard deviation across three independently cultivated MESIF chips per material environment and sampling time. DNA hydrogel-associated communities showed comparatively low initial alpha diversity, followed by increasing diversity during community development. During prolonged mesocosm cultivation, Shannon diversity values among MESIF-associated communities became more similar, whereas the water control showed a pronounced decrease at day 100.

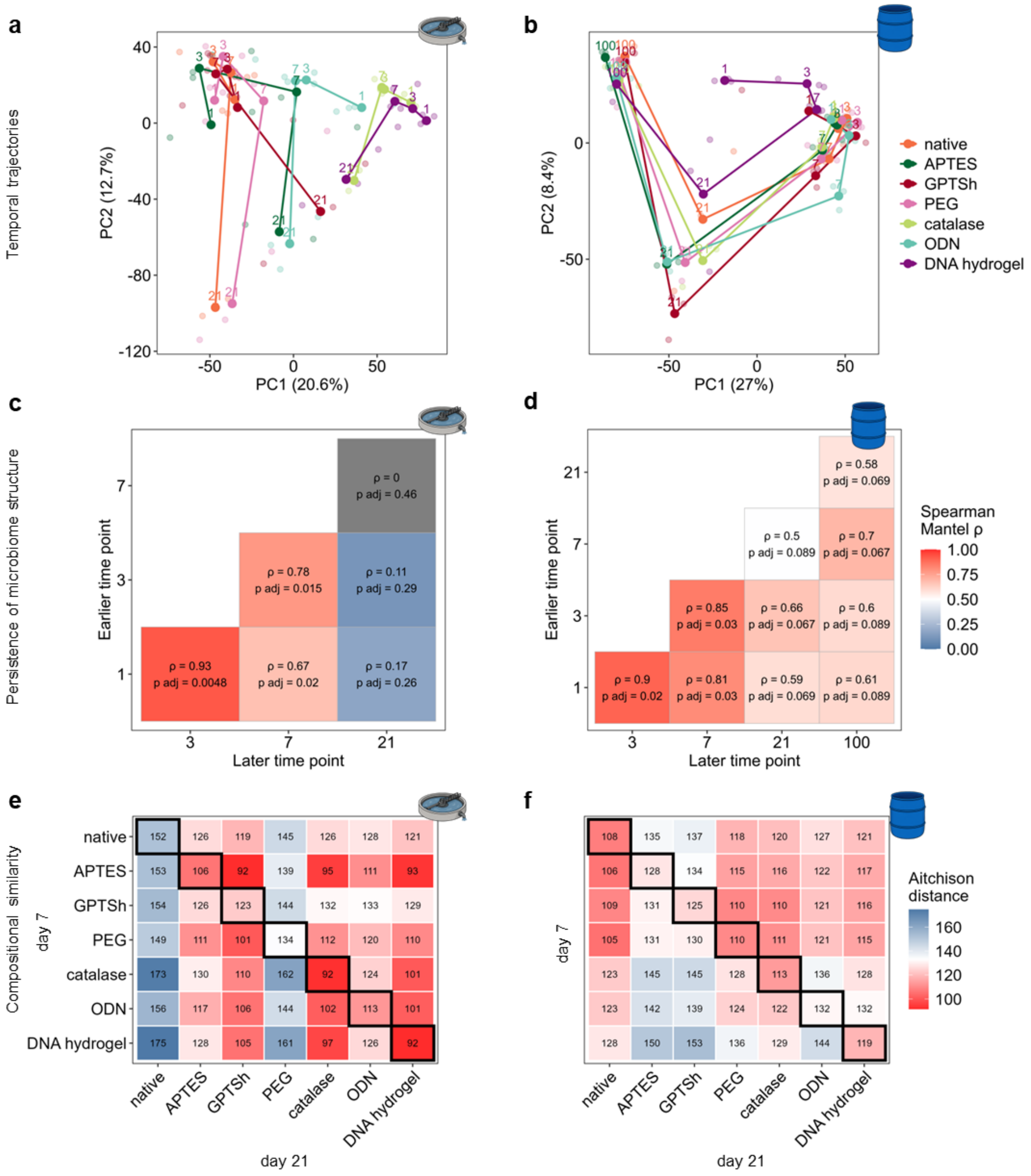


**Fig. S14 | Temporal reorganization and correspondence of material-associated microbial community structure. (a,b)** Temporal trajectories of microbial community composition associated with different MESIF material environments in the wastewater treatment plant (WWTP; a) and wastewater-derived mesocosm (b), visualized by principal component analysis (PCA) of centred log-ratio (CLR)-transformed amplicon sequence variant (ASV) abundances. Individual points represent biological replicates. For each material environment and sampling time, centroids were calculated from the mean PCA coordinates of biological replicates and connected in chronological order to visualize changes in community composition. Numbers indicate sampling days; axis labels show the proportion of variance explained by each principal component. **(c,d)** Temporal correspondence of the relative compositional relationships among material-associated communities in the WWTP (c) and mesocosm (d). Pairwise Aitchison-distance matrices between material-specific community centroids were calculated separately for each sampling time, and correspondence between time points was assessed using Spearman Mantel correlations with 9,999 permutations. Tiles show Mantel correlation coefficients (ρ) and Benjamini–Hochberg-adjusted *p* values calculated separately for WWTP and mesocosm datasets. Higher positive correlations indicate greater similarity in the relative arrangement of material-associated communities between sampling times, rather than unchanged community composition. **(e,f)** Pairwise Aitchison distances between material-specific community centroids at day 7 and day 21 in the WWTP (e) and mesocosm (f), calculated from CLR-transformed ASV abundances. Rows represent day-7 centroids and columns represent day-21 centroids. Outlined diagonal cells compare the same material environment across time, whereas off-diagonal cells compare different material environments. Lower distances indicate greater compositional similarity. These cross-time distance matrices provide descriptive comparisons and do not independently establish persistence of individual microbial populations.

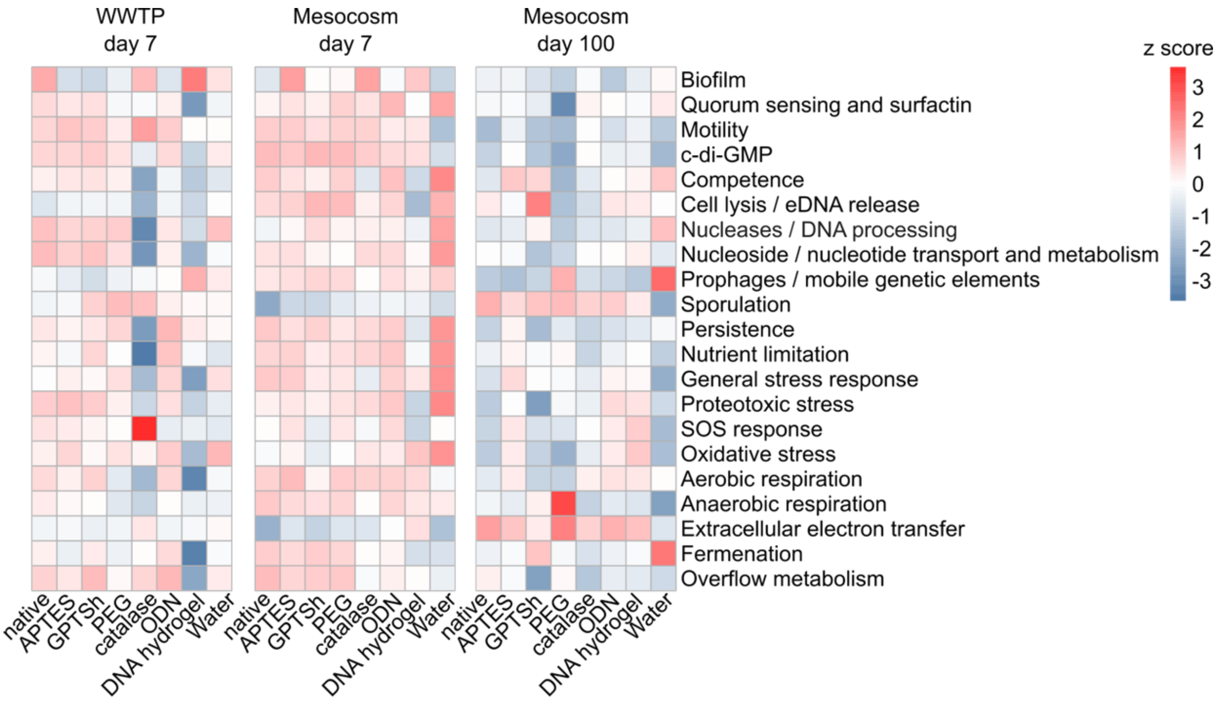


**Fig. S15 | Material environments are associated with distinct functional gene profiles in environmental microbial communities.** Functional gene profiles were derived from shotgun metagenomic sequencing of MESIF-associated microbial communities after 7 days of cultivation in the wastewater treatment plant (WWTP) and after 7 and 100 days in a wastewater-derived mesocosm. Columns represent native, APTES-, GPTSh-, PEG-, catalase-, ODN- and DNA hydrogel-functionalized MESIF and the corresponding water control. Rows represent manually curated functional categories encompassing surface-associated lifestyles and signaling, DNA processing and metabolism, mobile genetic elements, developmental and persistence-related functions, stress responses, and respiratory and metabolic processes. Functional assignments were obtained from eggNOG-mapper annotations using curated combinations of gene identifiers, annotation keywords and Pfam domains. Individual genes could contribute to multiple functional categories when their annotations matched more than one curated assignment criterion. Gene abundances were estimated from metagenomic read-mapping counts and normalized to sequencing depth as counts per million mapped reads (CPM). For visualization, $\log_2(\text{CPM} + 0.1)$ values were standardized separately for each functional category by calculating Z-scores across all displayed samples simultaneously. Red and blue indicate comparatively higher and lower standardized abundances, respectively, within each functional category; they do not indicate statistical enrichment or depletion. Each column represents a single metagenomic dataset, without averaging across biological replicates. The functional profiles are therefore descriptive and do not independently establish differences in functional activity or gene expression between material environments.

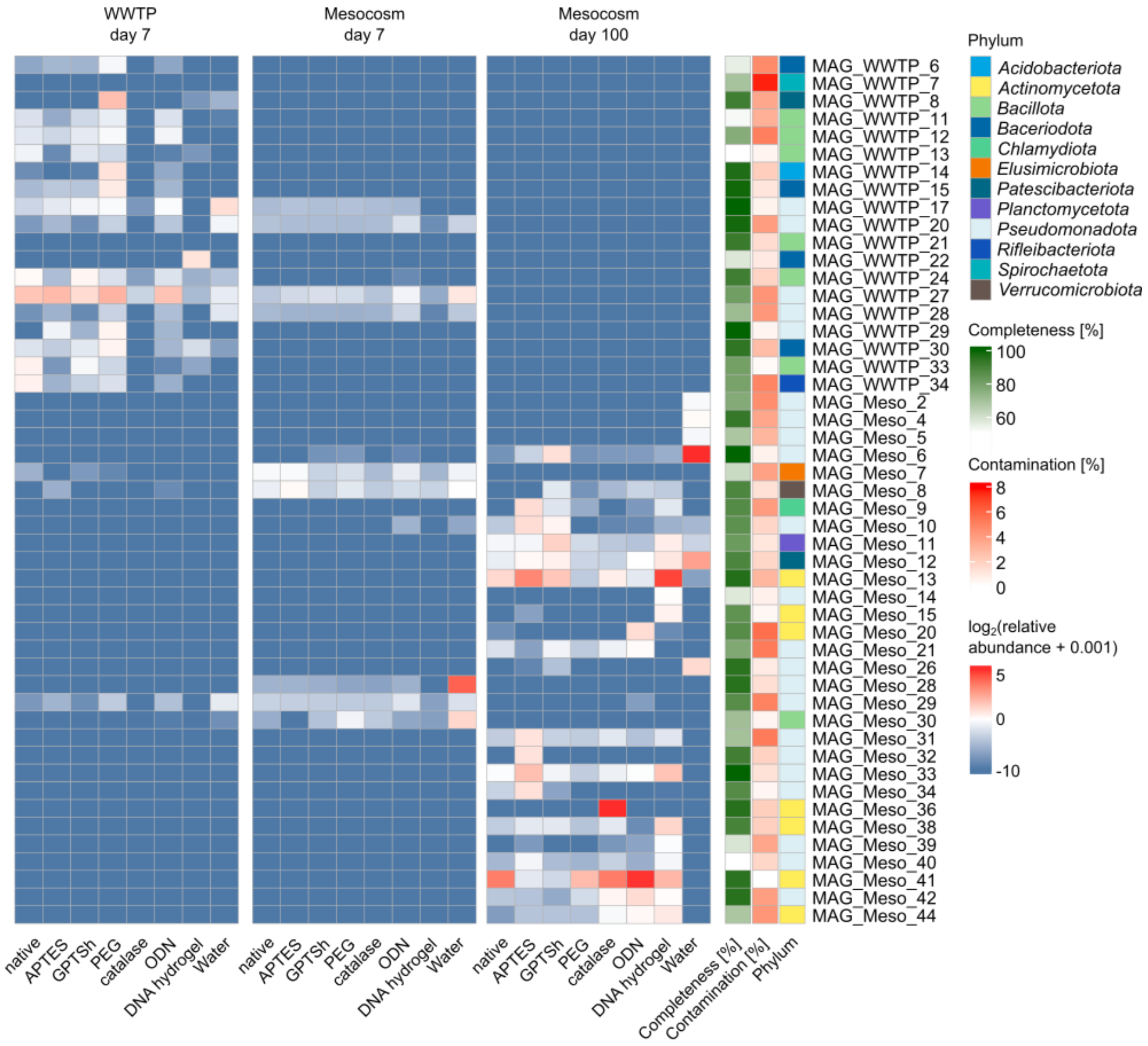


**Fig. S16 | Material environments are associated with shifts in the relative representation of individual microbial populations.** Heatmaps show relative abundance profiles of metagenome-assembled genomes (MAGs) recovered from MESIF-associated microbial communities after 7 days of cultivation in the wastewater treatment plant (WWTP) and after 7 and 100 days in a wastewater-derived mesocosm. Columns represent native, APTES-, GPTSh-, PEG-, catalase-, ODN- and DNA hydrogel-functionalized MESIF and the corresponding water control. Rows represent MAGs without species-level taxonomic assignments. Relative abundance estimates were obtained by mapping metagenomic sequencing reads to reconstructed MAGs using CoverM and visualized after $\log_2$ transformation with a pseudocount of 0.001. MAGs not detected in a given dataset had an original relative abundance of zero and are displayed at the lower end of the transformed color scale. Annotation columns indicate estimated genome completeness, contamination and phylum-level taxonomic affiliation. A total of 49 MAGs are displayed, comprising 19 MAGs recovered from the WWTP dataset and 30 from the mesocosm dataset. Each heatmap column represents one metagenomic dataset; no biological replicate averaging or statistical differential-abundance analysis was performed. The profiles therefore provide a descriptive comparison of the relative representation of reconstructed microbial populations across material environments and sampling times.

**Extended Description | Material-dependent assembly and succession of environmental microbial communities (Supplementary Figs. S13-S16)**

To determine whether material-dependent microbial responses remain detectable in taxonomically complex environmental communities, MESIF presenting distinct material environments were cultivated in two wastewater-derived settings with different experimental characteristics (Fig. S13a,b). For short-term *in situ* cultivation, MESIF were placed in perforated tubes and submerged directly in a chemical wastewater treatment tank for 1, 3, 7 and 21 days. The wastewater treatment plant (WWTP) provided a strongly agitated, operationally dynamic environment in which microbial community assembly could be examined under direct environmental exposure. For prolonged cultivation, a mesocosm was established using 40 L of chemical wastewater obtained from the same WWTP and maintained under continuous mixing at 180 rpm. MESIF were enclosed in mesh bags and cultivated for 1, 3, 7, 21 and 100 days. Previous characterization of the MESIF platform demonstrated sustained diffusion of nutrients and solutes through its interconnected porous matrix, enabling exchange between the internal material environment and the surrounding medium throughout cultivation.[1] Before deployment, MESIF were filled with autoclaved and sterile-filtered wastewater corresponding to the respective cultivation environment. Together, the two experimental settings enabled comparison of material-associated microbial community development during direct WWTP exposure and prolonged mesocosm succession.

Analysis of microbial alpha diversity revealed distinct temporal patterns among material environments (Fig. S13c,d). In the WWTP, the water control maintained comparatively high Shannon diversity, whereas communities associated with chemically modified MESIF generally exhibited relatively high diversity throughout cultivation. Several biomolecular environments showed lower diversity during early colonization, followed by increasing diversity over time. This pattern was particularly apparent for ODN-functionalized MESIF, whose Shannon diversity approached that of several chemically modified environments by day 21. DNA hydrogel-associated communities also showed comparatively low diversity during initial colonization. In the mesocosm, the low initial diversity of DNA hydrogel-associated communities was especially pronounced at day 1, followed by a rapid increase. By day 100, Shannon diversity values among MESIF-associated communities were comparatively similar, whereas the water control showed a marked decrease. The DNA hydrogel thus influenced early community assembly without preventing subsequent diversification. Importantly, the increasing similarity of Shannon diversity values did not imply convergence towards a common community composition.

Beta-diversity analysis revealed significant material-associated differentiation of microbial community composition in both cultivation settings (Fig. 5a,b). In the WWTP, communities associated with catalase-, ODN- and DNA hydrogel-functionalized MESIF occupied regions of the ordination that differed from several chemically modified environments and the water control. In the mesocosm, DNA hydrogel-associated communities also displayed pronounced compositional differentiation. Across all sampling times, material environment was significantly associated with community composition in both systems (PERMANOVA based on Bray–Curtis dissimilarities; WWTP: $R^2 = 0.39$, $p = 0.001$; mesocosm: $R^2 = 0.15$, $p = 0.001$). Sampling time was likewise associated with community composition (WWTP: $R^2 = 0.16$, $p = 0.001$; mesocosm: $R^2 = 0.43$, $p = 0.001$), consistent with pronounced temporal restructuring, particularly during prolonged mesocosm cultivation (Fig. 5c,d). Because material environment and sampling time were evaluated in separate PERMANOVA models, these $R^2$ values describe their respective associations with community variation rather than independent contributions from a joint variance-partitioning model. Nevertheless, the contrasting patterns indicate that material-associated differentiation was more prominent in the WWTP dataset, whereas sampling time showed the stronger association in the mesocosm dataset. Thus, material identity remained an important ecological variable even in a system undergoing substantial temporal succession.

To assess how material-associated differentiation developed during succession, community composition was analysed separately at each sampling time using Aitchison distances calculated from centred log-ratio-transformed amplicon sequence variant (ASV) abundances (Fig. 5e,f).

Material environment remained significantly associated with community composition at all examined time points, although the corresponding PERMANOVA $R^2$ values generally declined during cultivation. In the WWTP, the proportion of compositional variation explained by material environment decreased from approximately 0.59 at day 1 to 0.48 at day 21. In the mesocosm, it decreased from approximately 0.52 during early cultivation to approximately 0.40 at day 100. Notably, even after 100 days of mesocosm succession, material environment still accounted for approximately 40% of the observed compositional variation. Rather than erasing the influence of the extracellular material environment, prolonged succession therefore occurred alongside persistent material-associated community differentiation. The WWTP day-1 result additionally showed a significant difference in multivariate dispersion among groups (PERMDISP, Benjamini–Hochberg-adjusted $p = 0.0245$), warranting caution in attributing that particular PERMANOVA result exclusively to differences in group centroids.

Persistence of a detectable material-associated effect did not imply that the relationships among material-specific communities remained fixed over time. Principal component analysis of centred log-ratio-transformed ASV profiles revealed distinct temporal trajectories for individual material environments in both cultivation systems (Fig. S14a,b). Mantel comparisons of material-specific Aitchison-distance matrices further indicated that the relative configuration of communities was more similar between several early sampling points than between more widely separated stages of succession, particularly in the WWTP (Fig. S14c,d). Direct comparisons of material-specific community centroids between days 7 and 21 showed heterogeneous cross-time compositional distances among material environments (Fig. S14e,f). The DNA hydrogel-associated community exhibited a comparatively small cross-time distance in the WWTP, whereas the corresponding mesocosm distance was within the range observed for other material environments. These analyses illustrate continuing community reorganization and changes in the relative relationships among material-associated assemblages. They do not, by themselves, establish the persistence of individual microbial populations or unchanged community composition. The ecological imprint of a material is therefore better understood as a continuing influence on community assembly than as the preservation of a fixed community state.

The distinction between the two DNA-containing material environments was particularly informative. Although both ODN- and DNA hydrogel-functionalized MESIF presented extracellular DNA within the pore environment, their associated communities differed in alpha-diversity dynamics, compositional trajectories and their positions relative to other material-associated communities (Figs. 5, S13 and S14). DNA hydrogel-associated communities showed particularly pronounced differentiation in the mesocosm, whereas ODN-associated communities followed a distinct trajectory. These divergent ecological responses indicate that introducing DNA into an extracellular material environment does not produce a single, generic DNA-associated community phenotype. Instead, the DNA hydrogel establishes a highly concentrated, three-dimensional DNA-rich material environment that differs fundamentally from the presentation of DNA as ODNs. Importantly, ODN and DNA hydrogel differ in DNA abundance, polymer length, local density and three-dimensional organization. The present comparison therefore does not isolate the contribution of any single material property to the observed ecological differences.

Shotgun metagenomic sequencing provided a complementary view of community taxonomic composition and functional gene representation. Taxonomic profiles varied among material environments and changed substantially during prolonged mesocosm cultivation (Fig. 5g). Functional profiling further revealed heterogeneous relative representation of genes associated with surface-associated lifestyles, signaling, stress responses, respiration, metabolism and DNA-related processes (Fig. S15). The ODN- and DNA hydrogel-associated communities displayed non-equivalent functional profiles, including differences in categories related to biofilm formation, motility and c-di-GMP signaling. In contrast, categories associated with competence, nucleases and nucleotide transport or metabolism did not show a consistent increase in relative representation in DNA hydrogel-associated communities across the examined datasets. These observations provide functional genomic context for the material-associated community profiles but do not demonstrate increased extracellular DNA uptake, degradation or utilization. Because the metagenomic profiles were derived from single datasets per condition, the differences are descriptive rather than statistically validated material-specific effects. Moreover, gene abundance reflects genomic potential rather than gene expression or biochemical activity and cannot establish the molecular mechanisms underlying the hydrogel–microbial sequence associations examined in Fig. 6.

Genome-resolved analysis provided a further population-level perspective (Fig. S16 and Supplementary Table S4). Of the 86 recovered metagenome-assembled genomes (MAGs), 49 lacked species-level taxonomic assignments, comprising 19 MAGs from the WWTP and 30 from the mesocosm datasets. The displayed MAGs exhibited heterogeneous relative abundance profiles across material environments and cultivation times. Several MAGs were represented in multiple material-associated communities, while their relative abundances differed among samples. These descriptive patterns are consistent with material-associated differences in the representation of reconstructed microbial populations, particularly during prolonged mesocosm cultivation. However, the absence of replicate metagenomic datasets precludes statistical inference about preferential recruitment or differential abundance of individual MAGs.

Together, engineered extracellular material environments leave a persistent ecological imprint on microbial community assembly, even as succession progressively reorganizes community composition (Fig. 5a–f; Supplementary Fig. S14a–d). Material environment remained significantly associated with community composition at every examined time point, explaining approximately 48% of the compositional variation after 21 days in the WWTP and 40% after 100 days in the mesocosm (Fig. 5e,f). Prolonged succession therefore did not erase material-associated community differentiation, even as alpha-diversity values became more similar (Supplementary Fig. S13c,d). Particularly revealing is the divergent behavior of ODN- and DNA hydrogel-functionalized environments: both introduce DNA into the extracellular setting, yet they are associated with distinguishable community trajectories (Supplementary Fig. S14a,b) and functional gene profiles (Supplementary Fig. S15). Unlike ODN functionalization, the DNA hydrogel presents a highly concentrated, spatially organized three-dimensional DNA matrix, suggesting that the ecological consequences of extracellular DNA depend not only on its presence but also on its local abundance and material organization. Taxonomic profiles and the relative representation of reconstructed microbial populations provide complementary, descriptive evidence of material-associated community differences (Fig. 5g; Supplementary Fig. S16). Hence, these findings establish extracellular material presentation as an important determinant of microbial community assembly and identify concentrated, three-dimensional DNA matrices as a distinctive ecological interface through which engineered DNA materials may influence microbial selection.

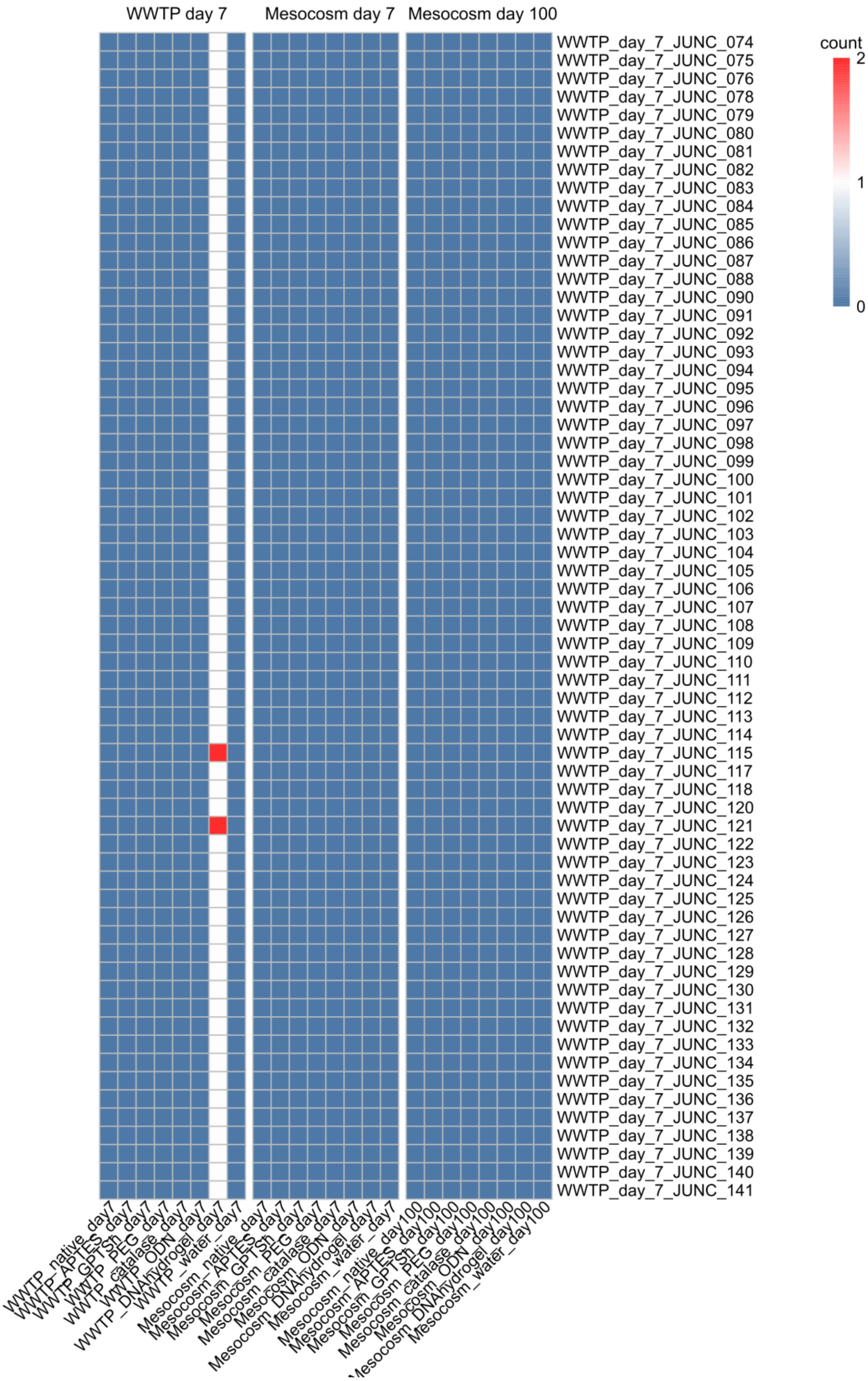

WWTP day 7
Mesocosm day 7
Mesocosm day 100
count
2
1
0
WWTP_day_7_JUNC_074
WWTP_day_7_JUNC_075
WWTP_day_7_JUNC_076
WWTP_day_7_JUNC_078
WWTP_day_7_JUNC_079
WWTP_day_7_JUNC_080
WWTP_day_7_JUNC_081
WWTP_day_7_JUNC_082
WWTP_day_7_JUNC_083
WWTP_day_7_JUNC_084
WWTP_day_7_JUNC_085
WWTP_day_7_JUNC_086
WWTP_day_7_JUNC_087
WWTP_day_7_JUNC_088
WWTP_day_7_JUNC_090
WWTP_day_7_JUNC_091
WWTP_day_7_JUNC_092
WWTP_day_7_JUNC_093
WWTP_day_7_JUNC_094
WWTP_day_7_JUNC_095
WWTP_day_7_JUNC_096
WWTP_day_7_JUNC_097
WWTP_day_7_JUNC_098
WWTP_day_7_JUNC_099
WWTP_day_7_JUNC_100
WWTP_day_7_JUNC_101
WWTP_day_7_JUNC_102
WWTP_day_7_JUNC_103
WWTP_day_7_JUNC_104
WWTP_day_7_JUNC_105
WWTP_day_7_JUNC_106
WWTP_day_7_JUNC_107
WWTP_day_7_JUNC_108
WWTP_day_7_JUNC_109
WWTP_day_7_JUNC_110
WWTP_day_7_JUNC_111
WWTP_day_7_JUNC_112
WWTP_day_7_JUNC_113
WWTP_day_7_JUNC_114
WWTP_day_7_JUNC_115
WWTP_day_7_JUNC_117
WWTP_day_7_JUNC_118
WWTP_day_7_JUNC_120
WWTP_day_7_JUNC_121
WWTP_day_7_JUNC_122
WWTP_day_7_JUNC_123
WWTP_day_7_JUNC_124
WWTP_day_7_JUNC_125
WWTP_day_7_JUNC_126
WWTP_day_7_JUNC_127
WWTP_day_7_JUNC_128
WWTP_day_7_JUNC_129
WWTP_day_7_JUNC_130
WWTP_day_7_JUNC_131
WWTP_day_7_JUNC_132
WWTP_day_7_JUNC_133
WWTP_day_7_JUNC_134
WWTP_day_7_JUNC_135
WWTP_day_7_JUNC_136
WWTP_day_7_JUNC_137
WWTP_day_7_JUNC_138
WWTP_day_7_JUNC_139
WWTP_day_7_JUNC_140
WWTP_day_7_JUNC_141
WWTP_native_day7
WWTP_APTES_day7
WWTP_GPTSh_day7
WWTP_PEG_day7
WWTP_catalase_day7
WWTP_ODN_day7
WWTP_DNAhydrogel_day7
WWTP_water_day7
Mesocosm_native_day7
Mesocosm_APTES_day7
Mesocosm_GPTSh_day7
Mesocosm_PEG_day7
Mesocosm_catalase_day7
Mesocosm_ODN_day7
Mesocosm_DNAhydrogel_day7
Mesocosm_water_day7
Mesocosm_native_day100
Mesocosm_APTES_day100
Mesocosm_GPTSh_day100
Mesocosm_PEG_day100
Mesocosm_catalase_day100
Mesocosm_ODN_day100
Mesocosm_DNAhydrogel_day100
Mesocosm_water_day100

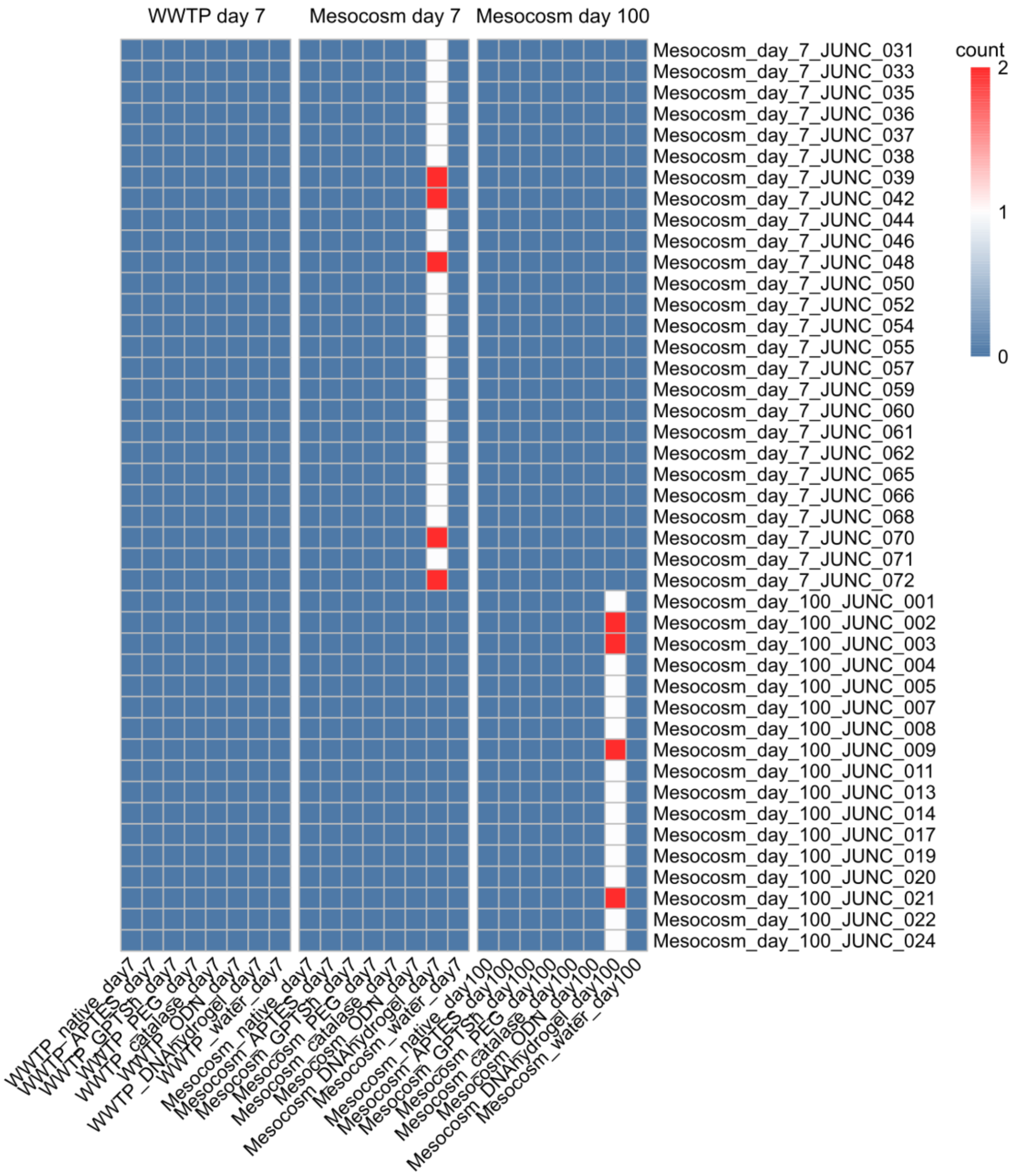


**Fig. S17 | Distribution and read support of candidate DNA hydrogel–microbial DNA junctions across environmental samples.** Heatmaps showing the number of independent read pairs supporting candidate DNA hydrogel–microbial DNA junctions in metagenomic datasets obtained after 7 days of cultivation in the wastewater treatment plant (WWTP) and after 7 and 100 days in the wastewater-derived mesocosm. The two heatmap sections display candidates initially identified in the mesocosm and WWTP datasets, respectively. Rows represent unique sequence-defined candidate junctions grouped by their dataset of initial identification, whereas columns represent the different material environments (native, APTES, GPTSh, PEG, catalase, ODN and DNA hydrogel-functionalized MESIF) and corresponding water controls. Values indicate the number of independent, non-duplicate read pairs supporting each candidate following breakpoint-specific rescreening across the metagenomic datasets. Rescreening required at least 25 aligned bases on either side of the candidate breakpoint, with 100% sequence identity across these regions. Blue indicates no supporting reads, white indicates one supporting read pair and red indicates two supporting read pairs. Candidate junctions passing the final filtering criteria were detected exclusively in DNA hydrogel-associated metagenomes, generally with low read support. No candidate junction was independently redetected across the different environmental datasets under the applied rescreening criteria. Junction identifiers indicate the dataset and sampling time of initial identification. Detailed alignment statistics, breakpoint information and microbial contig assignments are provided in Supplementary Table S5, and corresponding junction sequences are listed in Supplementary Table S6.

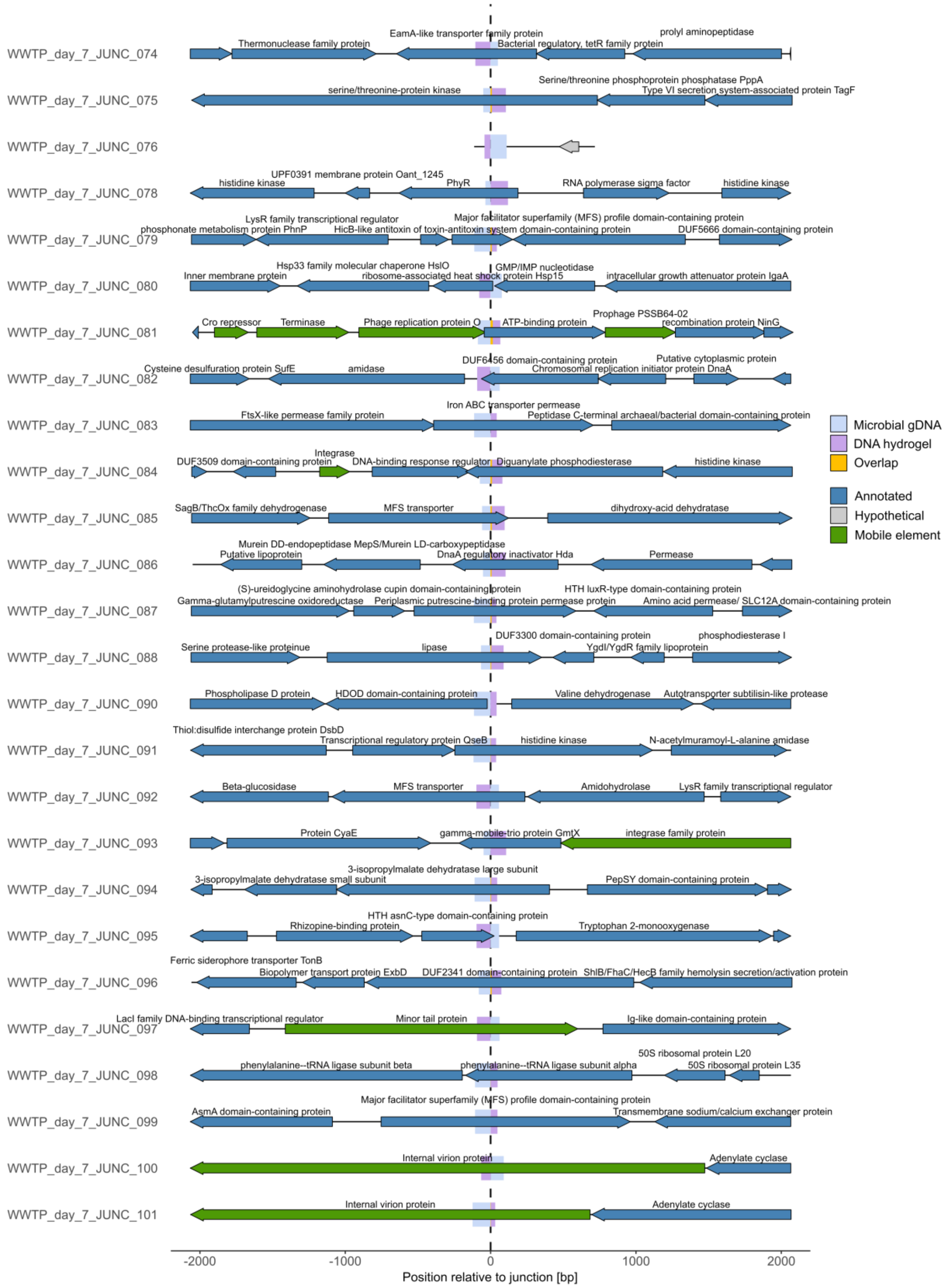
WWTP_day_7_JUNC_074
EamA-like transporter family protein
Thermonuclease family protein
Bacterial regulatory, tetR family protein
prolyl aminopeptidase
WWTP_day_7_JUNC_075
Serine/threonine phosphoprotein phosphatase PppA
serine/threonine-protein kinase
Type VI secretion system-associated protein TagF
WWTP_day_7_JUNC_076
WWTP_day_7_JUNC_078
UPF0391 membrane protein Oant_1245
histidine kinase
PhyR
RNA polymerase sigma factor
histidine kinase
WWTP_day_7_JUNC_079
LysR family transcriptional regulator
Major facilitator superfamily (MFS) profile domain-containing protein
phosphonate metabolism protein PhnP
HicB-like antitoxin of toxin-antitoxin system domain-containing protein
DUF5666 domain-containing protein
WWTP_day_7_JUNC_080
Hsp33 family molecular chaperone HslO
GMP/IMP nucleotidase
Inner membrane protein
ribosome-associated heat shock protein Hsp15
intracellular growth attenuator protein IgaA
WWTP_day_7_JUNC_081
Prophage PSSB64-02
Cro repressor
Terminase
Phage replication protein O
ATP-binding protein
recombination protein NinG
WWTP_day_7_JUNC_082
DUF6456 domain-containing protein
Putative cytoplasmic protein
Cysteine desulfuration protein SufE
amidase
Chromosomal replication initiator protein DnaA
WWTP_day_7_JUNC_083
Iron ABC transporter permease
FtsX-like permease family protein
Peptidase C-terminal archaeal/bacterial domain-containing protein
WWTP_day_7_JUNC_084
Integrase
DUF3509 domain-containing protein
DNA-binding response regulator
Diguanylate phosphodiesterase
histidine kinase
WWTP_day_7_JUNC_085
SagB/ThcOx family dehydrogenase
MFS transporter
dihydroxy-acid dehydratase
WWTP_day_7_JUNC_086
Murein DD-endopeptidase MepS/Murein LD-carboxypeptidase
Putative lipoprotein
DnaA regulatory inactivator Hda
Permease
WWTP_day_7_JUNC_087
(S)-ureidoglycine aminohydrolase cupin domain-containing protein
HTH luxR-type domain-containing protein
Gamma-glutamylputrescine oxidoreductase
Periplasmic putrescine-binding protein permease protein
Amino acid permease/ SLC12A domain-containing protein
WWTP_day_7_JUNC_088
DUF3300 domain-containing protein
phosphodiesterase I
Serine protease-like proteinue
lipase
YgdI/YgdR family lipoprotein
WWTP_day_7_JUNC_090
Phospholipase D protein
HDOD domain-containing protein
Valine dehydrogenase
Autotransporter subtilisin-like protease
WWTP_day_7_JUNC_091
Thiol:disulfide interchange protein DsbD
Transcriptional regulatory protein QseB
histidine kinase
N-acetylmuramoyl-L-alanine amidase
WWTP_day_7_JUNC_092
Beta-glucosidase
MFS transporter
Amidohydrolase
LysR family transcriptional regulator
WWTP_day_7_JUNC_093
Protein CyaE
gamma-mobile-trio protein GmtX
integrase family protein
WWTP_day_7_JUNC_094
3-isopropylmalate dehydratase large subunit
3-isopropylmalate dehydratase small subunit
PepSY domain-containing protein
WWTP_day_7_JUNC_095
HTH asnC-type domain-containing protein
Rhizopine-binding protein
Tryptophan 2-monooxygenase
WWTP_day_7_JUNC_096
Ferric siderophore transporter TonB
Biopolymer transport protein ExbD
DUF2341 domain-containing protein
ShlB/FhaC/HecB family hemolysin secretion/activation protein
WWTP_day_7_JUNC_097
LacI family DNA-binding transcriptional regulator
Minor tail protein
Ig-like domain-containing protein
WWTP_day_7_JUNC_098
50S ribosomal protein L20
phenylalanine--tRNA ligase subunit beta
phenylalanine--tRNA ligase subunit alpha
50S ribosomal protein L35
WWTP_day_7_JUNC_099
Major facilitator superfamily (MFS) profile domain-containing protein
AsmA domain-containing protein
Transmembrane sodium/calcium exchanger protein
WWTP_day_7_JUNC_100
Internal virion protein
Adenylate cyclase
WWTP_day_7_JUNC_101
Internal virion protein
Adenylate cyclase
-2000
-1000
0
1000
2000
Position relative to junction [bp]
Microbial gDNA
DNA hydrogel
Overlap
Annotated
Hypothetical
Mobile element

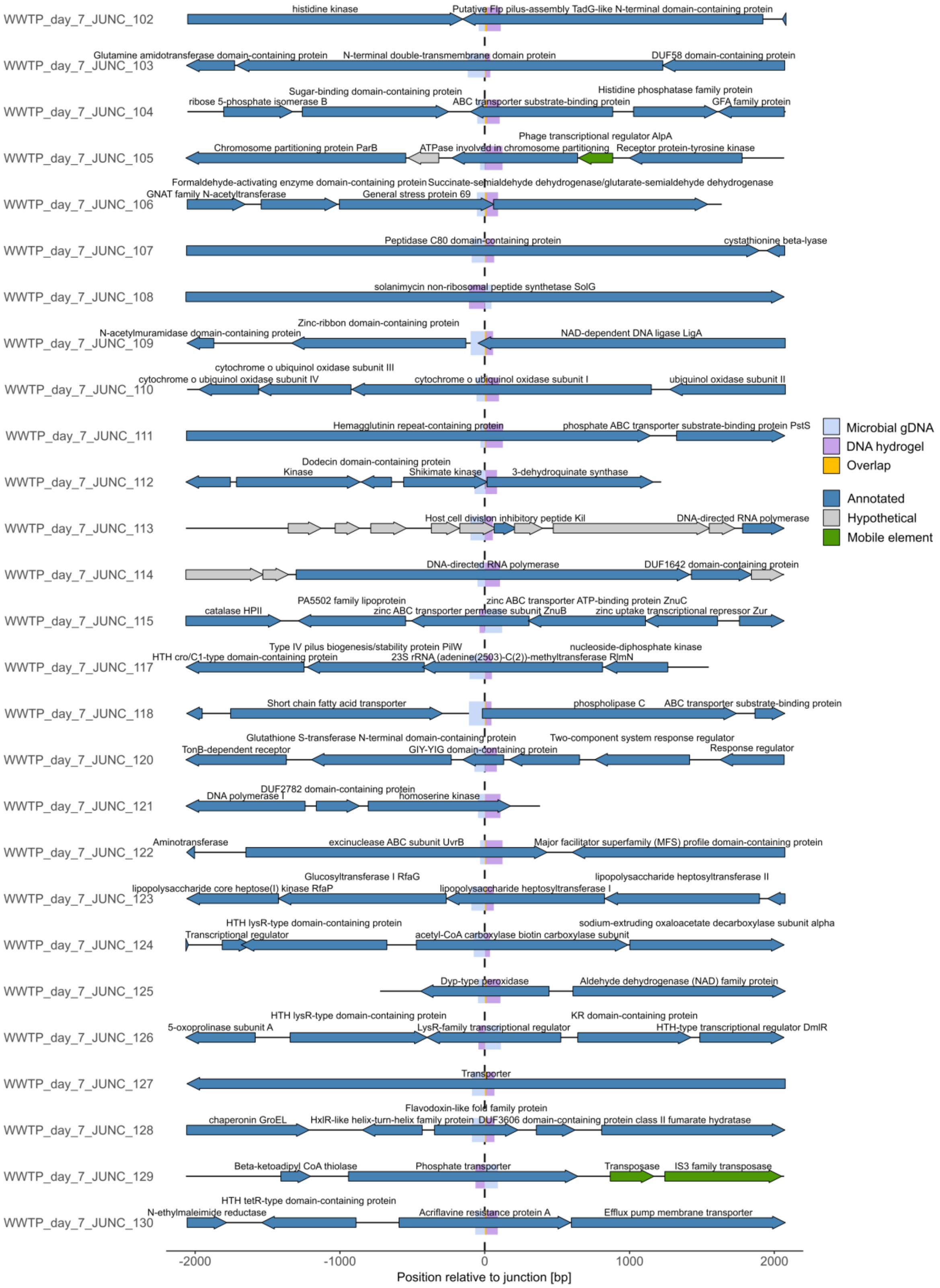
WWTP_day_7_JUNC_102
histidine kinase
Putative Flp pilus-assembly TadG-like N-terminal domain-containing protein
WWTP_day_7_JUNC_103
Glutamine amidotransferase domain-containing protein
N-terminal double-transmembrane domain protein
DUF58 domain-containing protein
WWTP_day_7_JUNC_104
Sugar-binding domain-containing protein
ribose 5-phosphate isomerase B
ABC transporter substrate-binding protein
Histidine phosphatase family protein
GFA family protein
WWTP_day_7_JUNC_105
Phage transcriptional regulator AlpA
Chromosome partitioning protein ParB
ATPase involved in chromosome partitioning
Receptor protein-tyrosine kinase
WWTP_day_7_JUNC_106
Formaldehyde-activating enzyme domain-containing protein
Succinate-semialdehyde dehydrogenase/glutarate-semialdehyde dehydrogenase
GNAT family N-acetyltransferase
General stress protein 69
WWTP_day_7_JUNC_107
Peptidase C80 domain-containing protein
cystathionine beta-lyase
WWTP_day_7_JUNC_108
solanimycin non-ribosomal peptide synthetase SolG
WWTP_day_7_JUNC_109
Zinc-ribbon domain-containing protein
N-acetylmuramidase domain-containing protein
NAD-dependent DNA ligase LigA
WWTP_day_7_JUNC_110
cytochrome o ubiquinol oxidase subunit III
cytochrome o ubiquinol oxidase subunit IV
cytochrome o ubiquinol oxidase subunit I
ubiquinol oxidase subunit II
WWTP_day_7_JUNC_111
Hemagglutinin repeat-containing protein
phosphate ABC transporter substrate-binding protein PstS
WWTP_day_7_JUNC_112
Dodecin domain-containing protein
Kinase
Shikimate kinase
3-dehydroquinate synthase
WWTP_day_7_JUNC_113
Host cell division inhibitory peptide Kil
DNA-directed RNA polymerase
WWTP_day_7_JUNC_114
DNA-directed RNA polymerase
DUF1642 domain-containing protein
WWTP_day_7_JUNC_115
PA5502 family lipoprotein
zinc ABC transporter ATP-binding protein ZnuC
catalase HPII
zinc ABC transporter permease subunit ZnuB
zinc uptake transcriptional repressor Zur
WWTP_day_7_JUNC_117
Type IV pilus biogenesis/stability protein PilW
nucleoside-diphosphate kinase
HTH cro/C1-type domain-containing protein
23S rRNA (adenine(2503)-C(2))-methyltransferase RlmN
WWTP_day_7_JUNC_118
Short chain fatty acid transporter
phospholipase C
ABC transporter substrate-binding protein
WWTP_day_7_JUNC_120
Glutathione S-transferase N-terminal domain-containing protein
Two-component system response regulator
TonB-dependent receptor
GIY-YIG domain-containing protein
Response regulator
WWTP_day_7_JUNC_121
DUF2782 domain-containing protein
DNA polymerase I
homoserine kinase
WWTP_day_7_JUNC_122
Aminotransferase
excinuclease ABC subunit UvrB
Major facilitator superfamily (MFS) profile domain-containing protein
WWTP_day_7_JUNC_123
Glucosyltransferase I RfaG
lipopolysaccharide heptosyltransferase II
lipopolysaccharide core heptose(I) kinase RfaP
lipopolysaccharide heptosyltransferase I
WWTP_day_7_JUNC_124
HTH lysR-type domain-containing protein
sodium-extruding oxaloacetate decarboxylase subunit alpha
Transcriptional regulator
acetyl-CoA carboxylase biotin carboxylase subunit
WWTP_day_7_JUNC_125
Dyp-type peroxidase
Aldehyde dehydrogenase (NAD) family protein
WWTP_day_7_JUNC_126
HTH lysR-type domain-containing protein
KR domain-containing protein
5-oxoprolinase subunit A
LysR-family transcriptional regulator
HTH-type transcriptional regulator DmlR
WWTP_day_7_JUNC_127
Transporter
WWTP_day_7_JUNC_128
Flavodoxin-like fold family protein
chaperonin GroEL
HxlR-like helix-turn-helix family protein
DUF3606 domain-containing protein
class II fumarate hydratase
WWTP_day_7_JUNC_129
Beta-ketoadipyl CoA thiolase
Phosphate transporter
Transposase
IS3 family transposase
WWTP_day_7_JUNC_130
HTH tetR-type domain-containing protein
N-ethylmaleimide reductase
Acriflavine resistance protein A
Efflux pump membrane transporter
-2000
-1000
0
1000
2000
Position relative to junction [bp]
Microbial gDNA
DNA hydrogel
Overlap
Annotated
Hypothetical
Mobile element

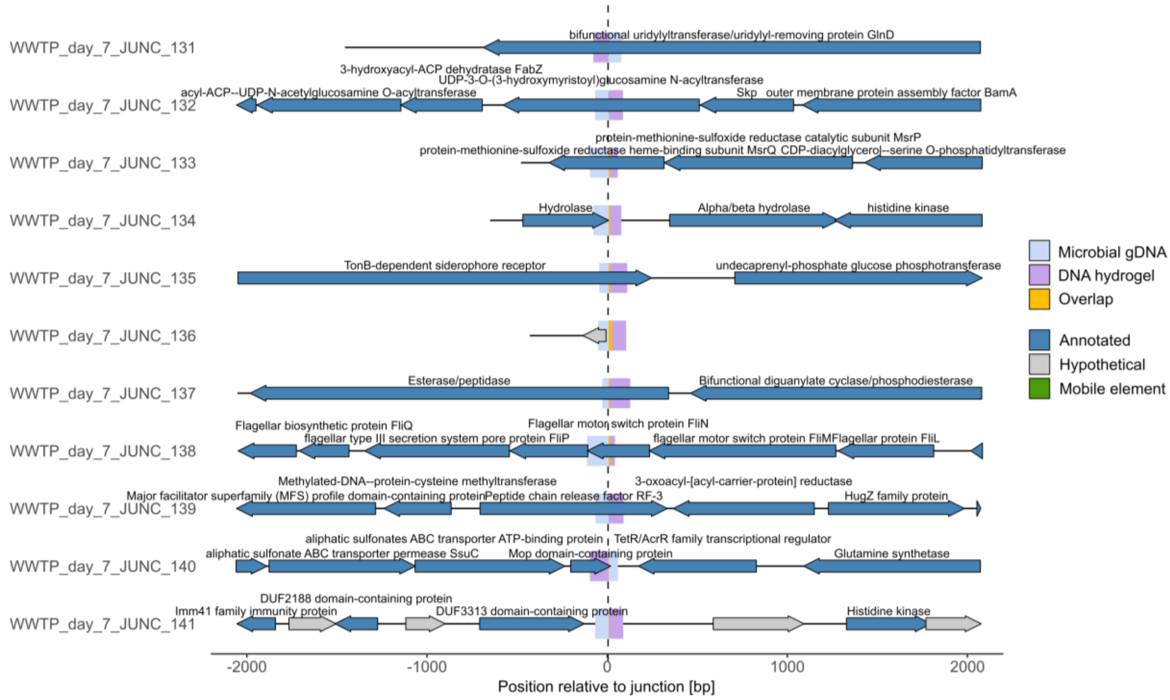


**Fig. S18 | Microbial genomic contexts associated with candidate DNA hydrogel–microbial DNA junctions detected after 7 days in the WWTP.** Genomic organization within ±2 kb of the mapped microbial breakpoints associated with individual candidate DNA hydrogel–microbial DNA junctions identified in wastewater treatment plant (WWTP) metagenomes after 7 days of cultivation. Each row represents an individual candidate junction, with genomic coordinates expressed relative to the mapped microbial breakpoint at 0 bp. The microbial portion of each junction-supporting read was aligned to the corresponding assembled microbial contig, with alignment coordinates, CIGAR strings, mapping qualities and breakpoint positions provided in Supplementary Table S5 and the corresponding junction sequences in Supplementary Table S6. Gene arrows indicate the positions and orientations of predicted coding sequences annotated using Bakta, with additional functional assignments from eggNOG-mapper and BLASTp. Annotated genes are shown in blue, hypothetical proteins in grey, and genes classified as mobile-element- or phage-associated (see Methods) in green. Colored regions around the mapped breakpoint illustrate the alignment structure of the junction-supporting read: light blue indicates the read segment aligning to the microbial contig, lilac indicates the segment aligning to the DNA hydrogel template and projected onto the microbial coordinate axis according to its position within the read, and pale yellow indicates sequence positions shared by the microbial and hydrogel-template alignments. These colored regions describe reference-alignment relationships within the sequencing read and do not independently demonstrate incorporation of hydrogel-derived DNA into the assembled microbial contig. The displayed microbial alignment corresponds to its mapped interval on the contig. Candidate junctions were associated with diverse genomic contexts, including genes involved in metabolism, transport and regulation, as well as hypothetical proteins. Mobile-element- or phage-associated genes occurred within the displayed genomic neighborhood of a subset of candidates; their proximity to the mapped microbial breakpoint does not establish a role in junction formation or DNA acquisition.

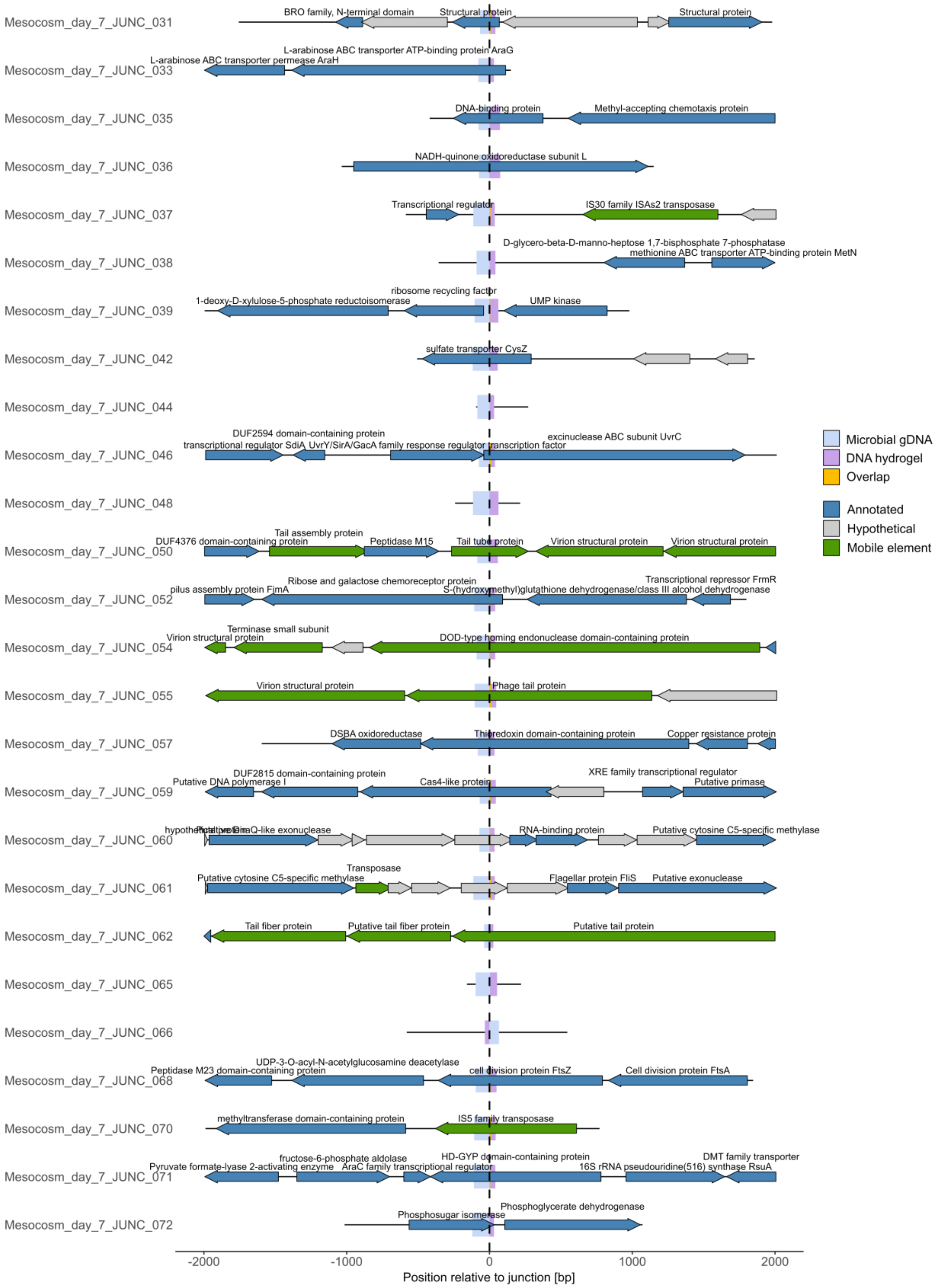


**Fig. S19 | Microbial genomic contexts associated with candidate DNA hydrogel–microbial DNA junctions detected after 7 days in the mesocosm.** Genomic organization within ±2 kb of the mapped microbial breakpoints associated with individual candidate DNA hydrogel–microbial DNA junctions identified in wastewater-derived mesocosm metagenomes after 7 days of cultivation. Each row represents an individual candidate junction, with genomic coordinates expressed relative to the mapped microbial breakpoint at 0 bp. The microbial portion of each junction-supporting read was aligned to the corresponding assembled microbial contig, with alignment coordinates, CIGAR strings, mapping qualities and breakpoint positions provided in Supplementary Table S5 and the corresponding junction sequences in Supplementary Table S6. Gene arrows indicate the positions and orientations of predicted coding sequences annotated using Bakta, with additional functional assignments from eggNOG-mapper and BLASTp. Annotated genes are shown in blue, hypothetical proteins in grey, and genes classified as mobile-element- or phage-associated (see Methods) in green. Colored regions around the mapped breakpoint illustrate the alignment structure of the junction-supporting read: light blue indicates the segment aligned to the microbial contig, lilac indicates the segment aligned to the DNA hydrogel template and projected onto the microbial coordinate axis according to its position within the read, and pale yellow indicates sequence positions shared by the microbial and hydrogel-template alignments in read coordinates. These colored regions describe reference-alignment relationships within the sequencing read and do not independently demonstrate incorporation of hydrogel-derived DNA into the assembled microbial contig. The displayed microbial alignment corresponds to its mapped interval on the contig. Candidate junctions were associated with diverse genomic contexts, including genes involved in metabolism, transport and regulation, as well as hypothetical proteins. Mobile-element- or phage-associated genes occurred within the displayed genomic neighborhood of a subset of candidates, including regions containing transposase-, phage-structural-protein- or endolysin-associated annotations. The proximity of these genes to a mapped microbial breakpoint does not establish a role in junction formation or DNA acquisition.

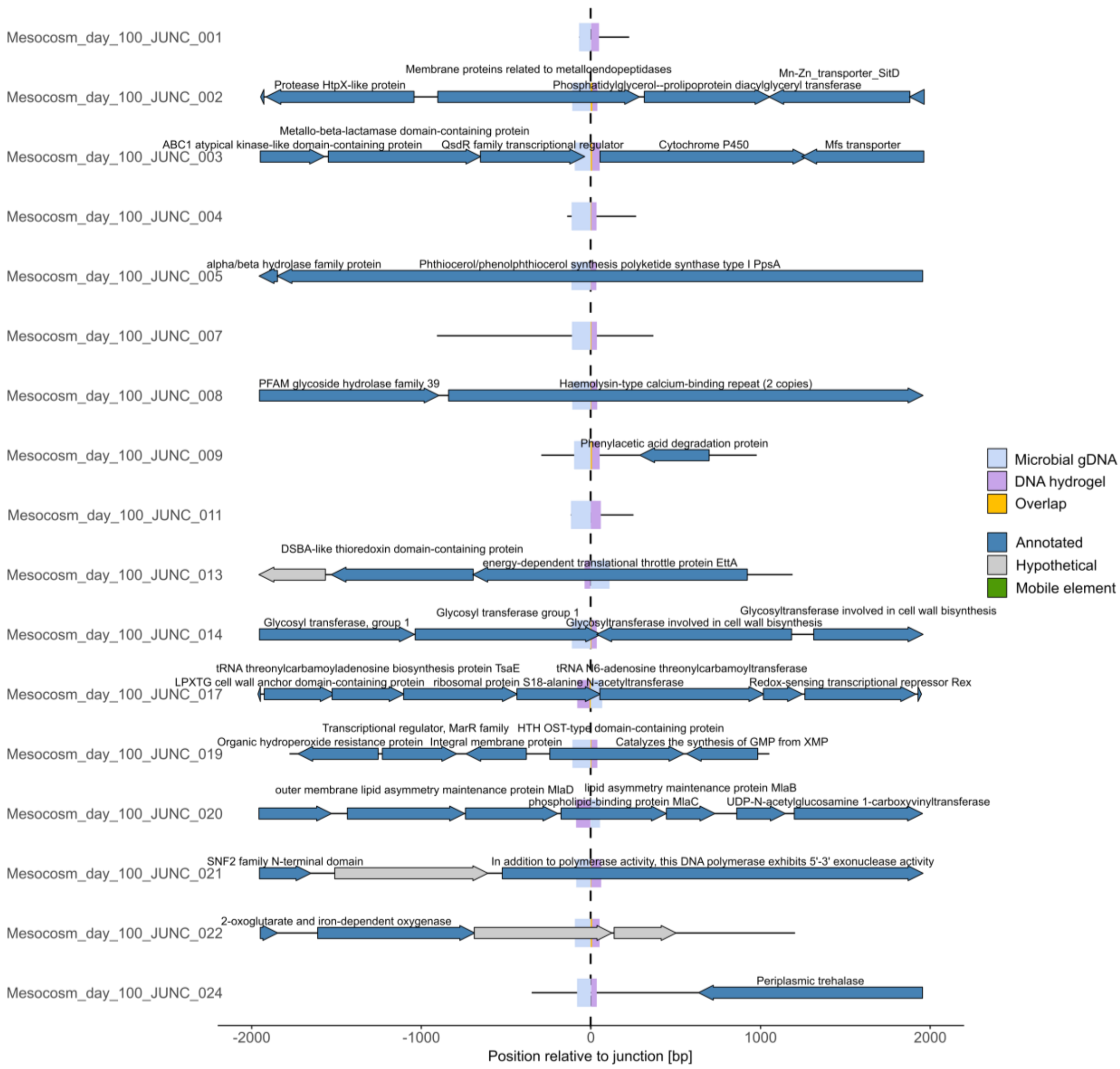


**Fig. S20 | Microbial genomic contexts associated with candidate DNA hydrogel–microbial DNA junctions detected after 100 days in the mesocosm.** Genomic organization within ±2 kb of the mapped microbial breakpoints associated with individual candidate DNA hydrogel–microbial DNA junctions identified in wastewater-derived mesocosm metagenomes after 100 days of cultivation. Each row represents an individual candidate junction, with genomic coordinates expressed relative to the mapped microbial breakpoint at 0 bp. For assembled contigs shorter than 4 kb, the available contig sequence is displayed; for longer contigs, the region spanning ±2 kb around the mapped breakpoint is shown. The microbial portion of each junction-supporting read was aligned to the corresponding assembled microbial contig, with alignment coordinates, CIGAR strings, mapping qualities and breakpoint positions provided in Supplementary Table S5 and the corresponding junction sequences in Supplementary Table S6. Gene arrows indicate the positions and orientations of predicted coding sequences annotated using Bakta, with additional functional assignments from eggNOG-mapper and BLASTp. Annotated genes are shown in blue, hypothetical proteins in grey, and genes classified as mobile-element- or phage-associated (see Methods) in green. Colored regions around the mapped breakpoint illustrate the alignment structure of the junction-supporting read: light blue indicates the segment aligned to the microbial contig, lilac indicates the segment aligned to the DNA hydrogel template and projected onto the microbial coordinate axis according to its position within the read, and pale yellow indicates sequence positions shared by the microbial and hydrogel-template alignments in read coordinates. These colored regions describe reference-alignment relationships within the sequencing read and do not independently demonstrate incorporation of hydrogel-derived DNA into the assembled microbial contig. The displayed microbial alignment corresponds to its mapped interval on the contig. Candidate junctions were associated with diverse genomic contexts, including genes involved in transport, metabolism, cellular regulation and other annotated functions, as well as hypothetical proteins. No genes classified as mobile-element- or phage-associated were identified within the displayed genomic neighborhoods of these day-100 candidates.

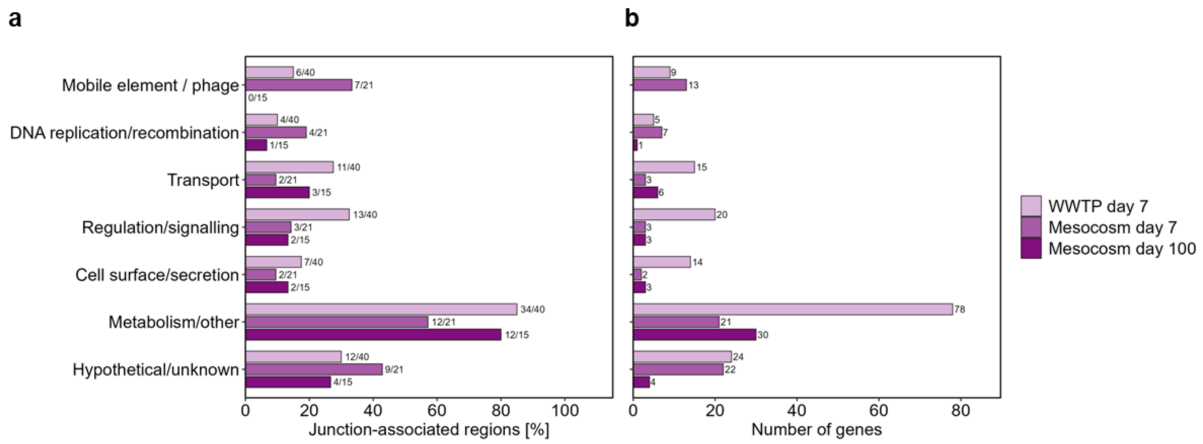


**Fig. S21 | Functional composition of microbial genomic regions associated with candidate DNA hydrogel–microbial DNA junctions.** Functional classification of genes located within ±2 kb of microbial breakpoint positions associated with candidate DNA hydrogel–microbial DNA junctions identified after 7 days in the wastewater treatment plant (WWTP) and after 7 and 100 days in the wastewater-derived mesocosm. The analysis summarizes the breakpoint-associated genomic regions shown individually in Figs. S18–S20. **(a)** Junction-level prevalence of functional categories. Bars indicate the percentage of candidate breakpoint-associated regions containing at least one gene assigned to the respective category. Numbers above bars indicate the number of positive regions relative to the total number of regions analyzed in each dataset (64, 26 and 17 for WWTP day 7, mesocosm day 7 and mesocosm day 100, respectively). Each region was counted only once per category regardless of the number of matching genes. **(b)** Gene-level functional composition of the same genomic regions, showing the total number of genes assigned to each category in the three datasets. Genes could contribute to more than one functional category; consequently, category counts are not necessarily mutually exclusive. Categories include mobile-element- or phage-associated functions, DNA replication/recombination, transport, regulation/signalling, cell surface/secretion, metabolism/other, and hypothetical/unknown functions. Genomic neighborhoods surrounding candidate breakpoints were functionally heterogeneous, and absolute gene counts should not be interpreted as evidence of functional enrichment. The proximity of annotated genes to breakpoint locations does not demonstrate involvement in junction formation, DNA acquisition, integration or transfer.

### 3. Supplementary Tables

*Table S1: Sequences and their modifications of all used oligonucleotides*

| | Name | Sequence (5'→3') | Modification |
|---|---|---|---|
| ODN | Amino modified ODN | Amine-$C_{12}$-$T_8$-$(CAT)_8$ | |
| | Complementary ODN | $(ATG)_8$ | |
| | Amino modified Cy3 ODN | Amine-C12-T8-$(ATG)_4$-Cy3 | |
| | Complementary Cy5 ODN | Cy5-$(CAT)_4$ | |
| DNA hydrogel[9] | Template A | GTA GGT GGA CAC GCC GCA TCC TAC ATC CAG CGT CAACGT CGG ACG CGA TTC ATT GAC CAG TCA GCC GCC ACC | Phosphate 5' |
| | Primer A | CCA CCT ACG GTG GCG GCT GA | |
| | A forward | GTG GCG GCT GAC TGG TCA ATG | |
| | A reverse | GGA CAC GCC GCA TCC TAC ATC | |
| | A probe | CGC GTC CGA CGT TGA CGC | FAM 5' TAMRA 3' |

*Table S2: Gene constructs and proteins used*

| | |
|---|---|
| His-SC[7] | ATGAGCTATTATCATCATCACCATCACCACGATTATGATATTCCGACCA<br>CCGAAAATCTGTATTTTCAGGGTGCAATGGTTGATACCCTGAGCGGTC<br>TGAGCAGCGAACAGGGTCAGAGCGGTGATATGACCATTGAAGAAGATA<br>GCGCAACCCACATCAAATTCAGCAAACGTGATGAAGATGGTAAAGAAC<br>TGGCAGGCGCAACCATGGAACTGCGTGATAGCAGCGGTAAAACCATTA<br>GCACCTGGATTAGTGATGGTCAGGTGAAAGATTTTTATCTGTACCCTG<br>GCAAATACACCTTTGTTGAAACCGCAGCACCGGATGGTTATGAAGTTG<br>CAACCGCAATTACCTTTACCGTTAATGAACAGGGCCAGGTTACCGTGA<br>ATGGTAAAGCAACCAAAGGTGATGCACATATTTAA |
| TaCAT-ST-His<br>*Thermoanaerobaculum aquaticum* | ATGAGCGAGAAACAGGGTCAGCAGAAACCGGTTCAGCGTGGTGCACTG<br>CGTCATACCGTTCTGGGTCATCTGGGTATTCGTCAGTGGTGGCCTCAGC<br>AGCTGAATCTGGGCATTCTGCATCAGCATAGCGAAAAAAGCAATCCGC<br>TGGGTCGTGATTTTAACTATCGTAAAGCATTTAGCGAGCTGGATTATT<br>GGGCACTGAAAAAAGATCTGGCAGAACTGATGACACAGAGCCAAGAAT<br>GGTGGCCTGCAGATTGGGGTCATTATGGTGGTCTGATGATTCGTATGG<br>CATGGCATAGCGCAGGCACCTATCGTATTTTTGATGGTCGTGGTGGTG<br>GTAGTACCGGTGCACAGCGTTTTATGCCGTTAGGTAGCTGGCCTGATA<br>ATGCAAATCTGGATAAAGCACGTCGTCTGCTGTGGCCGATTAAAAAGA<br>AATATGGCAACAAAATCAGCTGGGCTGATCTGATGATCCTGGCAGGTA<br>ATGTTGCACTGGAAACCATGGGCTTTAAAACCTTAGGCTTTGGTGGTG<br>GCCGTGAAGATGTTTGGGAACCGCCTGAAGATATTGATTGGGGACCTG<br>AAACCGAATGGCTGGGTGATCAGCGTCATAGCGGTCATCGTGAACTGG<br>CAAAACCGTATGCAGCAGTTCAGATGGGTCTGATTTATGTTAATCCGG<br>AAGGTCCTGAAGGTAAACCGGATCCTGTTGCAGCAGGTAAAGAAATTC<br>GTGAAAGCTTTGCCCGTATGGGCATGAATGATGAAGAAACCGTTGCAC<br>TGATTGTTGGTGGTCATACCTTTGGTAAAGCACATGGTGCAGTTCCGG<br>CAAGCCATGTTGGTCCGGAACCGGAAGCAGCACCGCTGGAACAGGTTG<br>GTTTAGGTTGGAAAAATACCGCAGGCAAAGGTCATAGCGAACATACCT<br>ATACCAGCGGTCTGGAAGGTGCATGGACCCCGACACCGACCAAATGGG<br>ATAATACCTTTCTGGAAATCCTGTTCAGCTATGAATGGGAGTTAGTTA<br>AAAGTCCGGCAGGCGCATGGCAGTGGCATCCGAAAAATGTTCGTGAAG<br>AACATATGGTTCCGGATGCACATGTTCCGGGTAAAAAGAATCCTCCGT<br>TTATGCTGACCACCGATCTGGCACTGCGTTTTGATCCGATTTATGAACC<br>GATTGCACGTCGTTTTTGGCAGAATCCGAAAGAACTGGAAGAGGCATT<br>TGCAAAAGCATGGTTTAAACTGACCCATCGTGATATGGGTCCGAAAAG<br>CCGTTATCTGGGTCCAGAAGTGCCGAAAGAAGATTTTGTTTGGCAGGA<br>TCCTCTGCCTCCGGTTGAAGGTGCTCCGCTGGGCAAAGCAGATATTCAG<br>ACCCTGAAACAGAAAATTCTGGCAAGCGGTGTTACTCCGCGTGATCTG<br>GTTTATGTGGCATGGTCAGCAGCAAGTACCTTTCGTATTAGCGATAAA<br>CGCGGTGGTGCAAATGGTGCACGTATTCGCCTGCTGCCGCAGAAAAAT<br>TGGCCTGTTAATGAACCGGAAACACTGGCCAAAGTTCTGGCAGCCCTG<br>GAAAAAATCAAAGGTGAATTTGAAGCAGCCAGCGGTAAAAAAGTTAGC<br>CTGGCCGATCTGATTGTTTTAGGTGGCACCGCAGCAGTTGAAGAAGCA<br>GCCCGTCAGGCAGGTTTTGCCGTTGAAGTTCCGTTTTTTCCGGGTCGTG<br>CAGATGCAAGCCAAGAACAGACCGAAGTTGATTTTTATGCAGTGATGG<br>AACCTGAAGCCGATGGTTTTCGTAATTTTCAGAAAGAGGATAGCGTTA<br>TCCCGCTGGAAGAAGCCCTGCTGGACAAAGCACAGCTGCTGTCACTGAC<br>CGCACCGGAAATGACCGCACTGGTTGGCGGTCTGCGTGTTCTGGGTGCA<br>GTTTTTGGTAATAGCAGCCTGGGTGTTCTGACCCAGCGTGTTGGTGCCC<br>TGACCAATGATTATTTTGTTAATCTGCTGGATATGCGCTATGAGTGGC<br>AGGACCTGCATGGTGATGGCCGTTTTTTTCAGGCACGTGGTCGTAAAA<br>CCGGTGCAGCAATTTGGACCGTTACACGTGCAGATCTGGTGTTTGCAGC |

| | |
|---|---|
| | ACATAGCCAGCTGCGTGGTTATGCCGAAGTTTATGCCTGTGATGATAA<br>CCAAGAAAAATTCGTGCGTGATTTTGCAGCAGCATGGTCAAAAGTTAT<br>GCATCTGGATCGCTTTGATCTGCAGCAAGGTGGTGGTGGCAGCGCACA<br>TATTGTTATGGTTGATGCATATAAACCGACGAAAGGTCATCATCACCA<br>TCATCATTAATGA |
| **mRFP1**[10] | ATGGCAAGCAGCGAAGATGTGATCAAAGAATTTATGCGTTTCAAGGTG<br>CGTATGGAAGGTAGCGTTAATGGTCATGAATTTGAAATCGAAGGTGAA<br>GGTGAGGGTCGTCCGTATGAAGGCACCCAGACCGCCAAACTGAAAGTT<br>ACCAAAGGCGGTCCGCTGCCGTTTGCATGGGATATTCTGAGTCCGCAGT<br>TTCAGTATGGTAGCAAAGCATATGTTAAACATCCGGCAGATATCCCGG<br>ATTATCTGAAACTGAGCTTTCCGGAAGGCTTTAAATGGGAACGTGTGA<br>TGAATTTTGAGGATGGTGGTGTTGTTACCGTTACACAGGATAGCAGCC<br>TGCAGGATGGTGAATTTATCTATAAAGTTAAACTGCGTGGCACGAATT<br>TTCCGTCAGATGGTCCTGTTATGCAGAAAAAAACCATGGGTTGGGAAG<br>CAAGCACCGAACGTATGTATCCGGAAGATGGTGCACTGAAAGGTGAAA<br>TCAAAATGCGTCTGAAGCTGAAAGATGGCGGTCATTATGATGCAGAAG<br>TTAAAACCACCTACATGGCCAAAAAACCGGTGCAGCTGCCTGGTGCAT<br>ATAAAACGGATATTAAACTGGATATCACCTCGCACAACGAGGATTATA<br>CCATTGTTGAACAGTATGAACGTGCAGAAGGTCGTCATAGTACCGGTG<br>CCTAATAA |
| **GFPmt3b**[10] | ATGCGTAAAGGTGAAGAACTGTTTACCGGTGTTGTTCCGATTCTGGTT<br>GAACTGGATGGTGATGTTAATGGCCACAAATTTTCAGTTAGCGGTGAA<br>GGCGAAGGTGATGCAACCTATGGTAAACTGACCCTGAAATTTATCTGT<br>ACCACCGGCAAACTGCCGGTTCCGTGGCCGACACTGGTTACCACCTTTG<br>GTTATGGTGTTCAGTGTTTTGCACGTTATCCGGATCATATGAAACAGC<br>ACGATTTTTTCAAAAGCGCAATGCCGGAAGGTTATGTTCAAGAACGTA<br>CCATCTTCTTCAAAGATGACGGCAACTATAAAACCCGTGCCGAAGTTA<br>AATTTGAAGGTGATACCCTGGTGAATCGCATTGAACTGAAAGGCATCG<br>ATTTTAAAGAGGATGGTAATATCCTGGGCCACAAACTGGAATATAATT<br>ATAATAGCCACAACGTGTACATCATGGCCGACAAACAGAAAAATGGCA<br>TCAAAGTGAACTTCAAGATCCGCCATAATATTGAAGATGGTTCAGTTC<br>AGCTGGCCGATCATTATCAGCAGAATACCCCGATTGGTGATGGTCCGG<br>TTCTGCTGCCGGATAATCATTATCTGAGCACCCAGAGCGCACTGAGCAA<br>AGATCCGAATGAAAAACGTGATCACATGGTGCTGCTGGAATTTGTTAC<br>CGCAGCAGGTATTACCCATGGTATGGATGAACTGTATAAATGATAA |
| **eGFP**[10] | ATGGTTAGCAAAGGTGAAGAACTGTTTACCGGTGTTGTTCCGATTCTG<br>GTTGAACTGGATGGTGATGTTAATGGCCACAAATTTTCAGTTAGCGGT<br>GAAGGTGAGGGTGATGCAACCTATGGTAAACTGACCCTGAAATTTATC<br>TGCACCACCGGCAAACTGCCGGTTCCGTGGCCGACCCTGGTTACCACCC<br>TGACCTATGGTGTTCAGTGTTTTAGCCGTTATCCGGATCATATGAAAC<br>AGCACGATTTTTTCAAAAGCGCAATGCCGGAAGGTTATGTTCAAGAAC<br>GTACCATCTTTTTCAAAGACGACGGCAACTATAAAACCCGTGCCGAAG<br>TTAAATTTGAAGGTGATACCCTGGTGAATCGCATTGAACTGAAAGGCA<br>TCGATTTTAAAGAAGATGGCAATATCCTGGGCCACAAACTGGAATATA<br>ATTATAATAGCCACAACGTGTACATCATGGCCGACAAACAGAAAAATG<br>GCATCAAAGTGAACTTCAAAATCCGCCACAATATTGAAGATGGCTCAG<br>TTCAGCTGGCAGATCATTATCAGCAGAATACCCCGATCGGTGATGGTCC<br>GGTTCTGCTGCCGGATAATCATTATCTGAGCACCCAGAGCGCACTGAGC<br>AAAGATCCGAATGAAAAACGTGATCACATGGTGCTGCTGGAATTTGTT<br>ACCGCAGCAGGTATTACCCTGGGTATGGATGAACTGTATAAA |

| | |
|---|---|
| **mTagBFP2**[10] | ATGGTTAGCAAAGGCGAAGAACTGATCAAAGAAAACATGCACATGAAA<br>CTGTATATGGAAGGCACCGTTGATAACCACCACTTTAAATGTACCAGC<br>GAAGGTGAAGGTAAACCGTATGAAGGCACCCAGACCATGCGTATTAAA<br>GTTGTTGAAGGTGGTCCGCTGCCGTTTGCATTTGATATTCTGGCAACCA<br>GCTTTCTGTATGGTAGCAAAACCTTTATCAATCATACCCAGGGTATCCC<br>GGATTTTTTCAAACAGAGCTTTCCGGAAGGTTTTACCTGGGAACGTGT<br>TACCACCTATGAAGATGGTGGTGTTCTGACCGCAACACAGGATACCAG<br>TCTGCAGGATGGTTGTCTGATTTATAATGTGAAAATTCGCGGTGTGAA<br>CTTCACCAGCAATGGTCCGGTTATGCAGAAAAAGACCTTAGGTTGGGA<br>AGCATTTACCGAAACACTGTATCCGGCAGATGGTGGCCTGGAAGGTCG<br>TAATGATATGGCACTGAAACTGGTTGGTGGTAGCCATCTGATTGCAAA<br>TGCCAAAACCACCTATCGTAGCAAAAAACCGGCAAAGAATCTGAAAAT<br>GCCTGGCGTTTATTATGTGGATTATCGTCTGGAACGCATCAAAGAAGC<br>AAACAACGAAACCTATGTTGAACAGCATGAAGTTGCAGTTGCACGTTA<br>TTGTGATCTGCCGAGCAAACTGGGTCATAAACTGAACTGA |
| **Knt**<br>Aminoglycoside O-nucleotidyltransferase ANT(4')-Ia | ATGAGAATAGTGAATGGACCAATAATAATGACTAGAGAAGAAAGAAT<br>GAAGATTGTTCATGAAATTAAGGAACGAATATTGGATAAATATGGGGA<br>TGATGTTAAGGCTATTGGTGTTTATGGCTCTCTTGGTCGTCAGACTGA<br>TGGGCCCTATTCGGATATTGAGATGATGTGTGTCATGTCAACAGAGGA<br>AGCAGAGTTCAGCCATGAATGGACAACCGGTGAGTGGAAGGTGGAAGT<br>GAATTTTTATAGCGAAGAGATTCTACTAGATTATGCATCTCAGGTGGA<br>ATCAGATTGGCCGCTTACACATGGTCAATTTTTCTCTATTTTGCCGATT<br>TATGATTCAGGTGGATACTTAGAGAAAGTGTATCAAACTGCTAAATCG<br>GTAGAAGCCCAAAAGTTCCACGATGCGATTTGTGCCCTTATCGTAGAA<br>GAGCTGTTTGAATATGCAGGCAAATGGCGTAATATTCGTGTGCAAGGA<br>CCGACAACATTTCTACCATCCTTGACTGTACAGGTAGCAATGGCAGGTG<br>CCATGTTGATTGGTCTGCATCATCGCATCTGTTATACGACGAGCGCTTC<br>GGTCTTAACTGAAGCAGTTAAGCAATCAGATCTTCCTTCAGGTTATGA<br>CCATCTGTGCCAGTTCGTAATGTCTGGTCAACTTTCCGACTCTGAGAAA<br>CTTCTGGAATCGCTAGAGAATTTCTGGAATGGGATTCAGGAGTGGACA<br>GAACGACACGGATATATAGTGGATGTGTCAAAACGCATACCATTTTGA<br>TAA |

*Table S3: Fluorescence excitation and emission wavelengths*

| Fluorescent protein | Color | Maxima [nm] | Synergy H1 filters [nm] | Fluorescent imager filters [nm] | CLSM filters [nm] |
|---|---|---|---|---|---|
| mTagBFP2 | Blue | $\lambda_{Ex}$: 399<br>$\lambda_{Em}$: 454 | $\lambda_{Ex}$: 400<br>$\lambda_{Em}$:454<br>Gain: 53 | $\lambda_{Ex}$: 440<br>$\lambda_{Em}$: F-500<br>400 ms | $\lambda_{Ex}$: 405<br>$\lambda_{Em}$: 450 |
| mRFP1 | Red | $\lambda_{Ex}$: 584<br>$\lambda_{Em}$: 607 | $\lambda_{Ex}$: 571<br>$\lambda_{Em}$: 607<br>Gain: 68 | $\lambda_{Ex}$: 580<br>$\lambda_{Em}$: F-650<br>100 ms | $\lambda_{Ex}$: 561<br>$\lambda_{Em}$: 588 |
| GFPmt3b/<br>eGFP | Green | $\lambda_{Ex}$: 507/488<br>$\lambda_{Em}$: 513/507 | $\lambda_{Ex}$: 483<br>$\lambda_{Em}$: 511<br>Gain: 50 | $\lambda_{Ex}$: 480<br>$\lambda_{Em}$: F-500<br>400 ms | $\lambda_{Ex}$: 488<br>$\lambda_{Em}$: 516 |

*Table S4: MAGs found in the metagenomic data*

| MAG | Comp [%] | Con [%] | GS [Mbp] | GC Content | Phylum | Order | Class | Family | Genus | Species |
|---|---|---|---|---|---|---|---|---|---|---|
| **MAG_Meso_1** | 99.98 | 1.78 | 3.78 | 0.66 | *Pseudomonadota* | *Gammaproteobacteria* | *Burkholderiales* | *Rhodocyclaceae* | *Methyloversatilis* | *Methyloversatilis sp016791045* |
| **MAG_Meso_2** | 76.65 | 4.51 | 1.71 | 0.46 | *Pseudomonadota* | *Alphaproteobacteria* | *Micavibrionales* | *Micavibrionaceae* | | |
| **MAG_Meso_3** | 100 | 0.21 | 3.15 | 0.39 | *Bacteroidota* | *Bacteroidia* | *Chitinophagales* | *Chitinophagaceae* | *Sediminibacterium* | *Sediminibacterium salmoneum* |
| **MAG_Meso_4** | 93.2 | 3.61 | 2.16 | 0.51 | *Pseudomonadota* | *Gammaproteobacteria* | *Burkholderiales* | *Burkholderiaceae* | *Algicoccus* | |
| **MAG_Meso_5** | 67.92 | 2.99 | 1.20 | 0.43 | *Pseudomonadota* | *Gammaproteobacteria* | *Coxiellales* | *Coxiellaceae* | | |
| **MAG_Meso_6** | 100 | 0.55 | 4.35 | 0.66 | *Pseudomonadota* | *Alphaproteobacteria* | *Zavarziniales* | *Zavarziniaceae* | *Zavarzinia* | |
| **MAG_Meso_7** | 61.26 | 3.79 | 1.88 | 0.64 | *Elusimicrobiota* | *Elusimicrobia* | *Elusimicrobiales* | *UBA9959* | *UBA9959* | |
| **MAG_Meso_8** | 88.64 | 1.24 | 4.31 | 0.68 | *Verrucomicrobiota* | *Verrucomicrobiae* | *Opitutales* | *Opitutaceae* | *Didemnitutus* | |
| **MAG_Meso_9** | 87.01 | 3.99 | 2.18 | 0.4 | *Chlamydiota* | *Chlamydiia* | *Chlamydiales* | *Parachlamydiaceae* | *JACCRY01* | |
| **MAG_Meso_10** | 85.14 | 1.67 | 3.03 | 0.66 | *Pseudomonadota* | *Gammaproteobacteria* | *Burkholderiales* | *Burkholderiaceae* | *PBB1* | |
| **MAG_Meso_11** | 81.89 | 1.00 | 4.94 | 0.7 | *Planctomycetota* | *UBA1135* | *UBA2386* | *UBA2386* | *UBA2386* | |
| **MAG_Meso_12** | 88.83 | 1.64 | 1.13 | 0.46 | *Patescibacteria* | *Saccharimonadia* | *Saccharimonadales* | *JAGQNJ01* | *HK-STAS-PATE-36* | |
| **MAG_Meso_13** | 96.24 | 2.95 | 4.62 | 0.73 | *Actinomycetota* | *Thermoleophilia* | *Solirubrobacterales* | *Solirubrobacteraceae* | *SYBY01* | |
| **MAG_Meso_14** | 56.93 | 0.60 | 2.70 | 0.67 | *Pseudomonadota* | *Gammaproteobacteria* | *Xanthomonadales* | *Xanthomonadaceae* | *SCMT01* | |
| **MAG_Meso_15** | 84.44 | 0.33 | 2.79 | 0.61 | *Actinomycetota* | *Actinomycetes* | *S36-B12* | *UBA10799* | *UBA10799* | |
| **MAG_Meso_16** | 60.06 | 7.84 | 4.11 | 0.65 | *Pseudomonadota* | *Alphaproteobacteria* | *Reyranellales* | *Reyranellaceae* | *Reyranella* | *Reyranella sp017307275* |
| **MAG_Meso_17** | 53.45 | 5.05 | 2.31 | 0.53 | *Pseudomonadota* | *Gammaproteobacteria* | *Enterobacterales* | *Enterobacteriaceae* | *Escherichia* | *Escherichia coli* |
| **MAG_Meso_18** | 100 | 0.50 | 6.52 | 0.66 | *Actinomycetota* | *Actinomycetes* | *Mycobacteriales* | *Mycobacteriaceae* | *Mycobacterium* | *Mycobacterium conceptionense* |
| **MAG_Meso_19** | 99.98 | 2.36 | 5.86 | 0.67 | *Actinomycetota* | *Actinomycetes* | *Mycobacteriales* | *Mycobacteriaceae* | *Mycobacterium* | *Mycobacterium sp001644575* |
| **MAG_Meso_20** | 86.59 | 5.65 | 3.22 | 0.71 | *Actinomycetota* | *Thermoleophilia* | *Solirubrobacterales* | *70-9* | *VAYN01* | |
| **MAG_Meso_21** | 77.55 | 5.24 | 2.66 | 0.68 | *Pseudomonadota* | *Alphaproteobacteria* | *Sphingomonadales* | *Sphingomonadaceae* | *JAEZQR01* | |
| **MAG_Meso_22** | 68.02 | 5.19 | 4.57 | 0.62 | *Pseudomonadota* | *Gammaproteobacteria* | *Pseudomonadales* | *Pseudomonadaceae* | *Pseudomonas_E* | *Pseudomonas_E yamanorum* |
| **MAG_Meso_23** | 71.47 | 2.06 | 1.84 | 0.68 | *Acidobacteriota* | *Holophagae* | *Holophagales* | *Holophagaceae* | *Holophaga* | *Holophaga sp002068835* |
| **MAG_Meso_24** | 76.73 | 3.96 | 1.82 | 0.66 | *Pseudomonadota* | *Gammaproteobacteria* | *Burkholderiales* | *Burkholderiaceae* | *Macromonas* | *Macromonas bipunctata* |
| **MAG_Meso_25** | 99.68 | 0.07 | 4.15 | 0.64 | *Pseudomonadota* | *Alphaproteobacteria* | *Rhodobacterales* | *Rhodobacteraceae* | *Gemmobacter* | *Gemmobacter fulvus* |
| **MAG_Meso_26** | 94.94 | 0.82 | 5.08 | 0.65 | *Pseudomonadota* | *Gammaproteobacteria* | *Nevskiales* | *Nevskiaceae* | *Hydrocarboniphaga* | |
| **MAG_Meso_27** | 96.92 | 1.10 | 4.04 | 0.62 | *Pseudomonadota* | *Gammaproteobacteria* | *Burkholderiales* | *Rhodocyclaceae* | *Azonexus* | *Azonexus sp016790885* |

*Table S4: MAGs found in the metagenomic data*

| | | | | | | | | | | |
|---|---|---|---|---|---|---|---|---|---|---|
| **MAG_Meso_28** | 95.49 | 1.34 | 3.35 | 0.62 | *Pseudomonadota* | *Gammaproteobacteria* | *Burkholderiales* | *Rhodocyclaceae* | *Sulfuricystis* | |
| **MAG_Meso_29** | 86.51 | 4.98 | 4.26 | 0.65 | *Pseudomonadota* | *Gammaproteobacteria* | *Chromatiales* | *Chromatiaceae* | *Thiocapsa* | |
| **MAG_Meso_30** | 70.35 | 0.44 | 0.73 | 0.4 | *Bacillota_I* | *Bacilli_A* | *RFN20* | *CAG-826* | | |
| **MAG_Meso_31** | 69.6 | 5.22 | 3.37 | 0.71 | *Pseudomonadota* | *Alphaproteobacteria* | *Rhizobiales* | *Pseudoxanthobacteraceae* | *Pseudoxanthobacter* | |
| **MAG_Meso_32** | 90.85 | 1.85 | 2.35 | 0.38 | *Pseudomonadota* | *Gammaproteobacteria* | *DSM-16500* | *DSM-16500* | *BMAH01* | |
| **MAG_Meso_33** | 99.88 | 1.20 | 3.13 | 0.67 | *Pseudomonadota* | *Alphaproteobacteria* | *Caulobacterales* | *Caulobacteraceae* | *Brevundimonas* | |
| **MAG_Meso_34** | 86.93 | 0.36 | 2.53 | 0.57 | *Pseudomonadota* | *Alphaproteobacteria* | *UBA11222* | *UBA11222* | *UBA11222* | |
| **MAG_Meso_35** | 95.35 | 1.48 | 2.45 | 0.68 | *Actinomycetota* | *Actinomycetes* | *Actinomycetales* | *Microbacteriaceae* | *Microbacterium* | *Microbacterium sp011620715* |
| **MAG_Meso_36** | 95.79 | 1.97 | 5.11 | 0.71 | *Actinomycetota* | *Thermoleophilia* | *Solirubrobacterales* | *Solirubrobacteraceae* | *SYBY01* | |
| **MAG_Meso_37** | 100 | 0.94 | 7.01 | 0.66 | *Actinomycetota* | *Actinomycetes* | *Mycobacteriales* | *Mycobacteriaceae* | *Mycobacterium* | *Mycobacterium fortuitum* |
| **MAG_Meso_38** | 89.79 | 1.92 | 3.59 | 0.72 | *Actinomycetota* | *Acidimicrobiia* | *Acidimicrobiales* | *Microtrichaceae* | | |
| **MAG_Meso_39** | 58.1 | 3.61 | 3.64 | 0.61 | *Pseudomonadota* | *Alphaproteobacteria* | *Rhizobiales* | *Rhizobiaceae* | *Ectorhizobium* | |
| **MAG_Meso_41** | 94.88 | 0.01 | 2.90 | 0.64 | *Actinomycetota* | *Thermoleophilia* | *Solirubrobacterales* | *70-9* | *67-14* | |
| **MAG_Meso_42** | 95.47 | 3.93 | 6.16 | 0.66 | *Pseudomonadota* | *Alphaproteobacteria* | *Rhizobiales* | *Hyphomicrobiaceae* | *AWTP1-13* | |
| **MAG_Meso_43** | 89.45 | 2.64 | 2.46 | 0.55 | *Desulfobacterota_D* | *UBA1144* | *UBA2774* | *UBA2774* | *JADLGH01* | *JADLGH01 sp020054175* |
| **MAG_Meso_44** | 67.96 | 4.22 | 3.32 | 0.71 | *Actinomycetota* | *Actinomycetes* | *S36-B12* | *S36-B12* | *UBA10649* | |
| **MAG_Meso_45** | 95.53 | 0.02 | 2.68 | 0.63 | *Actinomycetota* | *Thermoleophilia* | *Solirubrobacterales* | *70-9* | *67-14* | *67-14 sp030917845* |
| **MAG_Meso_46** | 92.16 | 6.72 | 6.14 | 0.66 | *Actinomycetota* | *Actinomycetes* | *Mycobacteriales* | *Mycobacteriaceae* | *Mycobacterium* | *Mycobacterium lutetiense* |
| **MAG_Meso_47** | 85.9 | 0.08 | 4.77 | 0.67 | *Actinomycetota* | *Actinomycetes* | *Mycobacteriales* | *Mycobacteriaceae* | *Mycobacterium* | *Mycobacterium novocastrense* |
| **MAG_Meso_48** | 100 | 0.90 | 4.55 | 0.66 | *Pseudomonadota* | *Gammaproteobacteria* | *Xanthomonadales* | *Xanthomonadaceae* | *Stenotrophomonas* | *Stenotrophomonas maltophilia* |
| **MAG_Meso_49** | 59.08 | 4.23 | 2.51 | 0.63 | *Pseudomonadota* | *Gammaproteobacteria* | *Pseudomonadales* | *Pseudomonadaceae* | *Pseudomonas_E* | *Pseudomonas_E sp019747075* |
| **MAG_Meso_50** | 88.15 | 4.56 | 4.02 | 0.63 | *Pseudomonadota* | *Gammaproteobacteria* | *Enterobacterales* | *Aeromonadaceae* | *Aeromonas* | *Aeromonas hydrophila* |
| **MAG_Meso_51** | 97.8 | 1.60 | 4.31 | 0.53 | *Pseudomonadota* | *Gammaproteobacteria* | *Enterobacterales* | *Enterobacteriaceae* | *Citrobacter* | *Citrobacter freundii* |
| **MAG_WWTP_1** | 52.16 | 3.18 | 1.57 | 0.65 | *Desulfobacterota* | *Desulfovibrionia* | *Desulfovibrionales* | *Desulfomicrobiaceae* | *Desulfomicrobium* | *Desulfomicrobium sp024709365* |
| **MAG_WWTP_2** | 100 | 0.59 | 2.61 | 0.38 | *Campylobacterota* | *Campylobacteria* | *Campylobacterales* | *Sulfurospirillaceae* | *Sulfurospirillum* | *Sulfurospirillum sp001548035* |
| **MAG_WWTP_3** | 93.47 | 0.85 | 4.20 | 0.63 | *Pseudomonadota* | *Gammaproteobacteria* | *Burkholderiales* | *Burkholderiaceae* | *Acidovorax* | *Acidovorax temperans* |
| **MAG_WWTP_4** | 91.87 | 3.42 | 2.36 | 0.33 | *Bacteroidota* | *Bacteroidia* | *Flavobacteriales* | *Weeksellaceae* | *Cloacibacterium* | *Cloacibacterium normanense* |

*Table S4: MAGs found in the metagenomic data*

| | | | | | | | | | | |
|---|---|---|---|---|---|---|---|---|---|---|
| **MAG_WWTP_5** | 100 | 0.43 | 4.22 | 0.4 | *Bacteroidota* | *Bacteroidia* | *Bacteroidales* | *Paludibacteraceae* | *Paludibacter* | *Paludibacter sp009929375* |
| **MAG_WWTP_6** | 55.11 | 4.65 | 2.35 | 0.36 | *Bacteroidota* | *Bacteroidia* | *Bacteroidales* | *ML635J-15* | *Bact-19* | |
| **MAG_WWTP_7** | 68.92 | 7.57 | 2.45 | 0.57 | *Spirochaetota* | *Spirochaetia* | *Treponematales* | *Treponemataceae* | *Teretinema* | |
| **MAG_WWTP_8** | 91.22 | 3.58 | 1.14 | 0.34 | *Patescibacteria* | *JAEDAM01* | *Absconditabacterales* | *Absconditicoccaceae* | *UBA5124* | |
| **MAG_WWTP_9** | 99.99 | 0.14 | 5.01 | 0.54 | *Pseudomonadota* | *Alphaproteobacteria* | *Rhizobiales* | *Rhizobiaceae* | *Ochrobactrum_A* | *Ochrobactrum_A quorumnocens* |
| **MAG_WWTP_10** | 100 | 0.60 | 4.72 | 0.64 | *Pseudomonadota* | *Gammaproteobacteria* | *Pseudomonadales* | *Pseudomonadaceae* | *Stutzerimonas* | *Stutzerimonas stutzeri* |
| **MAG_WWTP_11** | 51.71 | 3.23 | 0.89 | 0.5 | *Bacillota_I* | *Bacilli_A* | *Erysipelotrichales* | *Erysipelotrichaceae* | *UBA2212* | |
| **MAG_WWTP_12** | 76.12 | 5.07 | 1.61 | 0.39 | *Bacillota_I* | *Bacilli_A* | *Erysipelotrichales* | *Erysipelotrichaceae* | *UBA2212* | |
| **MAG_WWTP_14** | 96.87 | 2.02 | 3.84 | 0.68 | *Acidobacteriota* | *Holophagae* | *Holophagales* | *Holophagaceae* | *Holophaga* | |
| **MAG_WWTP_15** | 98.69 | 1.08 | 4.72 | 0.36 | *Bacteroidota* | *Bacteroidia* | *Bacteroidales* | *ML635J-15* | *Bact-19* | |
| **MAG_WWTP_16** | 100 | 0.30 | 4.65 | 0.66 | *Pseudomonadota* | *Gammaproteobacteria* | *Xanthomonadales* | *Xanthomonadaceae* | *Stenotrophomonas* | *Stenotrophomonas maltophilia* |
| **MAG_WWTP_17** | 99.99 | 0.50 | 4.00 | 0.63 | *Pseudomonadota* | *Gammaproteobacteria* | *Burkholderiales* | *Rhodocyclaceae* | *Azonexus* | |
| **MAG_WWTP_18** | 99.94 | 2.88 | 3.83 | 0.62 | *Pseudomonadota* | *Gammaproteobacteria* | *Chromatiales* | *Chromatiaceae* | *Thiobaca* | *Thiobaca trueperi* |
| **MAG_WWTP_19** | 74.62 | 3.19 | 3.34 | 0.64 | *Pseudomonadota* | *Gammaproteobacteria* | *Pseudomonadales* | *Pseudomonadaceae* | *Stutzerimonas* | *Stutzerimonas stutzeri* |
| **MAG_WWTP_20** | 98.25 | 3.97 | 4.30 | 0.65 | *Pseudomonadota* | *Gammaproteobacteria* | *Chromatiales* | *Chromatiaceae* | *Thiocapsa* | |
| **MAG_WWTP_21** | 92.53 | 1.45 | 2.09 | 0.34 | *Bacillota_A* | *Clostridia* | *Oscillospirales* | *Ruminococcaceae* | *SZUA-444* | |
| **MAG_WWTP_22** | 57.48 | 0.89 | 1.77 | 0.38 | *Bacteroidota* | *Bacteroidia* | *Bacteroidales* | *Paludibacteraceae* | *Paludibacter* | |
| **MAG_WWTP_23** | 100 | 1.53 | 4.73 | 0.52 | *Pseudomonadota* | *Gammaproteobacteria* | *Enterobacterales* | *Enterobacteriaceae* | *Citrobacter* | *Citrobacter freundii* |
| **MAG_WWTP_24** | 91.15 | 1.78 | 1.83 | 0.34 | *Bacillota_I* | *Bacilli_A* | *Erysipelotrichales* | *Erysipelotrichaceae* | *UBA2212* | |
| **MAG_WWTP_25** | 87.82 | 4.51 | 3.17 | 0.63 | *Pseudomonadota* | *Gammaproteobacteria* | *Burkholderiales* | *Rhodocyclaceae* | *Azonexus* | *Azonexus sp016790885* |
| **MAG_WWTP_26** | 72.4 | 1.53 | 2.07 | 0.47 | *Deferribacterota* | *Deferribacteres* | *Deferribacterales* | *Denitrovibrionaceae* | *Seleniivibrio* | *Seleniivibrio woodruffii_A* |
| **MAG_WWTP_27** | 80.84 | 4.28 | 1.71 | 0.54 | *Pseudomonadota* | *Gammaproteobacteria* | *Burkholderiales* | *Rhodocyclaceae* | *Fluviibacter* | |
| **MAG_WWTP_28** | 71.19 | 4.14 | 2.62 | 0.65 | *Pseudomonadota* | *Gammaproteobacteria* | *Chromatiales* | *Chromatiaceae* | *Thiocapsa* | |
| **MAG_WWTP_29** | 99.91 | 0.46 | 1.96 | 0.45 | *Pseudomonadota* | *Gammaproteobacteria* | *Burkholderiales* | *Burkholderiaceae* | *Polynucleobacter* | |
| **MAG_WWTP_30** | 94.2 | 2.72 | 3.67 | 0.37 | *Bacteroidota* | *Bacteroidia* | *Bacteroidales* | *Paludibacteraceae* | *Paludibacter* | |
| **MAG_WWTP_31** | 71.35 | 3.95 | 1.53 | 0.55 | *Pseudomonadota* | *Gammaproteobacteria* | *Burkholderiales* | *Rhodocyclaceae* | *Fluviibacter* | *Fluviibacter phosphoraccumulans* |
| **MAG_WWTP_32** | 100 | 0.12 | 6.89 | 0.61 | *Pseudomonadota* | *Gammaproteobacteria* | *Pseudomonadales* | *Pseudomonadaceae* | *Pseudomonas_E* | *Pseudomonas_E yamanorum* |
| **MAG_WWTP_33** | 80.23 | 0.20 | 1.06 | 0.43 | *Bacillota_I* | *Bacilli_A* | *RFN20* | | | |
| **MAG_WWTP_34** | 79.38 | 4.86 | 4.98 | 0.48 | *Riflebacteria* | *Ozemobacteria* | *Ozemobacterales* | *Ozemobacteraceae* | *Rifleibacterium* | |

*Table S5: DNA hydrogel junction read information.*

| Junction ID | Type | Read | Direction | Template cigar | Template query start | Template query end | Microbe cigar | Microbe query start | Microbe query end | Microbe contig | Microbe contig length | Contig ref start | Contig ref end |
|---|---|---|---|---|---|---|---|---|---|---|---|---|---|
| **WWTP_day_7_JUNC_074** | CLEAN_ADJACENT | R1 | TemplateA_then_microbe | 20M1D44M1I36M50S | 1 | 101 | 101S50M | 102 | 151 | k141_10721 | 181546 | 31357 | 31406 |
| **WWTP_day_7_JUNC_075** | OVERLAP | R2 | TemplateA_then_microbe | 48S7M1D40M1D56M | 1 | 103 | 58M93S | 94 | 151 | k141_1109 | 63583 | 49749 | 49806 |
| **WWTP_day_7_JUNC_076** | CLEAN_ADJACENT | R1 | TemplateA_then_microbe | 40M111S | 1 | 40 | 40S108M3S | 41 | 148 | k141_11570 | 804 | 109 | 216 |
| **WWTP_day_7_JUNC_078** | CLEAN_ADJACENT | R2 | TemplateA_then_microbe | 35S27M1D89M | 1 | 116 | 35M116S | 117 | 151 | k141_12129 | 186877 | 183611 | 183645 |
| **WWTP_day_7_JUNC_079** | OVERLAP | R2 | TemplateA_then_microbe | 108S43M | 1 | 43 | 117M34S | 35 | 151 | k141_12643 | 27256 | 24781 | 24897 |
| **WWTP_day_7_JUNC_080** | CLEAN_ADJACENT | R1 | TemplateA_then_microbe | 75M76S | 1 | 75 | 75S76M | 76 | 151 | k141_13090 | 64519 | 43555 | 43630 |
| **WWTP_day_7_JUNC_081** | OVERLAP | R2 | TemplateA_then_microbe | 85S44M1D22M | 1 | 66 | 88M1I12M50S | 51 | 151 | k141_13588 | 105105 | 50933 | 51032 |
| **WWTP_day_7_JUNC_082** | NEAR_ADJACENT | R1 | TemplateA_then_microbe | 88M63S | 1 | 88 | 89S62M | 90 | 151 | k141_14755 | 181184 | 29513 | 29574 |
| **WWTP_day_7_JUNC_083** | CLEAN_ADJACENT | R2 | TemplateA_then_microbe | 110S7M1D34M | 1 | 41 | 110M41S | 42 | 151 | k141_14757 | 16596 | 13208 | 13317 |
| **WWTP_day_7_JUNC_084** | OVERLAP | R2 | TemplateA_then_microbe | 71S80M | 1 | 80 | 82M69S | 70 | 151 | k141_15033 | 51239 | 14844 | 14925 |
| **WWTP_day_7_JUNC_085** | OVERLAP | R2 | TemplateA_then_microbe | 56S95M | 1 | 95 | 66M85S | 86 | 151 | k141_15041 | 130655 | 112411 | 112476 |
| **WWTP_day_7_JUNC_086** | OVERLAP | R2 | TemplateA_then_microbe | 50S101M | 1 | 101 | 58M93S | 94 | 151 | k141_15043 | 9738 | 1934 | 1991 |
| **WWTP_day_7_JUNC_087** | OVERLAP | R2 | TemplateA_then_microbe | 111S40M | 1 | 40 | 119M32S | 33 | 151 | k141_15158 | 45243 | 7307 | 7425 |
| **WWTP_day_7_JUNC_088** | OVERLAP | R2 | TemplateA_then_microbe | 63S67M2D21M | 1 | 88 | 69M82S | 83 | 151 | k141_15158 | 45243 | 25945 | 26013 |
| **WWTP_day_7_JUNC_090** | OVERLAP | R2 | TemplateA_then_microbe | 110S41M | 1 | 41 | 111M40S | 41 | 151 | k141_15224 | 83404 | 60329 | 60439 |
| **WWTP_day_7_JUNC_091** | OVERLAP | R2 | TemplateA_then_microbe | 113S38M | 1 | 38 | 116M35S | 36 | 151 | k141_15228 | 25732 | 21713 | 21828 |
| **WWTP_day_7_JUNC_092** | NEAR_ADJACENT | R1 | TemplateA_then_microbe | 93M58S | 1 | 93 | 94S57M | 95 | 151 | k141_15335 | 145498 | 109660 | 109716 |
| **WWTP_day_7_JUNC_093** | OVERLAP | R2 | TemplateA_then_microbe | 46S77M1D28M | 1 | 105 | 47M104S | 105 | 151 | k141_15691 | 72228 | 22036 | 22082 |

| | | | | | | | | | | | | | |
|---|---|---|---|---|---|---|---|---|---|---|---|---|---|
| **WWTP_day_7_JUNC_094** | OVERLAP | R2 | TemplateA_then_microbe | 106S45M | 1 | 45 | 111M40S | 41 | 151 | k141_16287 | 13598 | 6535 | 6645 |
| **WWTP_day_7_JUNC_095** | CLEAN_ADJACENT | R1 | TemplateA_then_microbe | 54M1D39M58S | 1 | 93 | 93S58M | 94 | 151 | k141_16315 | 109894 | 66073 | 66130 |
| **WWTP_day_7_JUNC_096** | OVERLAP | R2 | TemplateA_then_microbe | 78S73M | 1 | 73 | 87M64S | 65 | 151 | k141_16566 | 163379 | 67145 | 67231 |
| **WWTP_day_7_JUNC_097** | CLEAN_ADJACENT | R1 | TemplateA_then_microbe | 37M1D53M61S | 1 | 90 | 90S61M | 91 | 151 | k141_16833 | 27270 | 11476 | 11536 |
| **WWTP_day_7_JUNC_098** | OVERLAP | R2 | TemplateA_then_microbe | 103S48M | 1 | 48 | 105M46S | 47 | 151 | k141_16907 | 150856 | 33207 | 33311 |
| **WWTP_day_7_JUNC_099** | CLEAN_ADJACENT | R2 | TemplateA_then_microbe | 105S46M | 1 | 46 | 1S104M46S | 47 | 150 | k141_17126 | 35220 | 9218 | 9321 |
| **WWTP_day_7_JUNC_100** | OVERLAP | R1 | TemplateA_then_microbe | 22M1D41M88S | 1 | 63 | 62S89M | 63 | 151 | k141_17250 | 41441 | 8308 | 8396 |
| **WWTP_day_7_JUNC_101** | OVERLAP | R2 | TemplateA_then_microbe | 119S32M | 1 | 32 | 122M29S | 30 | 151 | k141_17250 | 41441 | 8953 | 9074 |
| **WWTP_day_7_JUNC_102** | OVERLAP | R2 | TemplateA_then_microbe | 42S106M3S | 4 | 109 | 55M96S | 97 | 151 | k141_17263 | 51352 | 32680 | 32734 |
| **WWTP_day_7_JUNC_103** | OVERLAP | R2 | TemplateA_then_microbe | 115S11M1D25M | 1 | 36 | 120M31S | 32 | 151 | k141_17357 | 92277 | 25018 | 25137 |
| **WWTP_day_7_JUNC_104** | OVERLAP | R2 | TemplateA_then_microbe | 52S35M1D64M | 1 | 99 | 63M88S | 89 | 151 | k141_174 | 35063 | 5713 | 5775 |
| **WWTP_day_7_JUNC_105** | CLEAN_ADJACENT | R2 | TemplateA_then_microbe | 34S27M1D71M1D19M | 1 | 117 | 34M117S | 118 | 151 | k141_18607 | 137237 | 56492 | 56525 |
| **WWTP_day_7_JUNC_106** | OVERLAP | R2 | TemplateA_then_microbe | 54S88M9S | 10 | 97 | 64M87S | 88 | 151 | k141_18768 | 4590 | 2953 | 3016 |
| **WWTP_day_7_JUNC_107** | OVERLAP | R2 | TemplateA_then_microbe | 89S9M1D53M | 1 | 62 | 94M57S | 58 | 151 | k141_19971 | 58328 | 13193 | 13286 |
| **WWTP_day_7_JUNC_108** | CLEAN_ADJACENT | R1 | TemplateA_then_microbe | 107M44S | 1 | 107 | 107S44M | 108 | 151 | k141_20089 | 25378 | 6303 | 6346 |
| **WWTP_day_7_JUNC_109** | OVERLAP | R2 | TemplateA_then_microbe | 96S55M | 1 | 55 | 104M47S | 48 | 151 | k141_20156 | 98553 | 84293 | 84396 |
| **WWTP_day_7_JUNC_110** | OVERLAP | R2 | TemplateA_then_microbe | 57S94M | 1 | 94 | 66M85S | 86 | 151 | k141_20664 | 190439 | 79298 | 79363 |
| **WWTP_day_7_JUNC_111** | OVERLAP | R2 | TemplateA_then_microbe | 31S120M | 1 | 120 | 36M115S | 116 | 151 | k141_2076 | 219623 | 14884 | 14919 |
| **WWTP_day_7_JUNC_112** | NEAR_ADJACENT | R2 | TemplateA_then_microbe | 72S58M1I20M | 1 | 79 | 71M80S | 81 | 151 | k141_20972 | 28562 | 27314 | 27384 |
| **WWTP_day_7_JUNC_113** | OVERLAP | R2 | TemplateA_then_microbe | 98S53M | 1 | 53 | 99M52S | 53 | 151 | k141_20991 | 44689 | 13010 | 13108 |

| | | | | | | | | | | | | | |
|---|---|---|---|---|---|---|---|---|---|---|---|---|---|
| **WWTP_day_7_JUNC_114** | OVERLAP | R2 | TemplateA_then_microbe | 50S47M1D54M | 1 | 101 | 51M100S | 101 | 151 | k141_20991 | 44689 | 16043 | 16093 |
| **WWTP_day_7_JUNC_115** | CLEAN_ADJACENT | R1, R2 | both_directions | 37M114S | 1 | 37 | 37S114M | 38 | 151 | k141_212 | 18853 | 16212 | 16325 |
| **WWTP_day_7_JUNC_117** | OVERLAP | R2 | TemplateA_then_microbe | 104S8M1D34M1D5M | 1 | 47 | 106M45S | 46 | 151 | k141_21861 | 7095 | 5495 | 5600 |
| **WWTP_day_7_JUNC_118** | OVERLAP | R2 | TemplateA_then_microbe | 107S27M1D17M | 1 | 44 | 113M38S | 39 | 151 | k141_21950 | 61987 | 46691 | 46803 |
| **WWTP_day_7_JUNC_120** | CLEAN_ADJACENT | R2 | TemplateA_then_microbe | 72S52M1D27M | 1 | 79 | 72M79S | 80 | 151 | k141_24109 | 36338 | 6300 | 6371 |
| **WWTP_day_7_JUNC_121** | OVERLAP | R2 | TemplateA_then_microbe | 47S104M | 1 | 104 | 49M102S | 103 | 151 | k141_24644 | 42850 | 42435 | 42483 |
| **WWTP_day_7_JUNC_122** | OVERLAP | R2 | TemplateA_then_microbe | 33S16M1D22M1D17M1D30M1D22M1D11M | 1 | 118 | 39M112S | 113 | 151 | k141_3333 | 14046 | 4760 | 4798 |
| **WWTP_day_7_JUNC_123** | OVERLAP | R2 | TemplateA_then_microbe | 91S60M | 1 | 60 | 98M53S | 54 | 151 | k141_5089 | 96924 | 41516 | 41613 |
| **WWTP_day_7_JUNC_124** | OVERLAP | R2 | TemplateA_then_microbe | 75S34M42S | 43 | 76 | 76M75S | 76 | 151 | k141_526 | 215862 | 152548 | 152623 |
| **WWTP_day_7_JUNC_125** | OVERLAP | R2 | TemplateA_then_microbe | 47S104M | 1 | 104 | 56M95S | 96 | 151 | k141_5265 | 2977 | 655 | 710 |
| **WWTP_day_7_JUNC_126** | OVERLAP | R1 | TemplateA_then_microbe | 45M106S | 1 | 45 | 43S108M | 44 | 151 | k141_5352 | 8116 | 4299 | 4406 |
| **WWTP_day_7_JUNC_127** | OVERLAP | R2 | TemplateA_then_microbe | 87S64M | 1 | 64 | 95M56S | 57 | 151 | k141_673 | 113569 | 99832 | 99926 |
| **WWTP_day_7_JUNC_128** | OVERLAP | R2 | TemplateA_then_microbe | 87S18M1D46M | 1 | 64 | 93M58S | 59 | 151 | k141_7017 | 320152 | 44699 | 44791 |
| **WWTP_day_7_JUNC_129** | CLEAN_ADJACENT | R1 | TemplateA_then_microbe | 64M87S | 1 | 64 | 64S87M | 65 | 151 | k141_7524 | 29434 | 27409 | 27495 |
| **WWTP_day_7_JUNC_130** | OVERLAP | R2 | TemplateA_then_microbe | 65S10M1D31M1I40M1D4M | 1 | 86 | 74M77S | 78 | 151 | k141_7524 | 29434 | 5194 | 5267 |
| **WWTP_day_7_JUNC_131** | CLEAN_ADJACENT | R1 | TemplateA_then_microbe | 59M1D23M69S | 1 | 82 | 82S69M | 83 | 151 | k141_7592 | 16357 | 1417 | 1485 |
| **WWTP_day_7_JUNC_132** | CLEAN_ADJACENT | R2 | TemplateA_then_microbe | 74S77M | 1 | 77 | 74M77S | 78 | 151 | k141_7722 | 52885 | 31579 | 31652 |
| **WWTP_day_7_JUNC_133** | OVERLAP | R2 | TemplateA_then_microbe | 101S50M | 1 | 50 | 110M41S | 42 | 151 | k141_7766 | 109021 | 370 | 479 |
| **WWTP_day_7_JUNC_134** | OVERLAP | R2 | TemplateA_then_microbe | 82S69M | 1 | 69 | 90M61S | 62 | 151 | k141_8402 | 36992 | 555 | 644 |

| | | | | | | | | | | | | | |
|---|---|---|---|---|---|---|---|---|---|---|---|---|---|
| **WWTP_day_7_JUNC_135** | OVERLAP | R2 | TemplateA_then_microbe | 50S101M | 1 | 101 | 59M92S | 93 | 151 | k141_8402 | 36992 | 13425 | 13483 |
| **WWTP_day_7_JUNC_136** | OVERLAP | R2 | TemplateA_then_microbe | 56S36M1D29M1I29M | 1 | 95 | 75M76S | 77 | 151 | k141_8456 | 442 | 368 | 442 |
| **WWTP_day_7_JUNC_137** | OVERLAP | R2 | TemplateA_then_microbe | 34S76M1D41M | 1 | 117 | 40M111S | 112 | 151 | k141_8552 | 34511 | 20295 | 20334 |
| **WWTP_day_7_JUNC_138** | OVERLAP | R2 | TemplateA_then_microbe | 116S35M | 1 | 35 | 126M25S | 26 | 151 | k141_8552 | 34511 | 25421 | 25546 |
| **WWTP_day_7_JUNC_139** | OVERLAP | R2 | TemplateA_then_microbe | 70S37M1D44M | 1 | 81 | 71M80S | 81 | 151 | k141_8633 | 33389 | 17912 | 17982 |
| **WWTP_day_7_JUNC_140** | OVERLAP | R1 | TemplateA_then_microbe | 101M50S | 1 | 101 | 98S53M | 99 | 151 | k141_8761 | 72578 | 60490 | 60542 |
| **WWTP_day_7_JUNC_141** | OVERLAP | R2 | TemplateA_then_microbe | 72S79M | 1 | 79 | 75M76S | 77 | 151 | k141_9630 | 54154 | 4911 | 4985 |
| **Mesocosm_day_7_JUNC_031** | OVERLAP | R2, R1 | both_directions | 66S42M | 67 | 108 | 75M33S | 1 | 75 | k141_15847 | 3736 | 1690 | 1764 |
| **Mesocosm_day_7_JUNC_033** | OVERLAP | R2, R1 | both_directions | 78S31M | 79 | 109 | 83M26S | 1 | 83 | k141_16524 | 2645 | 2417 | 2499 |
| **Mesocosm_day_7_JUNC_035** | OVERLAP | R2 | TemplateA_then_microbe | 77S74M | 1 | 74 | 78M73S | 74 | 151 | k141_18122 | 2995 | 342 | 419 |
| **Mesocosm_day_7_JUNC_036** | OVERLAP | R2 | TemplateA_then_microbe | 75S76M | 1 | 76 | 2S75M74S | 75 | 149 | k141_18754 | 2185 | 962 | 1036 |
| **Mesocosm_day_7_JUNC_037** | OVERLAP | R2 | TemplateA_then_microbe | 113S38M | 1 | 38 | 121M30S | 31 | 151 | k141_22519 | 3183 | 473 | 593 |
| **Mesocosm_day_7_JUNC_038** | CLEAN_ADJACENT | R2, R1 | both_directions | 91S41M | 92 | 132 | 91M41S | 1 | 91 | k141_22911 | 31718 | 266 | 356 |
| **Mesocosm_day_7_JUNC_039** | OVERLAP | R2, R1 | both_directions | 104S47M | 105 | 151 | 110M41S | 1 | 110 | k141_22911 | 31718 | 30634 | 30743 |
| **Mesocosm_day_7_JUNC_042** | OVERLAP | R2, R1 | both_directions | 117S34M | 118 | 151 | 119M32S | 1 | 119 | k141_26614 | 2365 | 390 | 508 |
| **Mesocosm_day_7_JUNC_044** | CLEAN_ADJACENT | R2, R1 | both_directions | 86S32M | 87 | 118 | 86M32S | 1 | 86 | k141_26742 | 365 | 9 | 94 |
| **Mesocosm_day_7_JUNC_046** | OVERLAP | R2, R1 | both_directions | 74S39M | 75 | 113 | 87M26S | 1 | 87 | k141_28505 | 9190 | 6819 | 6905 |
| **Mesocosm_day_7_JUNC_048** | OVERLAP | R2, R1 | both_directions | 115S36M | 116 | 151 | 118M33S | 1 | 118 | k141_29079 | 456 | 126 | 243 |
| **Mesocosm_day_7_JUNC_050** | OVERLAP | R2, R1 | both_directions | 85S59M | 86 | 144 | 90M54S | 1 | 90 | k141_30492 | 121659 | 54690 | 54779 |
| **Mesocosm_day_7_JUNC_052** | OVERLAP | R2, R1 | both_directions | 106S40M | 107 | 146 | 113M33S | 1 | 113 | k141_32541 | 4288 | 2382 | 2494 |

| | | | | | | | | | | | | | |
|---|---|---|---|---|---|---|---|---|---|---|---|---|---|
| **Mesocosm_day_7_JUNC_054** | OVERLAP | R2, R1 | both_directions | 91S40M | 92 | 131 | 96M35S | 1 | 96 | k141_3765 | 94542 | 22360 | 22455 |
| **Mesocosm_day_7_JUNC_055** | OVERLAP | R2 | TemplateA_then_microbe | 103S48M | 1 | 48 | 116M35S | 36 | 151 | k141_3765 | 94542 | 31041 | 31156 |
| **Mesocosm_day_7_JUNC_057** | OVERLAP | R2, R1 | both_directions | 82S36M | 83 | 118 | 86M32S | 1 | 86 | k141_38352 | 10916 | 1513 | 1598 |
| **Mesocosm_day_7_JUNC_059** | OVERLAP | R2, R1 | both_directions | 68S45M | 69 | 113 | 78M35S | 1 | 78 | k141_38857 | 62475 | 34059 | 34136 |
| **Mesocosm_day_7_JUNC_060** | OVERLAP | R2, R1 | both_directions | 70S36M | 71 | 106 | 76M30S | 1 | 76 | k141_38857 | 62475 | 46960 | 47035 |
| **Mesocosm_day_7_JUNC_061** | OVERLAP | R2 | TemplateA_then_microbe | 113S38M | 1 | 38 | 124M27S | 28 | 151 | k141_38857 | 62475 | 50341 | 50464 |
| **Mesocosm_day_7_JUNC_062** | CLEAN_ADJACENT | R1 | microbe_then_TemplateA | 38S27M | 39 | 65 | 38M27S | 1 | 38 | k141_38857 | 62475 | 6200 | 6237 |
| **Mesocosm_day_7_JUNC_065** | NEAR_ADJACENT | R2 | TemplateA_then_microbe | 99S52M | 1 | 52 | 98M53S | 54 | 151 | k141_41866 | 379 | 61 | 158 |
| **Mesocosm_day_7_JUNC_066** | CLEAN_ADJACENT | R1, R2 | both_directions | 32M67S | 1 | 32 | 32S67M | 33 | 99 | k141_42844 | 1124 | 579 | 645 |
| **Mesocosm_day_7_JUNC_068** | OVERLAP | R2, R1 | both_directions | 96S50M | 97 | 146 | 105M41S | 1 | 105 | k141_44666 | 8781 | 6838 | 6942 |
| **Mesocosm_day_7_JUNC_070** | OVERLAP | R2 | TemplateA_then_microbe | 107S44M | 1 | 44 | 119M32S | 33 | 151 | k141_50337 | 8375 | 7497 | 7615 |
| **Mesocosm_day_7_JUNC_071** | OVERLAP | R2 | TemplateA_then_microbe | 109S42M | 1 | 42 | 116M35S | 36 | 151 | k141_6051 | 8742 | 4801 | 4916 |
| **Mesocosm_day_7_JUNC_072** | OVERLAP | R2, R1 | both_directions | 120S31M | 121 | 151 | 123M28S | 1 | 123 | k141_6169 | 2087 | 895 | 1017 |
| **Mesocosm_day_100_JUNC_001** | NEAR_ADJACENT | R2 | TemplateA_then_microbe | 49M102S | 1 | 49 | 30S71M50S | 51 | 121 | k141_125614 | 303 | 1 | 71 |
| **Mesocosm_day_100_JUNC_002** | OVERLAP | R2 | TemplateA_then_microbe | 110S41M | 1 | 41 | 116M1D5M30S | 31 | 151 | k141_127981 | 64356 | 32493 | 32614 |
| **Mesocosm_day_100_JUNC_003** | OVERLAP | R2 | TemplateA_then_microbe | 96S55M | 1 | 55 | 105M46S | 47 | 151 | k141_145659 | 47586 | 32164 | 32268 |
| **Mesocosm_day_100_JUNC_004** | OVERLAP | R2 | TemplateA_then_microbe | 115S36M | 1 | 36 | 119M32S | 33 | 151 | k141_148640 | 416 | 27 | 145 |
| **Mesocosm_day_100_JUNC_005** | OVERLAP | R2 | TemplateA_then_microbe | 116S5M1D30M | 1 | 35 | 118M33S | 34 | 151 | k141_150686 | 62808 | 13667 | 13784 |
| **Mesocosm_day_100_JUNC_007** | OVERLAP | R2 | TemplateA_then_microbe | 113S38M | 1 | 38 | 118M33S | 34 | 151 | k141_17236 | 1305 | 814 | 931 |
| **Mesocosm_day_100_JUNC_008** | OVERLAP | R2 | TemplateA_then_microbe | 111S30M1D3M1D7M | 1 | 40 | 115M36S | 37 | 151 | k141_184241 | 249594 | 44874 | 44988 |

| | | | | | | | | | | | | | |
|---|---|---|---|---|---|---|---|---|---|---|---|---|---|
| **Mesocosm_day_100_JUNC_009** | OVERLAP | R2, R1 | both_directions | 100S51M | 101 | 151 | 108M43S | 1 | 108 | k141_184412 | 1299 | 198 | 305 |
| **Mesocosm_day_100_JUNC_011** | CLEAN_ADJACENT | R2, R1 | both_directions | 120S31M | 121 | 151 | 120M31S | 1 | 120 | k141_184760 | 379 | 2 | 121 |
| **Mesocosm_day_100_JUNC_013** | NEAR_ADJACENT | R1 | TemplateA_then_microbe | 36M115S | 1 | 36 | 37S114M | 38 | 151 | k141_19217 | 11359 | 10142 | 10255 |
| **Mesocosm_day_100_JUNC_014** | OVERLAP | R2, R1 | both_directions | 113S37M | 114 | 150 | 117M33S | 1 | 117 | k141_206680 | 231428 | 48285 | 48401 |
| **Mesocosm_day_100_JUNC_017** | OVERLAP | R1 | TemplateA_then_microbe | 82M69S | 1 | 82 | 75S76M | 76 | 151 | k141_211671 | 124427 | 112655 | 112730 |
| **Mesocosm_day_100_JUNC_019** | OVERLAP | R2 | TemplateA_then_microbe | 110S41M | 1 | 41 | 115M36S | 37 | 151 | k141_236618 | 2893 | 1706 | 1820 |
| **Mesocosm_day_100_JUNC_020** | NEAR_ADJACENT | R1 | TemplateA_then_microbe | 87M64S | 1 | 87 | 88S57M6S | 89 | 145 | k141_51754 | 12254 | 2302 | 2358 |
| **Mesocosm_day_100_JUNC_021** | OVERLAP | R2 | TemplateA_then_microbe | 87S50M1D14M | 1 | 64 | 91M60S | 61 | 151 | k141_71565 | 82488 | 5477 | 5567 |
| **Mesocosm_day_100_JUNC_022** | OVERLAP | R2, R1 | both_directions | 95S34M5D20M | 96 | 149 | 104M45S | 1 | 104 | k141_71565 | 82488 | 81161 | 81264 |
| **Mesocosm_day_100_JUNC_024** | NEAR_ADJACENT | R2, R1 | both_directions | 84S35M | 85 | 119 | 83M36S | 1 | 83 | k141_9751 | 5295 | 274 | 356 |

*Table S6: DNA hydrogel junction sequences*

a DNA hydrogel a gDNA a Overlap a Unassigned

| Junction | Sequence |
| --- | --- |
| WWTP_day_7_JUNC_074 | ACGTCGGACGCGATTCATTGCCAGTCAGCCGCCACCGTAGGTGGACACGCCGCATCCTACATCCAAGCGTCAACGTCGGACGCGATTCATTGACCAGTCAGACAACCGTTCCGATGGTAACACCTGCCATCCGCATGGAGGCGTAAAATGC |
| WWTP_day_7_JUNC_075 | GTCCGACGTTGACGCTGGATGTAGGATGCGGCGTGTCCACCTCCGGTGGCGGCTGATGGTCAATGAATCGCGTCCGACGTTGAAGCTGGATGTAGGTGCTGCGCAAGCTGCTGGCCAAAGACCCGGCGGAACGCTTCCCGGATGCGGCGGC |
| WWTP_day_7_JUNC_076 | ATCCTACATCCAGCGTCAACGTCGGACGCGATTCATTGACAGAGAAGATAGATATCCAATAGCAGATATTAGTCAAACACAAGCTCTAACAGCTTGTAATTGAATCTGAAGTTGATTTCACTTAATTACGAATAATGAATGGATGACAATT |
| WWTP_day_7_JUNC_078 | CCTACGGTGGCGGCTGACTGGTCAATGAATCGCGTCCGACGTTGACGCTGGATGTAGGATGCGGCGTGTCCACCTACGGTGGCGGCTGATGGTCAATGAATCGCGTCCGACGTTGAAGTCTCTACAAGACGGCGTCCGTCCAGATACCGGA |
| WWTP_day_7_JUNC_079 | AATGAATCGCGTCCGACGTTGACGCTGGATGTAGGATACGGCGAGATGTCGAGCTCCAACATTCCCCAGCCCATGCCGGCGTAATCCTCATTATCGTGATGGGCGGCCACCGACGTTGGCATCGGAATCACCCGGCCTTCTGCGGCGATCT |
| WWTP_day_7_JUNC_080 | CATCCAGCGTCAACGTCGGACGCGATTCATTGACCAGTCAGCCGCCACCGTAGGTGGACACGCCGCATCCTACATAGGGCTTTTCTTTCATGGCGTCTCCTTCACATCAGTGAGGGGATCAGTCGGCGGTAGTCATTCAGTGACGGATGGC |
| WWTP_day_7_JUNC_081 | GTCCACCTACGGTGGCGGCTGATGGTCAATGAATCGCGTCCGACGTTGACGCTGGATGTAGGATGCCCGGCGGGAGTTCTGGGCTTTGTTGGGCGGGCTCAGATCGCATAGGAACCGTCCTCACGCTGAATCAGGCCATCGGTGTAGTTGC |
| WWTP_day_7_JUNC_082 | CGTCGGACGCGATTCATTGACCAGTCAGCCGCCACCGTAGGTGGACACGCCGCATCCTACATCCAGCGTCAACGTCGGACGCGATTCAAGGAGATTGACGCTCTTTTCCTTGCGGATTGTAATGTCTATGCAGCATTGCAAGCCCCATCTT |
| WWTP_day_7_JUNC_083 | TGAATCGCGTCCGACGTTGACGCTGGATGTAGGAGCGGCGTCCCACGCCGCCACGGGCATAGCGTTCGGCAGCGTAGTGGTGAGTGGCCGAAGTGACGGCGGCGTGGCCGAGCCAGACCGTCGAGTCGTGCCCGCCCGATTGCGTCGGCCC |
| WWTP_day_7_JUNC_084 | GATGTAGGATGCGGCGTGTCCACCTCCGGTGGCGGCTGACTGGTCAATGAATCGCGTCCGACGTTGACGCTGGGTGTAGGGGTGGTAGTGACCGGGGTTGAAAGTGTCGCCGATTATCACGCGGTGAGTGAGTTGGGAGCTGCGTGGTTGC |
| WWTP_day_7_JUNC_085 | GGCGTGTCCACCTACGGTGGCGGCTGACTGGTCAATGAATCGCGTCCGACGTTGACGCTGGATGTAGGATGCGGCGTGTCCCCCTACGGCGGCGGTGATGAACATGATGCTGTTGCCGGCCGAAGCGGCGGTGCCGGCGCTGCTGGAGAAC |
| WWTP_day_7_JUNC_086 | GGCGTGTCCACCTACGGTGGCGGCTGACTGGTCAATGAATCGCGTCCGACGTTGACGCTGGATGTAGGATGCGGCGTGTCCACCTACGGTGGCGGTTTACTTGAAGTCGCGGCTGACCCTGGCGCTGATTTTCCAGATGCGCCCGTTGTCC |
| WWTP_day_7_JUNC_087 | TGGCGGCTGACTGGTCAATGAATCGCGTCCGACGTTGACGAGGATGGCCACGCCGCATTTCTGCAGTTTCTGCATGTTCTCGGGTTTGAAGATCAGGTCCCATGAATCTACCGGGGCGTTGTCGCCGAGTACCGCCTTGACCTTGTCGATG |
| WWTP_day_7_JUNC_088 | CTGGATGTAGGATGCGGCGTGCACCTACGGTGGCGGCTGACTGGTCAATGAATCGCGTCCGACGTTGACGCTGGATGTAGGATGCGGCTCGGCATTGCGGTTGAGGTCCTGCACCCAGGTATTGGCCCGGGCCGGGTCCGACAGGTTGGCA |
| WWTP_day_7_JUNC_090 | GGTGGCGGCTGACTGGTCAATGAATCGCGTCCGACGTTGACACAACGACAGGGCAGGCGGCGATGACGCCGGCCTGGTGGCGATCGCTGCGCGGCTGGCTGGCGCGCTCGACCGGGGCGGCTCCGTTGCGCCTGGCGGCCGTATCGCGACA |
| WWTP_day_7_JUNC_091 | GTTGACGCTGGATGTAGGATGCGGCGTGTCCCCCTACGCCATGCCACCCGGTACATTCAATTGTTCGGCGCTGAAGTGCGCGCCCTCAACCGTGCTCGACAGCCCCGGCATCTGCTCCATCAGCCCCGCGACCATACGCGCCGACGCCACC |
| WWTP_day_7_JUNC_092 | ACGCCGCATCCTACATCCAGCGTCAACGTCGGACGCGATTCATTGACCAGTCAGCCGCCACCGTAGGTGGACACGCCGCATCCTACATCCAGCTATGAAAAAACCGATGCCATAGGTGGCGAACGCCAGCAGCAGTTGGTTTGAGCCTTCC |
| WWTP_day_7_JUNC_093 | CGACGTTGACGCTGGATGTAGGATGCGGGTGTCCACCTACGGTGGCGGCTGACTGGTCAATGAATCGCGTCCGACGTTGACGCTGGATGTAGGATGCGGCGTGTCTCTGGCTGTATTCACGCTGTCTGAGCAGTTCACACCATCAGAACGC |
| WWTP_day_7_JUNC_094 | TCAATGAATCGCGTCCGACGTTGACGCTGGATGTAGGATGCGGCGACAGCCACACCACCACCTACGGTGCCTTGGGCGCATTCGGCTTTGGCATCGGCACCTCGGAAATCGAGCATTTGCTGGCCTCGCAAACCCTGGTCTACAAACGCCT |
| WWTP_day_7_JUNC_095 | CAGTCAGCCGCCACCGTAGGTGGACACGCCGCATCCTACATCCAGCGTCAACGTGGACGCGATTCATTGACCAGTCAGCCGCCACCGTAGGTGACTGGATCGCGGGCAACCAATCTGATCAACACCTTCGTCACTCTGACTAAAAACAATT |
| WWTP_day_7_JUNC_096 | GTAGGATGCGGCGTGTCCACCTACGGTGGCGGCCGAATGGTCAATGAATCGCGTCCGACGTTGACGCTGGATGAGTTGCGTATTTCGCGCGTGGCACGACCGGCCGCGCTGCTGCTGGCCGATGCCACCGCGCAGGGCGCCGACTCGCGCC |
| WWTP_day_7_JUNC_097 | GGTGGACACGCCGCATCCTACATCCAGCGTCAACGTCGACGCGATTCATTGACCAGTCAGCCGCCACCGTAGGTGGACACGCCGCATCCTGTTCAGTGATATTCCGTGGAGTGTGATTATGGGGGCAGGCAATGACGACAGAATGCCTGTT |
| WWTP_day_7_JUNC_098 | CGCTGGATGTAGGATGCGGCGTGTCCCCCTCCGGTGGCGGCTGACTGGACGTCTATCAGGGCTTCGCATGGGGCACGGGTATCGACCGTATCGCCATGCTGAAATATGGCATGCCTGATCTGCGCGCCTTCTTCGATGCCGATGTGCGCTG |
| WWTP_day_7_JUNC_099 | CGCTGGATGTAGGATGCGGCGTGTCCCCCTACGGTGGCGGCTGACTCGGTAGGCGGAACGCGGTAGGACAAGGCACCGCCGATCATGAACACGAAGTAGATCGCGGCCATGACCAAGAAGCTCTGCCACACGCCAACGCTGGTCGGCGAAA |
| WWTP_day_7_JUNC_100 | CCAGTCAGCCGCCACCGTAGGTGACACGCCGCCTCCTACATCCAGCGTCAACGTCGGACGCGACCGTCCATGCTGGTTGGAGTTGCACCGTCCGCAGCGCTTGGCACCTGTCCCGGCAACTCCTGAATGTCTGGCATCAGGCGCTCGTTGT |
| WWTP_day_7_JUNC_101 | GTCAATGAATCGCGTCCGACGTTGACGCTGGACGTGGAAGCTGGGCAGACTCAACGAGCAGCGCGAGAACATGAAGACACTTGGCGCGCACGGCGCTGGTGCTGCAACGTTCTTCGCCTTGGGCGCAGGAATCATGGACCCGGCTGGCTGG |
| WWTP_day_7_JUNC_102 | CGGCGTTGACGCTGGATGTAGGATGCGGCGTGTCCACCTACGGTGTCGGCTGACTGGTCAATGAATCGCGTCCGGCGTTGACGCTGGATGTAGGATGCGGAGTGTCCACGGCGTTGCTCAACGTGTTGAATGCCACGCTGATACCCCTGAT |
| WWTP_day_7_JUNC_103 | CGCTGGATGTAGGATGCGGCGTGTCACCTACGGTGGTACGCTGGTCCGCTTTGCCGGACCTCGCTTGGCAGGTGCCAGCGAAAATGATCCGCTACTGCCTGTCCGTCTTCGCAAGGGAGAACGCGCACTCGGTGGTACCTTGTCATGGTCA |
| WWTP_day_7_JUNC_104 | TGAATCGCGTCCGACGTTGACGCTGGATGTAGGATGCGGCGTGTCCACCTACGGTGGCGGCTGATGGTCAATGAATCGCGTCCGACGTTGACGCTGGATCAGTTGGGTGAACAGCAGAAGATTGCGGATGCGTTCTTCAAGGCGGGGTTGT |
| WWTP_day_7_JUNC_105 | GCGGCTGACTGGTCAATGATCGCGTCCGACGTTGACGCTGGATGTAGGATGCGGCGTGTCCACCTACGGTGGCGGCTGACTGGTCAATGATCGCGTCCGACGTTGACGCTGGATGTACTTGGATGTCCAATATCTCGGTACTAGACACCGT |
| WWTP_day_7_JUNC_106 | ATCGCGTGAACGTTGACGCTGGATGTAGGATGCGGCGTGTCCACCTACGGTGGCGGCTGACTGGTCAATGAATCGCGTCCGACGTTGGCGCAGGATGCGTTTGCCGGCGGTGTAGTCGTTGATCCCATCGCACGGCGTGTAGTTGCGTTCC |
| WWTP_day_7_JUNC_107 | CCGACGTTGACGCTGGATGTAGGATGCGGCGTGTCCACCTACGGTGGCGGCTGCTGGTCAATCAGGTACATGTTGCTGACAACATCAGGTGTTGTACCCGGCTCGCGCGTAAAGGCCAGGGGCCGGATAACGATCTCCTTGCCGTCGACGG |
| WWTP_day_7_JUNC_108 | TCAACGTCGGACGCGATTCATTGACCAGTCAGCCGCCACCGTAGGTGGACACGCCGCATCCTACATCCAGCGTCAACGTCGGACGCGATTCATTGACCAGTCAGCCGACATCGGTGGCGCGGGCGTGGCCAAGGGTTATCTGAACCAGCCG |
| WWTP_day_7_JUNC_109 | ACCTACGGTGGCGGCTGACTGGTCAATGAATCGCGTCCGACGTTGACGCTGGATGAAGAAGCGTTCGTGGCATTTTTGGCCAAGCACAACATCACGGTCTGATGATCGTTCCCACGCAGAGCGTGGGAACGATCTCTCCCGGGCAATGCTC |
| WWTP_day_7_JUNC_110 | GGCGTGTCCACCTACGGTGGCGGCTGACTGGTCAATGAATCGCGTCCGACGTTGACGCTGGATGTAGGATGCGGCGTGTCAACCTACGGTGGCGAAGATGTTCAACTGGCTGTTTACAATGTATCGCGGACGCATTCGCTTTGAAGTGCCG |
| WWTP_day_7_JUNC_111 | TCAATGAATCGCGTCCGACGTTGACGCTGGATGTAGGATGCGGCGTGTCCCCCTACGGTGGCGGCTGACTGGTCAATGAATCGCGTCCGACGTTGACGCTGGATGTAGGATGCGGCGTGTTCCCCCGCGTTGGCCAGCGTGGTTTTGCCGC |
| WWTP_day_7_JUNC_112 | GTTGACGCTGGATGTAGGATTGCGGCGTGTCCACCTACGGTGGCGGCTGACTGGTCAACGAATCGCGTCCGAAATTGACATGGCGATGCCATTGCGTGGCCAGCTTGAACACCAGGTGGGCGGTCGCGTCGGCAGCGGTGTACGGGTCGGT |
| WWTP_day_7_JUNC_113 | GTTGACGCTGGATGTAGGATGCGGCGTGTCCCCCTACGGTGGCGGCTGACTGGCTTGATTGTAAGCTTCAACACTTGCAAACATTCTCTCTCGCTCCCTGCGATGTAGTTCCATTAAGTTTAGCCAGCCATCACTGCCAACTTCTAGCTTG |
| WWTP_day_7_JUNC_114 | CCGACGTTGACGCTGGATGTAGGATGCGGCGTGTCCACCTCCGGTGGCGGCTGATGGTCAATGAATCGCGTCCGACGTTGACGCTGGATGTAGGATGCGGCTCGAAAGTCCAGAGTGTACACAAAGTAGATTGCATCCAATCCAACGTAGC |
| WWTP_day_7_JUNC_115 | GGACGCGATTCATTGACCAGTCAGCCGCCACCGTAGGACCAGGCCCAGGGACAAGGTGCTTGGGGCCAGGATGCCCAGCAGCGTATCGGAAGCCAGCGGCTGGCGCTGATGCAGGGTCACCAGCAGCACCGCCAGCAACAGGCAGCCGACG |
| WWTP_day_7_JUNC_117 | CTGGTAATGAATCGCGTCCGACGTTGACGCTGGATGTAGATGCGGCGCCGAACTGATGGCCTCGTGCGCACGCTACCTGCGCGCCAACAAGCGCCGCGAATCGGTCACCTTCGAATACACCCTGATGAAGGGCATCAACGACAAGCCCGAG |
| WWTP_day_7_JUNC_118 | GATGTAGGATGCGGCGTTCCACCTACGGTGGCGGCTGACTGGTCTTCTGTTTCGTCAGTCATCCGAGGCTTCTCACTGCTGTTTTTGGTTTATTTTATTTCTGGATGTTTCATTGAAATGGCGTGAAATATGGCAAAGCTTCAACGCTTCA |
| WWTP_day_7_JUNC_120 | GTCCACCTACGGTGGCGGCTGACTGGTAATGAATCGCGTCCGACGTTGACGCTGGATGTAGGATGCGGCGTGTCCACCTCGCGCGGTTTTTTCTCTCAAGCCCTGCCGTGGCGCTGGTGTACACCGAGGCATGCATCAGCAAGGCCGAGGC |
| WWTP_day_7_JUNC_121 | GCGGCTGACTGGTCAATGAATCGCGTCCGACGTTGACGCTGGATGTAGGATGCGGCGTGTCCACCTACGGTGGCGGCTGACTGGTCAATGAATCGCGTCCGACGCTTCCTTGGCGGTGAACGGTCGCAAGCCCGCATCAGCCCCCCACAAT |
| WWTP_day_7_JUNC_122 | TACGGTGGCGGTGACTGGTCAATGAATCGCGTCGACGTTGACGCTGGATGTAGGATGCGGCGTTCCACCTACGGTGGCGGTGACTGGTCAATGAATCGCGTCGACGTTGACGCTGGATCAAGGAGCGCTCGGACCGCAGGAAGCCTTCCTT |
| WWTP_day_7_JUNC_123 | GAATCGCGTCCGACGTTGACGCTGGATGTAGGATGCGGCGTGTCCACCTACGGTGGCGGCCGCGCTGGATGTGCCGACGATTTCCCTGTTCGGCCCCACCAATCCAGGCCTGACCGGCGCCTACGGCAAGGCGCAGGTCCACCTGGCCAGC |
| WWTP_day_7_JUNC_124 | GTGGGAAGCGGGAACAGACCTGGGGTACGGCACCACCACGTTTGACTGGTCAATGAATCGCGTCCGACGTTGACGCCGGGTAACCGATACGGTCGCCTTCGGTGAGGGCTTCAGCAATGTCGGCAACGTTACCTTCGGTGCCAGGGGTGAC |
| WWTP_day_7_JUNC_125 | TCCGACGTTGACGCTGGATGTAGGATGCGGCGTGTCCACCTACGGTGGCGGCTGACTGGTCAATGAATCGCGTCCGACGTTGACGCTGGATGTAGGATGCGGCGGTGGAAGCGGCGGTGGCGCAGGGTGGTGCGAGCTTTGCGGCGATCCA |
| WWTP_day_7_JUNC_126 | CAACGTCGGACGCGATTCATTGACCAGTCAGCCGCCACCGTAGGTCGCGCTGGCGCAATTGATCATCGTCATGCTACCCAGCAACCGCGCCACTACCGACTGGTCCGGCTGCGCGCCAATCCGCAATACGCAGTCGAAACCTTCGGCCAGC |
| WWTP_day_7_JUNC_127 | GTCCGACGTTGACGCTGGATGTAGGATGCGGCGTGTCCACCTACGGTGGCGGCTGACTGGTCAAGAATGGTGCGGGCCTGCTGACCCTCAACAACACCAACACCTTCGGTGGTGGCCTGACCTTGAACGGTGGTGGCCTGGTGGTCGGCGC |
| WWTP_day_7_JUNC_128 | TGGATGTAGGATGCGGCGTGTCCACCTACGGTGGCGGCTGACTGGTAATGAATCGCGTCCGACGCCGCCGTGGCCTACAAAACCCATCGGCTTCTGGATCCCGTCACCGTAGTCAATCGCGTTCTTTCAAACGGCCGTGGGAGCGTGGTTG |

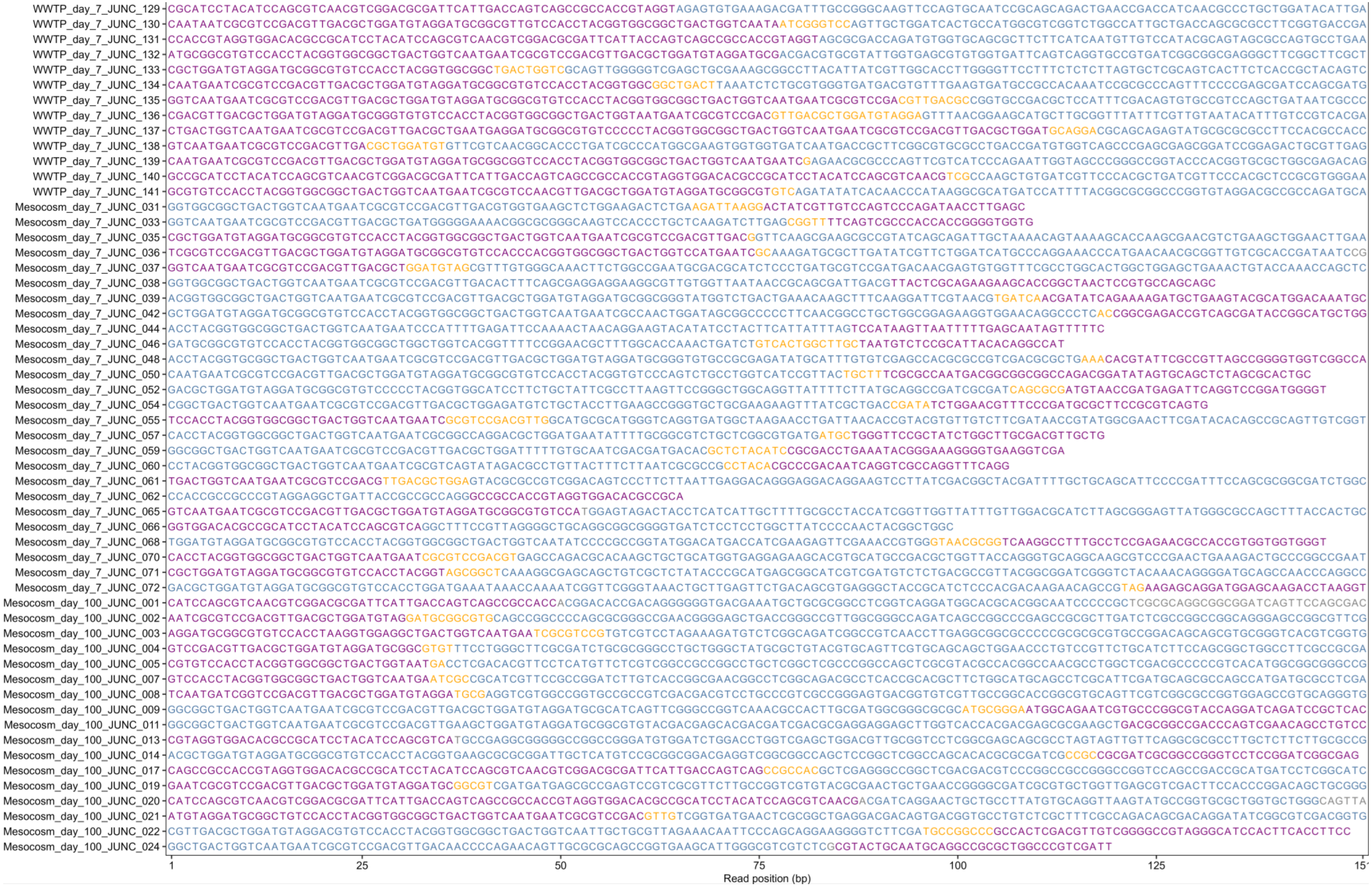
WWTP_day_7_JUNC_129
WWTP_day_7_JUNC_130
WWTP_day_7_JUNC_131
WWTP_day_7_JUNC_132
WWTP_day_7_JUNC_133
WWTP_day_7_JUNC_134
WWTP_day_7_JUNC_135
WWTP_day_7_JUNC_136
WWTP_day_7_JUNC_137
WWTP_day_7_JUNC_138
WWTP_day_7_JUNC_139
WWTP_day_7_JUNC_140
WWTP_day_7_JUNC_141
Mesocosm_day_7_JUNC_031
Mesocosm_day_7_JUNC_033
Mesocosm_day_7_JUNC_035
Mesocosm_day_7_JUNC_036
Mesocosm_day_7_JUNC_037
Mesocosm_day_7_JUNC_038
Mesocosm_day_7_JUNC_039
Mesocosm_day_7_JUNC_042
Mesocosm_day_7_JUNC_044
Mesocosm_day_7_JUNC_046
Mesocosm_day_7_JUNC_048
Mesocosm_day_7_JUNC_050
Mesocosm_day_7_JUNC_052
Mesocosm_day_7_JUNC_054
Mesocosm_day_7_JUNC_055
Mesocosm_day_7_JUNC_057
Mesocosm_day_7_JUNC_059
Mesocosm_day_7_JUNC_060
Mesocosm_day_7_JUNC_061
Mesocosm_day_7_JUNC_062
Mesocosm_day_7_JUNC_065
Mesocosm_day_7_JUNC_066
Mesocosm_day_7_JUNC_068
Mesocosm_day_7_JUNC_070
Mesocosm_day_7_JUNC_071
Mesocosm_day_7_JUNC_072
Mesocosm_day_100_JUNC_001
Mesocosm_day_100_JUNC_002
Mesocosm_day_100_JUNC_003
Mesocosm_day_100_JUNC_004
Mesocosm_day_100_JUNC_005
Mesocosm_day_100_JUNC_007
Mesocosm_day_100_JUNC_008
Mesocosm_day_100_JUNC_009
Mesocosm_day_100_JUNC_011
Mesocosm_day_100_JUNC_013
Mesocosm_day_100_JUNC_014
Mesocosm_day_100_JUNC_017
Mesocosm_day_100_JUNC_019
Mesocosm_day_100_JUNC_020
Mesocosm_day_100_JUNC_021
Mesocosm_day_100_JUNC_022
Mesocosm_day_100_JUNC_024
1
25
50
75
100
125
Read position (bp)
